# Balancing selection maintains hyper-divergent haplotypes in *C. elegans*

**DOI:** 10.1101/2020.07.23.218420

**Authors:** Daehan Lee, Stefan Zdraljevic, Lewis Stevens, Ye Wang, Robyn E. Tanny, Timothy A. Crombie, Daniel E. Cook, Amy K. Webster, Rojin Chirakar, L. Ryan Baugh, Mark G. Sterken, Christian Braendle, Marie-Anne Félix, Matthew V. Rockman, Erik C. Andersen

## Abstract

Across diverse taxa, selfing species have evolved independently from outcrossing species thousands of times. The transition from outcrossing to selfing significantly decreases the effective population size, effective recombination rate, and heterozygosity within a species. These changes lead to a reduction in genetic diversity, and therefore adaptive potential, by intensifying the effects of random genetic drift and linked selection. Within the nematode genus *Caenorhabditis*, selfing has evolved at least three times and all three species, including in the model organism *Caenorhabditis elegans*, show substantially reduced genetic diversity relative to outcrossing species. Selfing and outcrossing *Caenorhabditis* species are often found in the same niches, but we still do not know how selfing species with limited genetic diversity can adapt to these environments. Here, we examine the whole-genome sequences from 609 wild *C. elegans* strains isolated worldwide and show that genetic variation is concentrated in punctuated hyper-divergent regions that cover 20% of the *C. elegans* reference genome. These regions are enriched in environmental response genes that mediate sensory perception, pathogen response, and xenobiotic stress response. Population genomic evidence suggests that genetic diversity in these regions has been maintained by long-term balancing selection. Using long-read genome assemblies for 15 wild strains, we show that hyper-divergent haplotypes contain unique sets of genes and show levels of divergence comparable to levels found between *Caenorhabditis* species that diverged millions of years ago. These results provide an example for how species can avoid the evolutionary “dead end” associated with selfing.

## Introduction

Across the tree of life, selfing species have evolved independently from outcrossing species thousands of times^1, 2^. This reproductive mode transition has profound effects on the evolutionary processes that shape the genome. For example, selfing reduces the effective population size by a factor of two^3^ and the efficacy of recombination, leading to stronger effects of linked selection (selective sweeps of beneficial mutations or background selection against deleterious mutations)^4, 5^. Furthermore, because a single selfing individual can seed an entire population, founder effects are expected to be more severe^6, 7^. Together, these factors lead to a substantial reduction in the genetic diversity in selfing species^8^. This reduced diversity is expected to limit the ability of these lineages to adapt to new environments and could ultimately lead to their extinction, known as the “evolutionary dead end” hypothesis, potentially explaining why most selfing lineages are evolutionarily young^9^.

Outcrossing nematode species have some of the highest levels of diversity across all eukaryotes, but the species that predominantly reproduce by self-fertilization have at least an order of magnitude less variation than their outcrossing relatives^10–12^. In the nematode genus *Caenorhabditis*, self-fertilization has independently evolved at least three times, including in the model organism *Caenorhabditis elegans*^13^. Although *C. elegans* is capable of outcrossing, genetic evidence suggests that the selfing rate of this species is 99%^14–16^. As expected in species with high selfing rates, linked selection strongly influences genome-wide patterns of genetic diversity. Four out of the six *C. elegans* chromosomes show evidence of large-scale selective sweeps that have reduced diversity across large portions of the worldwide population^10^. Despite limited genetic diversity, *C. elegans* has colonized diverse environmental niches throughout the world^17^. Perhaps counterintuitively, strains whose genomes have been subject to the selective sweeps are found much more frequently and exhibit a wider ecological range than strains that have avoided the selective sweeps^10, 18^. This observation and evidence that the selective sweeps likely occurred within the last 200-300 years led to the hypothesis that the swept haplotype might harbor adaptive alleles that are favorable in human-associated habitats^10^. This hypothesis is further supported by the observation that divergent strains are typically isolated from natural habitats that are more isolated from human activity^18^.

Balancing selection can maintain adaptive genetic variation for long periods of time against evolutionary forces that constantly reduce genetic diversity (*e.g*. genetic drift, background selection). Though genome-wide scans for signatures of long-term balancing selection often have limited abilities to detect balanced loci, a handful of examples have been identified, predominantly in selfing species^19–22^. For example, recent scans in the *Capsella* genus have identified polymorphisms in immune-response loci that have been maintained since the divergence of selfing and outcrossing species^23^. Additionally, long-term balancing selection in immune-response genes have also been found in flies^24^ and humans^25^. Long-term balancing selection increases the coalescent times of neutral sites that are tightly linked to the selected alleles^26^, causing balanced haplotypes to accumulat^e^ higher-than-average levels of variation over time. It is hypothesized that the evolutionary forces that reduce genetic variation in selfing species have the potential to increase this genomic footprint of loci under balancing selection^27, 28^. Intriguingly, genomic loci under long-term balancing selection have also been characterized in *C. elegans*^29–31^, and previous comparisons between the genome of the laboratory reference strain N2 and a *de novo* genome assembly of a divergent Hawaiian strain (CB4856) led to the discovery of punctuated regions of extremely high divergence^32, 33^.

Here, by examining the whole-genome sequences of 609 wild *C. elegans* strains isolated across the world, we discovered previously uncharacterized levels and patterns of genetic diversity in this species. Our analysis of short- and long-read sequence data, led to the identification of 366 distinct hyper-divergent regions that span approximately 20% of the *C. elegans* reference genome. These regions are enriched for genes that mediate environmental responses, contain genes that are not present in the reference N2 genome, and often shared among many wild strains, suggesting that genetic diversity in hyper-divergent regions might be maintained to enable the species to thrive in diverse environments. Furthermore, the hyper-divergent haplotypes within these regions have levels of divergence comparable to that found between *Caenorhabditis* species that diverged millions of years ago. Finally, by characterizing hyper-divergent regions of another selfing species, *Caenorhabditis briggsae*, we show that punctuated regions of hyper-divergence are likely a common feature in the genomes of selfing *Caenorhabditis* species.

## Results

### Global distribution of *C. elegans* genetic diversity

To explore the genetic diversity in the *C. elegans* species, we examined whole-genome sequence data from 609 wild *C. elegans* strains isolated from six continents and several oceanic islands, including 103 wild strains that have not been studied previously (Fig. 1a, Supplementary Table 1). We identified 328 distinct genome-wide genotypes (henceforth, referred to as isotypes) (Methods)^15, 34^. The majority of wild strains (368 strains) were classified into isotypes in which each strain in the isotype was sampled from the same location, which is consistent with previous observations that local habitats frequently consist of clonal populations^15, 34^. However, we discovered 14 isotypes that were sampled from locations at least 50 km apart (Supplementary Fig. 1a, Supplementary Table. 2), suggesting that individuals can migrate long distances in the wild. We used species-wide variant data (2,431,645 single nucleotide variants (SNVs) and 845,797 ≤ 50 bp insertion and deletion variants) to identify the most highly divergent isotypes, which were isolated exclusively from a Pacific region that encompasses the Hawaiian islands, New Zealand, and the Pacific coast of the United States (Fig. 1a,b, Supplementary Fig. 1b-e).

**Fig. 1.**
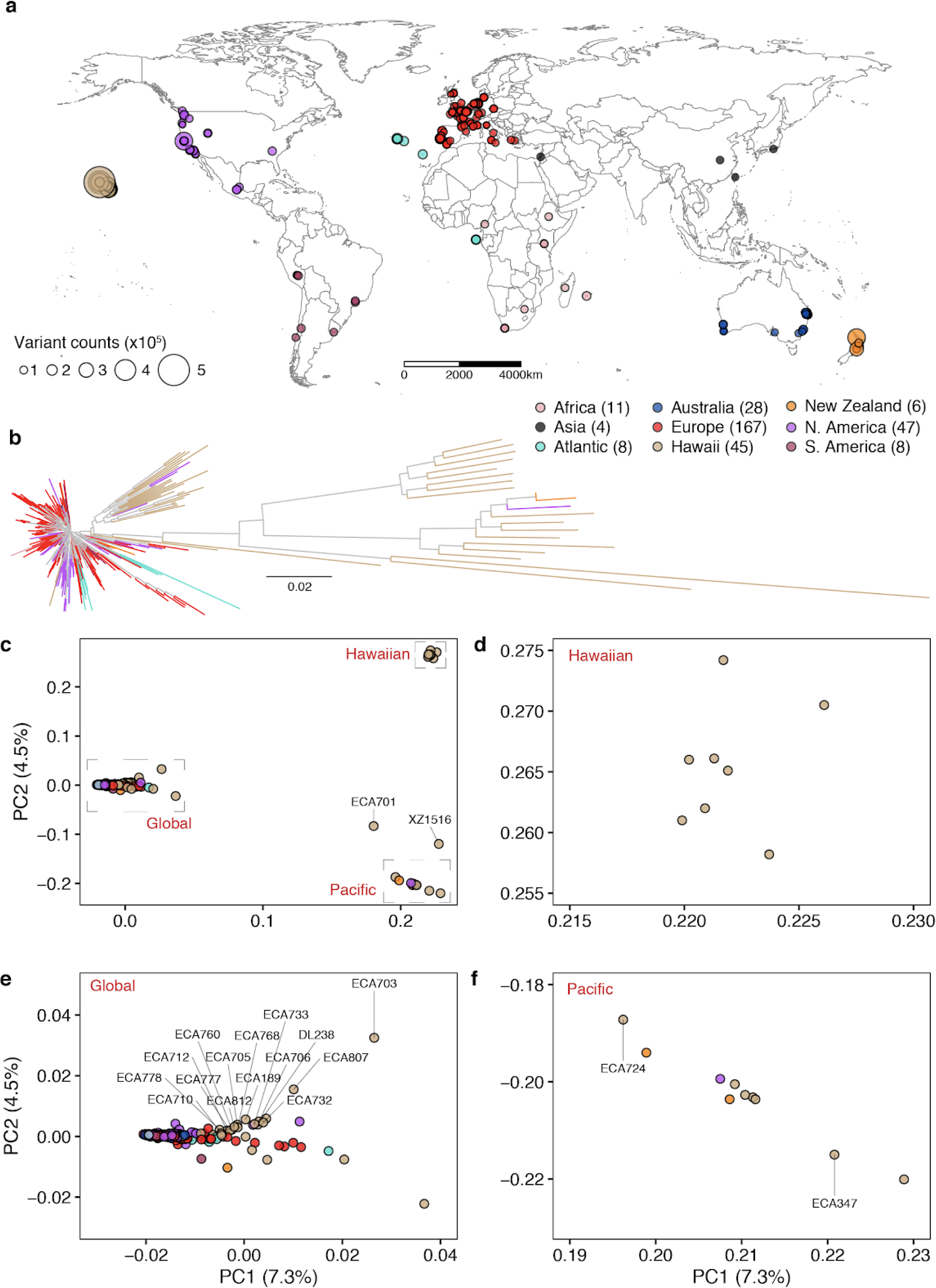
Genetically divergent wild *C. elegans* strains were isolated from the Pacific region. (a) The global distribution of 324 isotypes is shown. Each circle corresponds to one of the 324 isotypes and is colored by the geographic origin. Four isotypes that do not have geographic information are not shown. The size of each circle corresponds to the number of non-reference homozygous alleles. The number of isotypes from each geographic origin is labeled in parentheses. (b) A neighbour-joining tree of 328 *C. elegans* isotypes generated from 963,027 biallelic segregating sites is shown. The tips for four isotypes with unknown geographic origins are colored grey and the other 324 isotypes are colored the same as (a) by their geographic origins. (c) Plots of the 328 isotypes according to their values for each of the two significant axes of variation, as determined by principal component analysis (PCA) of the genotype covariances. Each point is one of the 328 isotypes, which is colored by their geographic origins same as (b). Two divergent isotypes, XZ1516 and ECA701, are labeled with grey lines. (d-f) Zoomed in plots of the Hawaiian, Global, and Pacific groups of (c), respectively. (c, e,f) Isotypes from the Big Island of Hawaii are labeled with grey lines. All isotypes in the Hawaiian group were isolated from the Big Island Hawaii.

To further characterize the geographic population structure within the *C. elegans* species, we performed principal component analysis (PCA)^35^ and found that most of the isotypes (326 isotypes) could be classified into three genetically distinct groups (Global, Hawaiian, and Pacific groups) (Fig. 1c-f, Supplementary Fig. 2). The largest Global group includes 93.9% (308) of all isotypes from six continents and oceanic islands (Fig. 1e). The Hawaiian group consists of eight isotypes from the Big Island of Hawaii (Fig. 1d), and the Pacific group includes ten isotypes from Hawaii, California, and New Zealand (Fig. 1f). This population structure is consistent with previous studies, where Hawaiian strains were shown to harbor higher genetic diversity than the rest of the worldwide population^18^. Hawaii is the only location where the isotypes of all three groups have been found. For example, the Big Island harbors 25 isotypes from all three groups, whereas all 167 European isotypes belong only to the Global group (Fig. 1d-f). Furthermore, the two most divergent isotypes, XZ1516 and ECA701, which do not belong to any of the three groups, were sampled from Kauai, the oldest sampled Hawaiian island (Fig. 1c). The remarkable genetic diversity sampled from the Hawaiian Islands suggests that *C. elegans* could have originated from the Pacific region^10, 18^. The current geographic location with the most divergent strains might not reflect the origin of the species because worldwide sampling is uneven and some sites, including Asia, have not been sampled extensively. Therefore, *C. elegans* could have originated elsewhere and spread throughout the Pacific region.

Next, we attempted to further subdivide the Global group using PCA. This group includes all non-Pacific isotypes and the majority of isotypes from the Pacific Rim. Although we found weak genetic differentiation of Global isotypes isolated from Hawaii, North America, and Atlantic locations from the rest of Global isotypes, this group was not further clustered into distinct genetic groups (Supplementary Fig. 2,3). Additionally, the Global group could not be separated into distinct geographic locations because many genetically similar isotypes have been sampled from different continents. By characterizing genomic regions predicted to be identical by descent, we found that the previously reported chromosome-scale selective sweeps^10^ contribute substantially to the genetic similarity we observed among geographically distant isotypes (Extended Data Fig. 1, Supplementary Fig. 3). For example, a large haplotype block on the center of chromosome V is shared by isotypes from six continents (Africa, Asia, Australia, Europe, North America, and South America) (Extended Data Fig. 1). In addition to the Hawaiian isotypes that were reported to have avoided these selective sweeps^18^, we found that the genomes of isotypes from Atlantic islands (*e.g.* Azores, Madeira, and São Tomé), in contrast to continental isotypes, show less evidence of the globally distributed haplotype that swept through the species (Extended Data Fig. 1, Supplementary Table 3).

### Discovery of species-wide hyper-divergent genomic regions

Previous studies have shown that genetic variation is distributed non-randomly across each of the five *C. elegans* autosomes, with the chromosome arms harboring more genetic variation than chromosome centers^10^. This distribution is shaped by higher recombination rates on chromosome arms than centers, which influences the effects of background selection^36–38^. Consistent with previous studies^10^, we observed 2.2-fold higher levels of nucleotide diversity (π) on chromosome arms compared to the centers (Welch’s t-test, *P* < 2.2 × 10^-16^) (Extended Data Fig. 2). Furthermore, we found that variation is concentrated in punctuated regions of extremely high diversity that are marked by a higher-than-average density of small variants and large genomic spans where short sequence reads fail to align to the N2 reference genome (Fig. 2a, Supplementary Fig. 4).

**Fig. 2.**
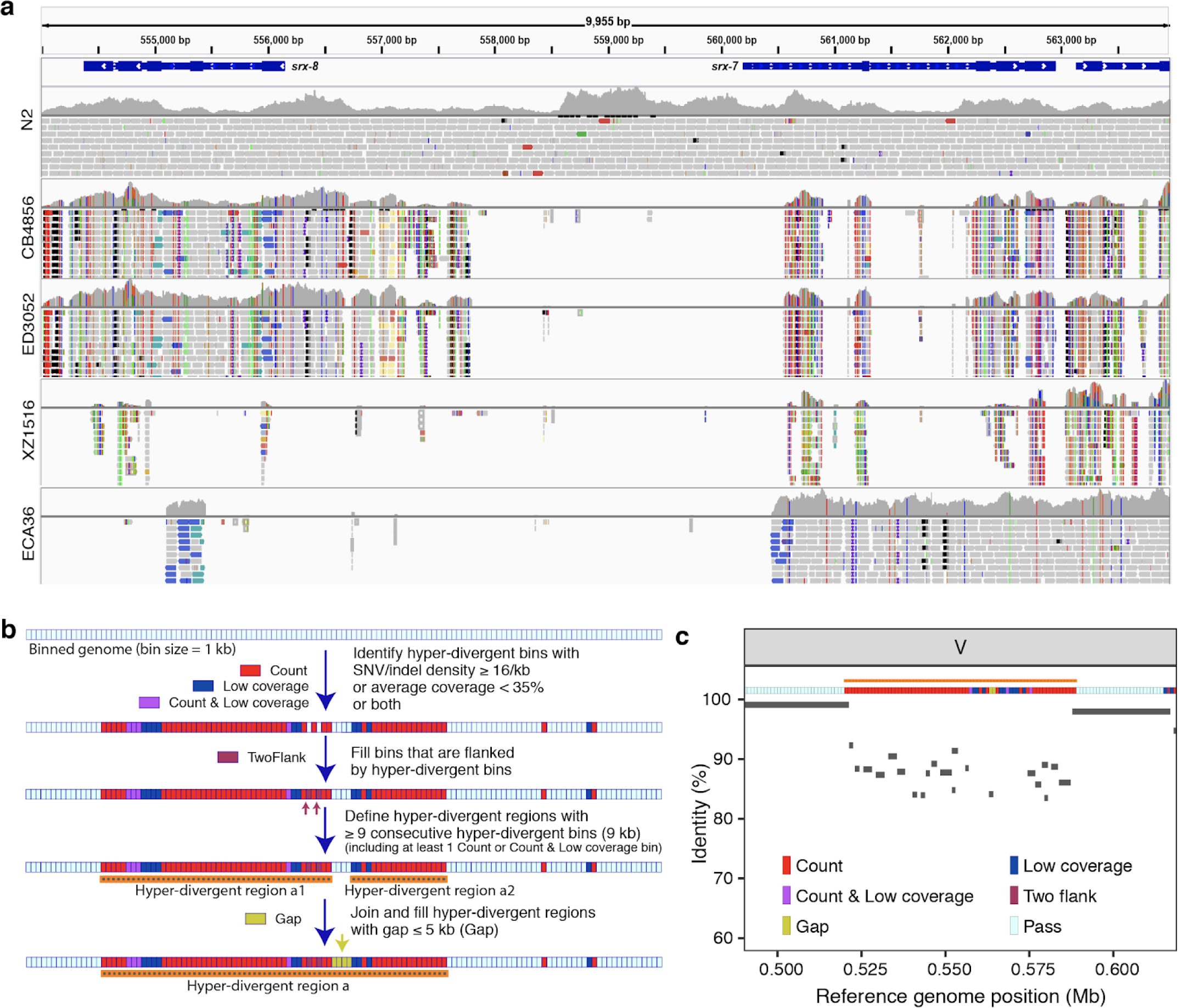
Characterization of hyper-divergent regions at the isotype level. (a) Short-read alignments from five isotypes (N2, CB4856, ED3052, XZ1516, and ECA36) to the N2 reference genome (WS245) for a region of chromosome V (V:554,000-564,000) are shown. Genes (*srx-7* and *srx-8*) in the interval are shown at the top. For each isotype, the top panel shows the coverage of genomic positions and the bottom panel shows aligned short-reads at genomic positions (gray: normal reads, red: reads with putative deletion, blue: reads with putative insertion, navy and turquoise: reads with putative inversion, green: reads with putative duplication or translocation). Colored vertical lines indicate mismatched bases at the position. (b) The workflow for the characterization of hyper-divergent regions at the isotype level is shown (See Methods). (c) An example plot showing the short-read based characterization of a hyper-divergent region and the percent sequence identity from a long-read alignment. The orange horizontal line shows a hyper-divergent region (V:520,000-589,000) in the locus (V:490,000-619,000) of CB4856 classified using this approach. The multi-colored horizontal rectangle shows classifications of genomic bins in the locus. Gray bars correspond to the alignments from CB4856 long-read sequences to the N2 reference genome (WS245), and identities of alignments are shown on the y-axis.

We sought to characterize the species- and genome-wide distributions of these regions (henceforth, referred to as hyper-divergent regions). To facilitate the accurate identification of these hyper-divergent regions across the *C. elegans* species, we generated high-quality genome assemblies using long-read sequencing data for 14 isotypes that span the species-wide diversity (Methods, Supplementary Table 4). We first aligned these assemblies, along with a previously published long-read assembly of the Hawaiian isotype CB4856^33^, to the N2 reference genome. Next, we used these alignments to classify hyper-divergent regions across a range of parameter values (*e.g*. alignment coverage and sequence divergence). We performed a similar parameter-search procedure using short-read alignments of these 15 isotypes and identified a set of short-read parameters that maximized the overlap between long- and short-read hyper-divergent classification (Methods, Extended Data Fig. 3). This analysis identified the optimal parameters to classify hyper-divergent regions as at least nine consecutive 1 kb bins that have one or both of the following criteria: (1) more than or equal to 16 SNVs/indels or (2) lower than 35% read depth to the genome-wide average (Fig. 2b, c). To validate our approach, we compared the hyper-divergent regions we identified in the CB4856 isotype to what was previously reported in this isotype (2.8 Mb)^32^ and found that we detected a similar number and extent of these regions (3.2 Mb). Finally, we applied these optimized short-read hyper-divergent region classification parameters to the entire set of 327 non-reference isotypes.

**Fig. 3.**
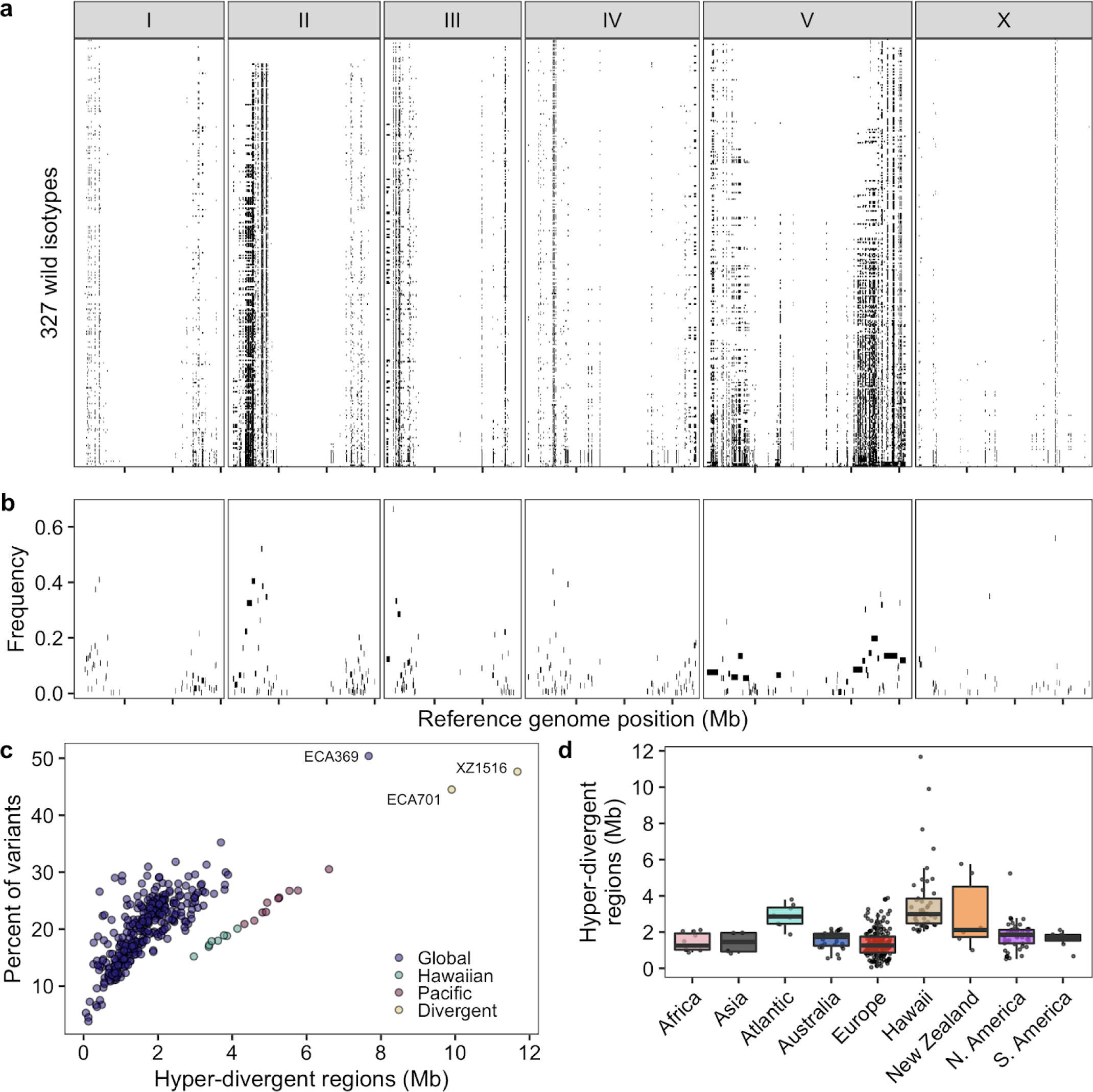
Punctuated hyper-divergent genomic regions are widespread across the *C. elegans* species. (a) The genome-wide distribution of hyper-divergent regions across 327 non-reference wild *C. elegans* isotypes is shown. Each row represents one of the 327 isotypes, ordered by the total amount of genome covered by hyper-divergent regions (black). The genomic position using the N2 reference in Mb is plotted on the x-axis, and each tick represents 5 Mb of the chromosome. (b) The species-wide frequencies of hyper-divergent calls for 366 species-wide hyper-divergent regions (Methods) are shown. Each rectangle corresponds to a block of species-wide hyper-divergent regions. The reference genomic position in Mb is plotted on the x-axis, and each tick represents 5 Mb of the chromosome. The average frequencies of hyper-divergent calls across 1 kb bins in each block across 327 non-reference isotypes are shown on the y-axis. (c) A scatter plot of the extent of the N2 reference genome that is hyper-divergent in each isotype (x-axis) and the fraction of total variants in hyper-divergent regions of 327 non-reference isotypes (y-axis) is shown. Each point corresponds to one of the 327 isotypes and is colored by the genetic grouping (Fig. 1c). The names of three isotypes with the largest genome-wide extents of hyper-divergent regions are shown. (d) Tukey box plots of the total amount of genome covered by hyper-divergent regions are shown with data points plotted behind. Isotypes are grouped by their geographic origins. Each point corresponds to one of the 324 isotypes with a known geographic origin, and the extent of the N2 reference genome that is hyper-divergent in each isotype is shown on the y-axis. The horizontal line in the middle of the box is the median, and the box denotes the 25th to 75th quantiles of the data. The vertical line represents the 1.5x interquartile range.

Across all isotypes, we identified 366 non-overlapping genomic regions that are hyper-divergent from the reference isotype N2 in at least one isotype (Fig. 3a, b, Supplementary Table 5, Supplementary Data 1). These regions range in size from 9 kb to 1.35 Mb (mean = 56 kb, median = 19 kb) (Supplementary Fig. 5) and cover approximately 20% of the N2 reference genome (20.5 Mb). The majority of these regions (69%) are found on autosomal arms and have 10.3-fold higher variant (SNVs/indels) densities than the non-divergent autosomal arm regions (16.6-fold more than the genome-wide average) (Supplementary Table 6, Extended Data Fig. 4). Although variant counts are likely underestimated because short sequencing reads often fail to align to the reference genome in hyper-divergent regions, we note that the level of genetic diversity in these regions is similar to the level of genetic diversity reported in outcrossing *Caenorhabditis* species^39, 40^. Across all non-reference isotypes, we found substantial differences in the extent of the genome that was classified as hyper-divergent (0.06 - 11.6% of the genome) (Fig. 3c,d). The genome of the most divergent isotype (XZ1516) contains a striking 11.7 Mb of hyper-divergent regions. In general, smaller fractions of the genomes of Global group isotypes were classified as hyper-divergent, which reflects that these isotypes are more closely related to the reference N2 isotype (Fig. 3c). A notable exception to this trend is the Global group isotypes that were isolated from Atlantic islands, which have, on average, a larger extent of their genome classified as hyper-divergent than the rest of the Global group (Fig. 3d). On average, 20.2% of the variation in a typical isotype is found in hyper-divergent regions that span 1.9% of the reference genome (Fig. 3c, Supplementary Table 3).

**Fig. 4.**
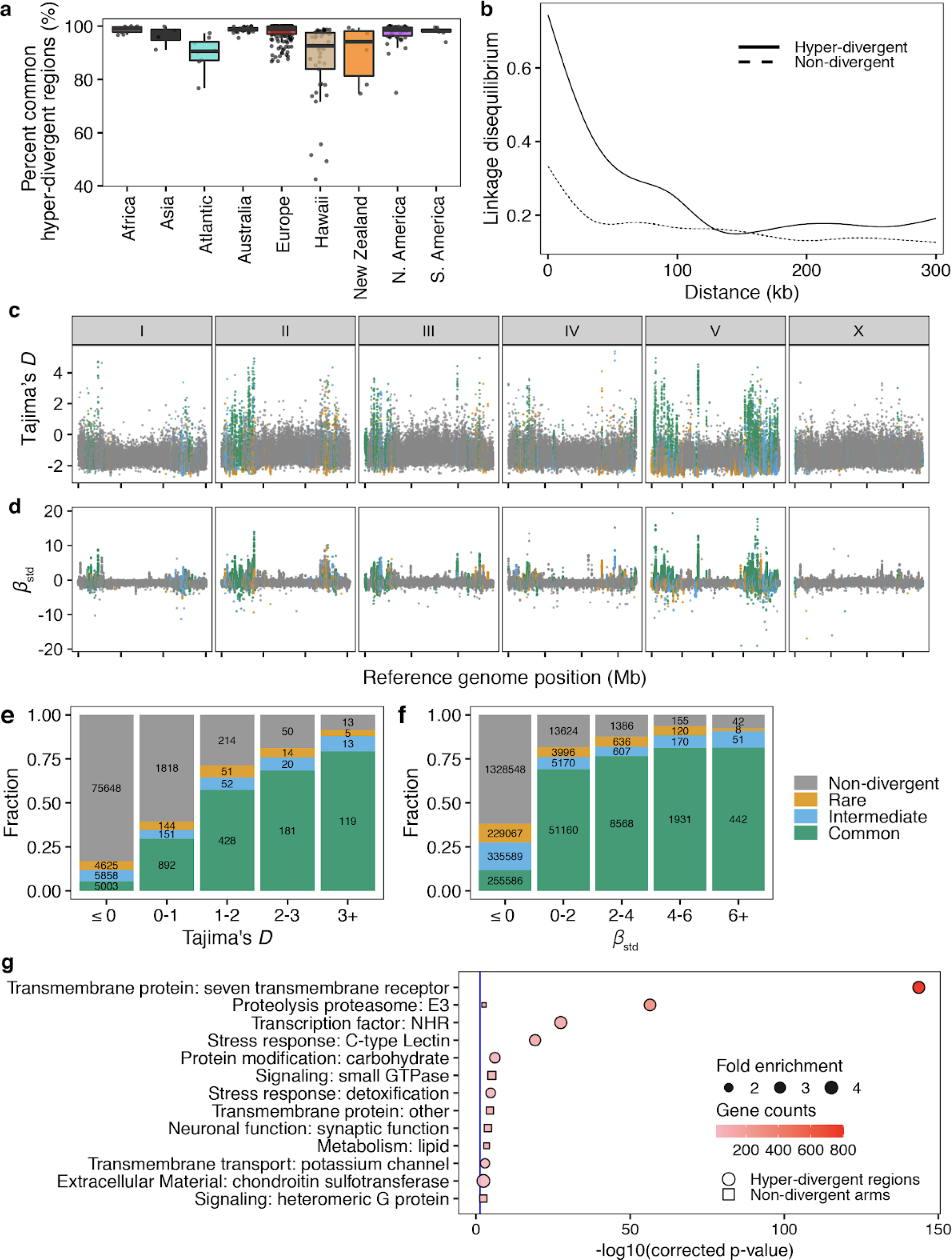
Balancing selection has maintained hyper-divergent haplotypes enriched in environmental-response genes. (a) Tukey box plots of the total amount of genome covered by common hyper-divergent regions are shown with data points plotted behind. Isotypes are grouped by their geographic origin. Each point corresponds to one of the 327 non-reference isotypes. The horizontal line in the middle of the box is the median, and the box denotes the 25th to 75th quantiles. The vertical line represents the 1.5x interquartile range. (b) Linkage disequilibrium (LD) decay in non-divergent and hyper-divergent regions in autosomal arms is shown. Note that 99.9% confidence intervals are represented around each smoothed line (generalized additive model fitting) as grey bands, but not visible because of the narrow range of the intervals. Distances between SNVs are shown on the x-axis, and LD statistics (*r*^2^) are shown on the y-axis. (c) The genome-wide pattern of Tajima’s *D* is shown. Each point represents a non-overlapping 1 kb genomic region and is colored by its divergent classification, (non-divergent region (gray), rare (<1%, orange), intermediate (≥1% and <5%, blue), or common (≥ 5%, green) hyper-divergent region). (d) The genome-wide pattern of standardized beta statistics is shown. Each dot corresponds to the value for a particular variant, colored by its location, as (c). (c,d) The reference genome position in Mb is plotted on the x-axis, and each tick represents 5 Mb of the chromosome. (e,f) Stacked bar plots for fractions of genomic bins (1 kb) for Tajima’s *D* (e) or fractions of variants for standardized beta (f) are shown. Genomic bins and variants are grouped and colored by their location same as (c). Numbers in bar plots represent counts of genomic bins (e) or variants (f). (g) Gene-set enrichment for autosomal arm regions (square) and hyper-divergent regions (circle) are shown. Bonferroni-corrected significance values for gene-set enrichment are shown on the x-axis. Sizes of squares and circles correspond to the fold enrichment of the annotation, and colors of square and circle correspond to the gene counts of the annotation. The blue line shows the Bonferroni-corrected significance threshold (corrected *p*-value = 0.05).

### Maintenance of hyper-divergent haplotypes

The species-wide distribution of hyper-divergent regions revealed that many non-reference isotypes are often hyper-divergent at the same regions of the N2 reference genome (Fig. 3a,b, Fig. 4a). We found that these regions range from being divergent in a single isotype to divergent in 280 isotypes (85%). Interestingly, we find that SNVs in hyper-divergent regions have a lower rate of linkage disequilibrium (LD) decay than SNVs within non-divergent regions on the autosomal arms (Fig. 4b), suggesting that these regions are inherited as large haplotype blocks. We performed genome-wide scans using commonly used statistics (Tajima’s *D* and standardized *β*) to identify regions under long-term balancing selection^21, 23, 30^, which could explain the presence of hyper-divergent haplotype blocks that are shared by a substantial fraction of isotypes^26^. We found that signatures of long-term balancing selection are concentrated in genomic regions that are frequently classified as hyper-divergent across the species (Fig. 4c-f). We note that estimates for Tajima’s *D* and *β* are likely biased toward lower values in hyper-divergent regions because short reads often fail to align to in these regions (Extended Data Fig. 5), which causes genetic diversity to be underestimated. Together, these results suggest that hyper-divergent haplotypes, which are frequently shared among many isotypes, have been maintained in the *C. elegans* species by long-term balancing selection.

**Fig. 5.**
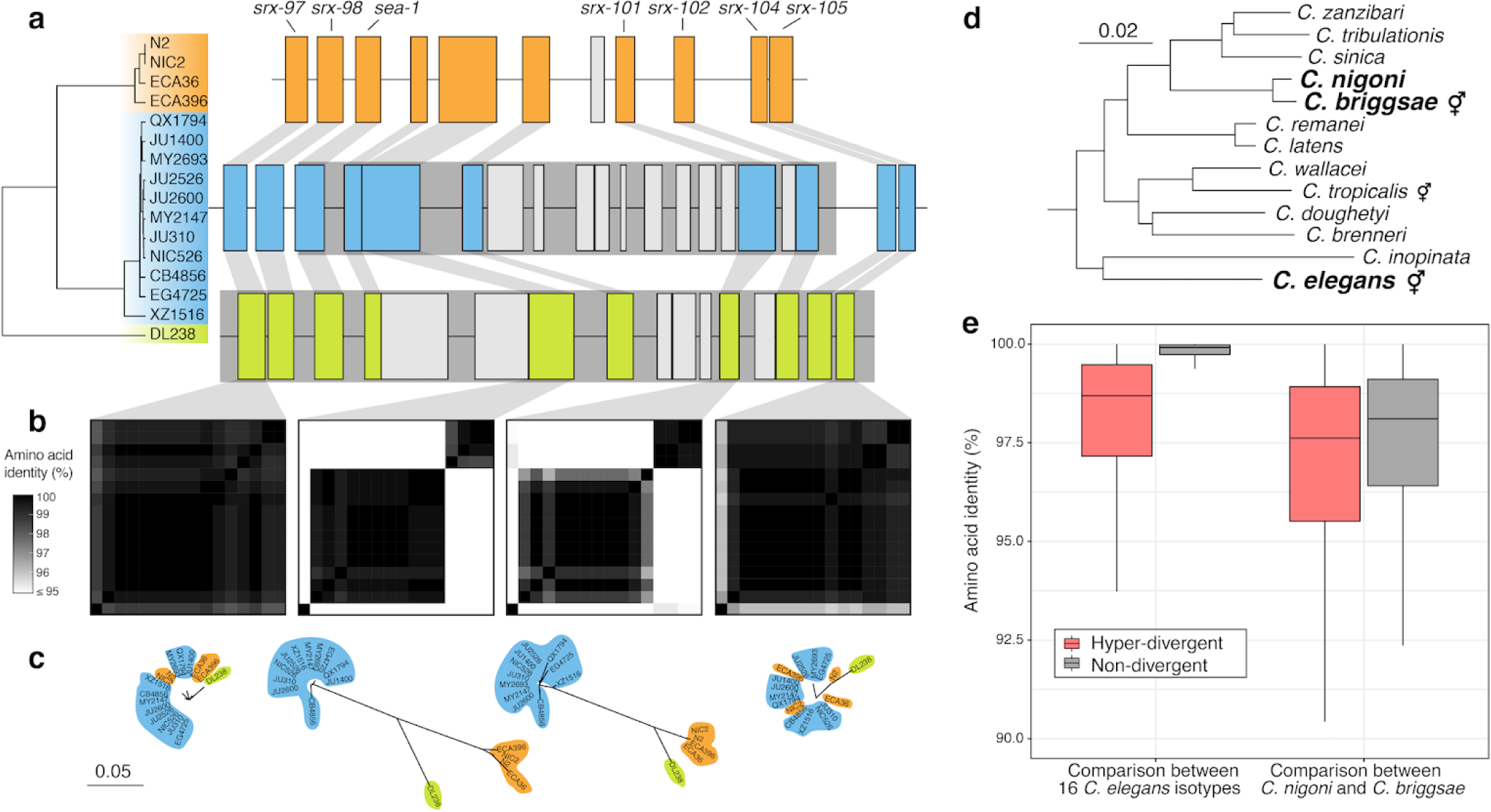
Hyper-divergent haplotypes contain ancient genetic diversity. (a) The protein-coding gene contents of three hyper-divergent haplotypes in a region on the left arm of chromosome II (II:3,667,179-3,701,405 of the N2 reference genome). The tree was inferred using SNVs and coloured by the inferred haplotypes. For each distinct haplotype, we chose a single isotype as a haplotype representative (orange haplotype: N2, blue haplotype: CB4856, green haplotype: DL238) and predicted protein-coding genes using both protein-based alignments and *ab initio* approaches (see Methods). Protein-coding genes are shown as boxes; those genes that are conserved in all haplotypes are coloured based on their haplotype, and those genes that are not are coloured light gray. Dark grey boxes behind loci indicate the coordinates of the hyper-divergent regions. Genes with locus names in N2 are highlighted. (b) Heatmaps showing amino acid identity for alleles of four loci (*srx-97*, *F19B10.10*, *srx-101*, and *srx-105*). Percentage identity was calculated using alignments of proteins sequences from all 16 isotypes. Heatmaps are ordered by the SNV tree shown in (a). (c) Maximum-likelihood gene trees of four loci (*srx-97*, *F19B10.10*, *srx-101*, and *srx-105*) inferred using amino acid alignments. Trees are plotted on the same scale (scale shown; scale is in amino acid substitutions per site). Isotype names are coloured by their haplotype. (d) *Caenorhabditis* phylogeny showing relationships within the *Elegans* subgroup^53^. The positions of *C. elegans*, *C. briggsae*, and *C. nigoni* are highlighted. Species that reproduce via self-fertilisation are indicated. Scale is in amino acid substitutions per site. (e) Tukey box plots showing amino acid identity of 8,741 genes that are single-copy and present in all 16 *C. elegans* isotypes, *C. briggsae*, and *C. nigoni*. Identities of alleles and their orthologs are shown separately for hyper-divergent and non-divergent regions.

Balancing selection can maintain genetic diversity that contributes to the adaptive potential of a population in the presence of environmental heterogeneity^41^. To investigate if hyper-divergent regions are functionally enriched for genes that enable *C. elegans* to thrive in diverse habitats, we performed gene-set enrichment analysis using two complementary approaches (Methods, Supplementary Data 2,3). Sensory perception and xenobiotic stress response were among the most significantly enriched gene classes (Fig. 4g, Extended Data Fig. 6, Supplementary Fig. 6). For example, we found that 54.9% (802/1461) of seven-transmembrane receptor class genes (*e.g.* G-protein coupled receptors, GPCRs) are located in hyper-divergent regions. In addition to GPCRs, 48.2% (124/257) of genes that encode for C-type lectins, 53% (317/598) of the E3 ligase genes, and 86.8% (33/38) of the *pals* genes, which are involved in response to diverse pathogens^31, 42–45^, are found in hyper-divergent regions (Supplementary Fig. 7, Supplementary Data 2). In agreement with these enrichment results, we found that 66.8% (131/196) and 65.4% (85/130) of genes that are differentially expressed in the reference N2 isotype upon exposure to the natural pathogens *Nematocida parisii*^46^ or Orsay virus^47^, respectively, are located in these regions^48^ (Supplementary Fig. 8). Furthermore, we found that genes in hyper-divergent regions are more strongly induced than genes in non-divergent regions of the genome (Welch’s t-test, *p*-value = 0.00606, for *N. parisii* and *p*-value = 0.0002226 for Orsay virus, respectively) (Supplementary Fig. 8). Notably, we found that hyper-divergent regions overlap with previously characterized genomic loci underlying natural variation in responses to the *C. elegans* pathogens *N. parisii*^49^ and the Orsay virus^50^, as well as responses to pathogens not found associated with *C. elegans* in nature^51^. These results suggest that high levels of variation in hyper-divergent regions likely facilitate diverse pathogen responses across the species.

### Hyper-divergent haplotypes contain potentially ancient genetic diversity

A common feature of the hyper-divergent regions is that short-read sequencing coverage is lower than the average genome-wide coverage (47% less coverage, on average), suggesting that divergence in these regions is too high for accurate alignment of short reads to the reference genome. Therefore, we took advantage of the 15 long-read genome assemblies to assess the content of these hyper-divergent regions. Strikingly, we found that these regions contain multiple hyper-divergent haplotypes that contain unique sets of genes and alleles that have substantially diverged at the amino acid level. For example, a hyper-divergent region on chromosome II (II:3,667,179-3,701,405 in the N2 reference genome) contains three distinct hyper-divergent haplotypes across the 16 isotypes for which we have high-quality assemblies (Fig. 5a). In this region, the reference isotype (N2) shares a haplotype with three isotypes and contains 11 protein-coding genes, including six GPCRs (*srx-97*, *srx-98*, *srx-101*, *srx-102*, *srx-104*, and *srx-105*). A second haplotype is shared among 11 isotypes and contains 20 protein-coding genes, including ten that are not conserved in the reference haplotype. The third haplotype is found only in a single isotype (DL238) and contains 17 protein-coding genes, including six that are not present in the reference haplotype. Of the 16 genes that are not conserved across the three haplotypes, 13 have clear homology to *srx-101*, *F40H7.12*, or *F19B10.10* (all found within this region in the reference haplotype) and three genes have homology to genes elsewhere in the genome, suggesting they have originated via duplication and diversification of existing genes, rather than *de novo* gene birth (Fig. 5a; Supplementary Data 4). For those genes that are conserved across all three haplotypes, alleles in different hyper-divergent haplotypes commonly show less than 95% amino acid identity (*e.g. F19B10.10* has an average between-haplotype identity of 88.3%, while *srx-97*, which lies outside the divergent region, has an average between-haplotype identity of 98.4%; Fig. 5b). The relationships inferred for this region using short-read SNV data were consistent with those relationships inferred using the long-read assemblies (Fig. 5a), allowing us to infer the haplotype composition of all non-reference isotypes for this region (Methods, Extended Data Fig. 7a). We found that a total of 59 isotypes contain the reference haplotype, 267 isotypes contain the second divergent haplotype, and one other isotype shares the DL238 haplotype.

In other regions of the genome, we found different numbers of hyper-divergent haplotypes across the 16 isotypes. For example, at the *peel-1 zeel-1* incompatibility locus on chromosome I (I:2,318,291-2,381,851 in the N2 reference genome), which was previously characterized to be maintained by long-term balancing selection^29^, we found two distinct hyper-divergent haplotypes across the 16 isotypes (Extended Data Fig. 8). The reference haplotype contains the *peel-1* and *zeel-1* toxin-antidote genes, which are absent in the alternative haplotype (Supplementary Fig. 8a). As expected under long-term balancing selection, genes that are linked to the *peel-1 zeel-1* locus show elevated sequence divergence. For example, the gene immediately adjacent to *zeel-1, ugt-31,* has an average between-haplotype identity of 95.7%, but *mcm-4,* which lies outside the divergent region, has an average between-haplotype identity of 99.9% (Extended Data Fig. 8b). In another region on the right arm of chromosome V (V:20,193,463-20,267,244 in the N2 reference genome) that contains three F-box genes (*fbxa-114*, *fbxa-113*, and *fbxb-59*), we found between six and seven hyper-divergent haplotypes, each with their own complement of genes (Extended Data Fig. 9; Supplementary Data 4). Strikingly, the six hyper-divergent alleles of the F-box gene *fbxb-59* all show less that 95% identity to each other (Extended Data Fig. 9b). Consistent with our hypothesis of long-term balancing selection, the sharing of haplotypes in all three regions presented here does not reflect the species-wide relationships (Fig. 5c, Extended Data Fig. 7, 8c, 9c). Instead, divergent isotypes from the Hawaiian and Pacific groups often share haplotypes with the Global group isotypes, suggesting that these haplotypes have been maintained since the last common ancestor of all sampled *C. elegans* isotypes.

To contextualize the high divergence we observe in hyper-divergent regions, we compared the level of divergence between hyper-divergent haplotypes to that observed between closely related *Caenorhabditis* species. As *C. elegans* lacks a closely related sister species (Fig. 5d), we compared the amino acid identities of *C. elegans* hyper-divergent alleles with the divergence between their orthologs in *Caenorhabditis briggsae* and *Caenorhabditis nigoni*, a closely related pair of species that are believed to have diverged from each other approximately 3.5 million years ago^52^. Notably, the average identity between hyper-divergent alleles within *C. elegans* is comparable to the divergence of their orthologs between *C. briggsae* and *C. nigoni* (mean identity in hyper-divergent regions of 97.7% and 96.4%, respectively, compared with a mean identity in non-divergent regions of 99.6% and 97.0%, respectively; Fig. 5e). Taken together, these results suggest hyper-divergent haplotypes might have diverged millions of years ago.

### Hyper-divergent regions are common features in the genomes of selfing *Caenorhabditis* species

Selfing has evolved at least three times independently in the *Caenorhabditis* genus and is the main reproductive mode for *C. elegans*, *C. briggsae*, and *C. tropicalis*^16, 54^ (Fig. 5d). We hypothesized that long-term balancing selection might have maintained punctuated hyper-divergent haplotypes in other selfing species. We used our short-read classification approach to identify hyper-divergent regions in the genomes of 35 wild *C. briggsae* strains^52^. In agreement with our hypothesis, we found that hyper-divergent regions are widespread in the genomes of wild *C. briggsae* strains (Extended Data Fig. 10, Methods). Moreover, we found that the same regions are divergent in strains from the ‘Tropical’ *C. briggsae* clade and divergent strains from other clades^52^. Therefore, it is likely that the same evolutionary processes that have maintained genetic diversity in *C. elegans* have also shaped the genome of *C. briggsae* and that hyper-divergent regions are common features in the genomes of selfing *Caenorhabditis* species.

## Discussion

Theory and empirical evidence show that predominantly selfing species have less genetic diversity than obligately outcrossing species^38^. It follows that reduced levels of genetic diversity in selfing species would limit the adaptive potential and range of ecological niches that the species can inhabit^55^. However, *C. elegans* and other selfing *Caenorhabditis* species are globally distributed, found in diverse habitats, and often share niches with outcrossing nematode species^16, 56^. We provide evidence that remarkable genetic diversity in hyper-divergent regions likely contributes to the distribution of *C. elegans* across diverse niches. We found that these genomic regions are significantly enriched for genes that modulate responses to bacteria, competitors, and pathogens in wild habitats^17^, including GPCRs, C-type lectins, and the *pals* gene family. Previous studies have functionally characterized hyper-divergent alleles of such genes within these regions that have profound impacts on *C. elegans* physiology and ecology. For example, distinct density-dependent foraging strategies have been shown to be mediated by hyper-divergent alleles of two neighboring genes (*srx-43* and *srx-44*) that encode for GPCRs^30, 57^. A quantitative trait locus on chromosome III that affects pheromone-induced developmental plasticity^58^ is another example of a hyper-divergent region associated with an ecologically relevant *C. elegans* phenotype. Furthermore, the enrichment of immune-response genes that we observed in hyper-divergent regions suggests that these regions might harbor variation that enables *C. elegans* to survive various pathogenic threats. In support of this hypothesis, previous studies have shown substantial genetic and phenotypic variation in response to the co-evolved fungal and viral pathogens, *N. parisii*^49^ and the Orsay virus^50^, respectively. Four genomic regions were originally identified to affect *C. elegans* susceptibility to *N. parisii*, all of which overlap with hyper-divergent regions we identified. Subsequent studies showed that the *pals* genes mediate responses to *N. parisii* and the Orsay virus^31, 42–45^, of which 86.8% (33/38) of these genes are located in hyper-divergent regions. The hyper-divergent regions we present here also include other loci previously found to underlie natural resistance to starvation^59^ and toxic bacterial metabolite^60^, and genetic incompatibilities in the species^29, 61^. However, the loci discussed here only represent a handful of the 366 distinct hyper-divergent regions we identified, suggesting that we still have much to learn about the role of these regions on *C. elegans* ecology and physiology.

The punctuated nature of the hyper-divergent haplotypes can be explained by the predominantly selfing lifestyle of *C. elegans*. Selfing leads to a reduction in the effective recombination rate and therefore increases LD, which increases the overall footprint of balancing selection^27, 28^. Furthermore, long-term sefling causes an overall reduction in genome-wide diversity^8^, making the high genetic diversity in balanced regions of the genome more pronounced. Given the outcrossing *Caenorhabditis* species also inhabit diverse niches as *C. elegans*, long-term balancing selection for environmental response genes could have occurred and shaped hyper-divergent haplotypes in those species as well. However, hyper-divergent haplotypes under long-term balancing selection in outcrossing species are expected to be more difficult to identify because they are likely to be smaller and less pronounced from the background of much higher genetic diversity.

Our findings highlight the limitations of short-read reference-based mapping approaches for characterizing variation in genetically diverse species. Using *de novo* genome assemblies generated from long-read data, we have shown that hyper-divergent haplotypes contain levels of genetic variation that are substantially underestimated using short-read based approaches, including genes that are not present in the reference genome and alleles that are highly diverged at the amino acid level. A similar observation was recently reported in soybean^62^, where thousands of dispensable genes and large structural variants were discovered using genome assemblies generated using long-read data. Exploiting these new sequencing technologies will be critical if we are to fully capture patterns of genetic variation in *C. elegans* and other species.

The origin of hyper-divergent haplotypes in *C. elegans* remains elusive. We hypothesize that ancient balancing selection and adaptive introgression could be two possible sources of hyper-divergent haplotypes. Hyper-divergent haplotypes could represent ancestral genetic diversity that has been maintained by balancing selection since the evolution of selfing in *C. elegans*, which is believed to have occured in the last four million years^63^. The amino acid divergence among *C. elegans* hyper-divergent haplotypes is similar to the divergence between *C. briggsae* and *C. nigoni*, two species that diverged approximately 3.5 million years ago^52^, suggesting that these haplotypes could have been maintained for millions of years. Moreover, outcrossing *Caenorhabditis* species typically have extremely high levels of genetic diversity^11, 12^. Assuming the outcrossing ancestor of *C. elegans* was similarly diverse, these hyper-divergent regions might represent the last remnants of ancestral genetic diversity that has otherwise been lost because of the long-term effects of selfing. Similar observations have been reported in the *Capsella* genus, where the selfing species, *Capsella rubella*, shares variants with its outcrossing sister species *Capsella grandiflora*^64^. These trans-specific polymorphisms are predominantly found at loci involved in immune response^23^, which match our findings in *C. elegans*. Alternatively, hyper-divergent haplotypes could have been introgressed from other species. Introgression was recently shown to explain the existence of large, divergent haplotypes that underlie ecotypic adaptation in sunflowers^65^ and characterized in other species^66^. Niche sharing of different *Caenorhabditis* species is often observed in natural habitats^16, 18, 56^, suggesting that hybridization with closely related species could have occurred in the *C. elegans* lineage. Because *C. elegans* and its closest known relative, *C. inopinata*, are estimated to have diverged 10.5 million years ago^67^, it is not possible to distinguish between retained ancestral polymorphism and relatively recent introgression by identifying trans-specific polymorphisms. However, the presence of up to seven *C. elegans* hyper-divergent haplotypes at a single locus suggests that introgression is unlikely to be an exclusive source of hyper-divergent haplotypes. Notably, we characterized a similar pattern of punctuated hyper-divergent regions in *C. briggsae*, which is a selfing species with a closely related outcrossing species *C. nigoni*. As similar population-wide variant data become available for *C. briggsae*, it might be possible to test if the patterns of divergence are consistent with introgression, retained ancestral genetic diversity, or both.

Regardless of their origin, the existence of these regions in *C. elegans* has important implications for how we understand the genetic and genomic consequences of selfing. It has been proposed that the evolution of selfing represents an evolutionary “dead end”, whereby the reduction in genetic diversity, and therefore adaptive potential, of a species will eventually lead to extinction^9^. However, our findings suggest that it is possible to avoid this fate by maintaining a substantial fraction of the ancestral or introgressed genetic diversity at key regions of the genome.

## Methods

### Strains

Nematodes were reared at 20°C using *Escherichia coli* bacteria (strain OP50) grown on modified nematode growth medium (NGMA)^68^, containing 1% agar and 0.7% agarose to prevent animals from burrowing. All 609 wild *C. elegans* strains are available on CeNDR^69^ and strain information can be found in the accompanying metadata (Supplementary Table 1).

### Sequencing and isotype characterization

#### Sequencing

To extract DNA, we transferred nematodes from two 10 cm NGMA plates spotted with OP50 into a 15 ml conical tube by washing with 10 mL of M9. We then used gravity to settle animals in a conical tube, removed the supernatant, and added 10 mL of fresh M9. We repeated this wash method three times over the course of one hour to serially dilute the *E. coli* in the M9 and allow the animals time to purge ingested *E. coli*. Genomic DNA was isolated from 100 to 300 µl nematode pellets using the Blood and Tissue DNA isolation kit (cat#69506, QIAGEN, Valencia, CA) following established protocols^70^. The DNA concentration was determined for each sample with the Qubit dsDNA Broad Range Assay Kit (cat#Q32850, Invitrogen, Carlsbad, CA). The DNA samples were then submitted to the Duke Sequencing and Genomic Technologies Shared Resource per their requirements. The Illumina library construction and sequencing were performed at Duke University using KAPA Hyper Prep kits (Kapa Biosystems, Wilmington, MA) and the Illumina NovaSeq 6000 platform (paired-end 150 bp reads). The raw sequencing reads for strains used in this project are available from the NCBI Sequence Read Archive (Project PRJNA549503).

#### Isotype characterization

Raw sequencing reads from 609 wild strains were trimmed using the Trimmomatic (v0.36)^71^ to remove low-quality bases and adapter sequences. Following trimming, we called SNVs using the BCFtools (v.1.9)^72^ and the following filters: Depth (DP) ≥ 3; Mapping Quality (MQ) > 40; Variant quality (QUAL) > 30; Allelic Depth (FORMAT/AD)/Num of high-quality bases (FORMAT/DP) ratio > 0.5. We classified two or more strains as the same isotype if they have the same call at 99.9% of all sites called across the full panel of wild strains. If a strain did not meet this criterion, we considered it as a unique isotype. Newly assigned isotypes were added to CeNDR^69^.

### Pacific Biosciences continuous long-read sequencing

To extract DNA, we transferred nematodes from twelve 10 cm NGMA plates spotted with OP50 into a 50 ml conical tube by washing with 30 mL of M9. We then used gravity to settle animals for 1.5 hours, removed the supernatant, and added 15 mL of fresh M9. We allowed nematodes to settle for 1 hour, removed the supernatant, and transferred nematodes to a microfuge tube using a Pasteur pettle. We settled animals for an additional 1 hour and any supernatant was removed. We stored pellets at −80°C prior to DNA extraction. Genomic DNA was isolated from 400 to 500 µl nematode pellets using the MagAttract HMW DNA kit (cat#67563, QIAGEN, Valencia, CA). The DNA concentration was determined for each sample with the Qubit dsDNA Broad Range Assay Kit (cat#Q32850, Invitrogen, Carlsbad, CA). The DNA samples were submitted to the Duke Sequencing and Genomic Technologies Shared Resource for library construction and sequencing using the Pacific Biosciences Sequel platform. Quality control was performed with the Qubit dsDNA Broad Range Assay Kit and Agilent Tapestation. Some samples had 30-50 kb fragment sizes, and a 15 kb cutoff was used for size selection of these samples during library preparation. A 20 kb cutoff was used for the other samples. The raw sequencing reads for strains used in this project are available from the NCBI Sequence Read Archive (Project PRJNA692613).

### Sequence alignments and variant calling

After isotypes are assigned, we used BWA (v0.7.17-r1188)^73, 74^ to align trimmed sequence data for distinct isotypes to the N2 reference genome (WS245)^75^. Variants were called using GATK4 (v4.1.0)^76^. First, gVCFs were generated for each isotype using the *HaplotypeCaller* function in GATK4 with the following parameters: *--max-genotype-count*=3000 and *--max-alternate-alleles*=100, using the WS245 N2 reference genome. Next, individual isotype gVCFs were merged using the *MergeVcfs* function in GATK4 and imported to a genomics database using the *GenomicsDBImport* function. Genotyping of the gVCFs was performed using the *GenotypeGVCFs* function in GATK4 with the following parameter *--use-new-qual-calculator*. The 328 isotype cohort VCF was annotated using SnpEff^77^ and an annotation database that was built with the WS261 gene annotations^77^. Following VCF annotation, we applied soft filters (QD < 5.0, SOR > 5.0, QUAL < 30.0, ReadPosRankSum < −5.0, FS > 50.0, DP < 5) to the VCF variants using the *VariantFiltration* function in GATK4. We applied a final isotype-specific soft filter called *dv_dp* using the bcftools *filter* function, which required the alternate allele depth to at least 50% of the total read depth for an individual isotype. All variant sites that failed to meet the variant-level filter criteria were removed from the soft-filtered VCF, and all isotype-level variants that did not meet the *dv_dp* criteria were set to missing. Finally, we removed sites that had more than 5% missing genotype data or more than 10% of samples were called heterozygous.

### Genetic relatedness

#### Similarity analysis

Using BCFtools (v.1.9), with the command filter -i N_MISSING=0, we filtered the high-quality VCF file and generated a VCF file (complete-site VCF) containing 963,027 biallelic SNVs that are genotyped for all 328 *C. elegans* isotypes. We used the vcf2phylip.py script^78^ to convert the complete-site VCF file to the PHYLIP format. The distance matrix and unrooted neighbor-joining tree were made from this PHYLIP file using *dist.ml* and *NJ* function using the phangorn (v2.5.5) R package^79^. The tree was visualized using the ggtree (version 1.16.6) R package^80^.

#### Principal component analysis

The *smartpca* executable from the EIGENSOFT (v6.1.4)^35, 81^ was used to perform principal component analysis. We performed analysis with the complete-site VCF with or without removing outlier isotypes to analyze the population structure with highly genetically divergent isotypes. When analyzing the population without removing outlier isotypes, we used the following parameters: altnormstyle: NO, numoutevec: 50, familynames: NO. When analyzing the population with outlier isotype removal, we set numoutlieriter to 15.

### Population genomic analyses

#### Haplotype analysis

We determined identity-by-descent (IBD) of genome segments using IBDSeq (version r1206)^82^ run on the complete-site VCF with the following parameters:minalleles = 0.01, r2window = 1500, ibdtrim = 0, r2max = 0.3 for genome-wide haplotype analysis. IBD segments were then used to infer haplotype structure among isotypes as described previously^10^. We defined the most common haplotype in chromosome I, IV, V, and X, of which the species-wide average fraction of the most common haplotype is greater than 30%, as the swept haplotype for each chromosome. We then retained the swept haplotypes within isotypes that passed the following per chromosome filters: total length >1 Mb; total length/maximum population-wide swept haplotype length >0.03.

#### Population genetics

We only considered bi-allelic SNVs to calculate population genomic statistics. Tajima’s *D*, Watterson’s theta, and Pi were all calculated using scikit-allel^83^. Each of these statistics was calculated for the same non-overlapping 1000 bp windows as hyper-divergent regions (described below). We calculated the standardized *β*^(1)84, 85^ using the BetaScan.py script by providing Watterson’s theta estimates with the -thetaMap flag. Default window parameters surrounding each marker were used (+/− 500 bp from the focal marker).

#### Linkage disequilibrium (LD) decay

We filtered the complete-site VCF file using BCFtools (v1.9) with the command view -q 0.05:minor and generated a VCF file (MAF-05 VCF) containing 123,830 SNVs of which minor allele frequencies are greater than or equal to 5%. Then, we selected 41,368 SNVs on autosomal arms (MAF-05-autoarm VCF) and split the MAF-05-autoarm VCF by the location of SNVs; MAF-05-autoarm-div VCF with 17,419 SNVs within hyper-divergent autosomal arm and MAF-05-autoarm-nondiv VCF with 23,949 SNVs within the non-divergent autosomal arms. We analyzed LD decay by running PopLDdecay (v3.31)^86^ on both MAF-05-autoarm-div VCF and MAF-05-autoarm-nondiv VCF with the default parameters (MaxDist=300, Het=0.88, Miss=0.25).

### Long-read genome assembly

We extracted PacBio reads in FASTQ format from the subread BAM files using the PacBio bam2fastx tool (version 1.3.0; available at https://github.com/PacificBiosciences/bam2fastx). For each of the 14 isotypes, we assembled the PacBio reads using three genome assemblers: wtdbg2 (version 0.0; using the parameters *-x sq -g 102m*)^87^, flye (version 2.7; using the parameters *--pacbio-raw -g 102m*)^88^, and Canu (version 1.9; using the parameters genomeSize=102m -pacbio-raw)^89^. For each assembly, we assessed the biological completeness using BUSCO^90^ (version 4.0.6; using the options *-l nematoda_odb10 -m genome*) and contiguity by calculating a range of numerical metrics using scaffold_stats.pl (available here: https://github.com/sujaikumar/assemblage). We selected the Canu assemblies as our final assemblies because they had both high contiguity and high biological completeness. To correct sequencing errors that remained in the Canu assemblies, we aligned short-read Illumina data for each isotype to the corresponding assembly using BWA-MEM^74^ (version 0.7.17) and provided the BAM files to Pilon for error correction (version 1.23; using the parameters *--changes --fix bases*)^91^. To remove assembled contigs that originated from non-target organisms (such as the *E. coli* food source), we screened each assembly using taxon-annotated GC-coverage plots as implemented in BlobTools^92^. Briefly, we aligned the PacBio reads to the assembly using minimap2^93^ (version 2.17; using the parameters *-a -x map-pb*) and sorted then indexed the BAM files using SAMtools^94^ (version 1.9). We searched each assembled contig against the NCBI nucleotide database (‘nt’) using BLASTN^95^ (version 2.9.0+; using the parameters *-max_target_seqs 1 -max_hsps 1 -evalue 1e-25*) and against UniProt Reference Proteomes^96^ using Diamond (version 0.9.17; using the parameters *blastx --max-target-seqs 1 --sensitive --evalue 1e-25*)^97^. We removed contigs that were annotated as being of bacterial origin and that had coverage and percent GC that differed from the target genome. Genome assembly metrics are shown in Supplementary Table 4.

### Characterization of hyper-divergent regions

To characterize hyper-divergent regions across the *C. elegans* species, we first analyzed short-read and long-read alignments of 15 isotypes. For all non-overlapping 1 kb windows in the reference *C. elegans* genome, we calculated the number of small variants (SNVs and indels) (variant count) using the coverage subroutine in the BEDtools (v2.27.1)^98^ suite and the average sequencing depth using mosdepth (v0.2.3)^99^. We converted the coverage values to coverage fraction (average sequencing depth of the window/genome-wide average depth). In parallel, we aligned all 14 long-read assemblies we generated along with a long-read assembly for the Hawaiian isotype CB4856^33^ to the N2 reference genome (WS255)^100^ using NUCmer (version 3.1) with the following parameters: *--maxgap=500 --mincluster=100*)^101^. Coordinates and identities of the aligned sequences were extracted from the alignment files using the ‘show-coords’ function with NUCmer. Then, we calculated the average alignment coverage (alignment coverage) and average alignment identity (alignment identity) for each non-overlapping window in the reference genome. Next, we used the long- and short-read alignment datasets to identify the optimal coverage fraction and variant count parameters to apply to the rest of the population for which we do not have long-read sequence data. We tested a wide range of parameters to define hyper-divergent regions from short-read and long-read alignments. For the short-read based approach, we classified each window as hyper-divergent if its variant count ≥ x or coverage fraction < y% or both; we also classified windows that are flanked by two hyper-divergent windows as hyper-divergent. Then we clustered contiguous hyper-divergent windows and defined clusters that are greater than or equal to 9 kb of N2 reference genome length as hyper-divergent regions^32^. For the long-read based approach, we classified each window as hyper-divergent if its alignment coverage < z% or alignment coverage < w% or both; we also classified windows that are flanked by hyper-divergent windows as hyper-divergent. Then, we clustered contiguous hyper-divergent windows and defined clusters that are greater than or equal to 9 kb of the N2 reference genome length as hyper-divergent regions. Because we lacked a “true” hyper-divergent region dataset to which we could tune our parameters, we identified the set of x, y, z, w values that maximized the overlap between hyper-divergent regions identified by short- and long-read based approaches (Extended Data Fig. 3). To minimize the amount of false positives that we detected, we manually validated hundreds of regions that were classified as hyper-divergent using IGV^102^. Using this optimization, we selected the optimal short-read classification parameters (variant counts ≥ 16 and coverage fraction < 35%), which we then applied to all 327 non-reference isotypes. With these classification parameters, we identified a similar size of hyper-divergent regions (3.2 Mb) in CB4856 to the total size of hyper-divergent regions (2.8 Mb) identified in CB4856 previously^32^. Additionally, we confirmed that selected parameters do not detect any hyper-divergent region from short-read alignments of N2 reference strain to its own reference genome. To exclude large deletions that could be classified as hyper-divergent regions, we filtered out hyper-divergent regions without any window with high variant density that exceed variant count threshold. To classify species-wide hyper-divergent regions, we combined continuous hyper-divergent regions that were identified in individual strains. To compare small variant (SNVs/indels) density between hyper-divergent and non-divergent regions for chromosomal centers, arms, and tips (Supplementary Table 6), we used previously defined genomic coordinates for centers, arms, and tips of six chromosomes^36^. For bins with no isotype classified as hyper-divergent, we measured the variant density as the average variant density of all 328 isotypes. For bins with at least one isotype classified as hyper-divergent, we measured the variant density as the average variant density of isotypes that are classified as hyper-divergent for each bin. We calculated the percent of 327 non-reference isotypes that is classified as hyper-divergent (percent divergence) for each 1 kb bin, and classified each bin into one of three frequency groups based on its percent divergence: rare < 1%, 1% ≤ intermediate < 5%, common ≥ 5%.

For *C. briggsae*, we performed a sliding window analysis with a 1 kb window size and a 1 kb step size for each of 35 non-reference wild *C. briggsae* genomes^52^. We only used variant counts to classify hyper-divergent regions. Because the *C. briggsae* genome-wide variant density is 1.55 greater than *C. elegans*, we used a modified variant count threshold (variant counts ≥ 24). We also classified windows that were flanked by hyper-divergent windows as hyper-divergent.

### Gene-set enrichment analysis

We analyzed the gene-set enrichment of hyper-divergent regions using web-based WormCat^103^ that contains a near-complete annotation of genes from the *C. elegans* reference strain N2. We also performed a conventional gene ontology (GO) enrichment analysis using the clusterProfiler (v3.12.0) R package^104^ and org.Ce.eg.db: Genome-wide annotation for Worm^105^. Because the majority of hyper-divergent regions are found on autosomal arms (Fig. 3a, Supplementary Table 6), we used gene-set enrichment on the autosomal arms as a control dataset^36^.

### Protein-coding gene prediction

To generate *ab initio* gene predictions for all 14 assemblies and for a previously published long-read assembly for the Hawaiian isotype CB4856^33^, we first used RepeatMasker^106^ (ver 4.0.9; using the parameter *-xsmall*) to identify and soft-mask repetitive elements in each genome assembly using a library of known Rhabditida repeats from RepBase^107^. We predicted genes in the masked assemblies using AUGUSTUS (version 3.3.3; using the parameters *--genemodel=partial --gff3=on --species=caenorhabditis --codingseq=on*)^108^. We extracted protein sequences and coding sequences from the GFF3 file using the getAnnoFasta.pl script from AUGUSTUS. We assessed the biological completeness of each predicted gene set using BUSCO (version 4.0.6; using the options *-l nematoda_odb10 -m proteins*) using the longest isoforms of each gene only. Predicted protein-coding gene counts and BUSCO completeness scores are shown in Supplementary Table 4.

### Characterising gene contents of hyper-divergent haplotypes

To characterise and compare the gene contents of hyper-divergent haplotypes, we used OrthoFinder^109^ (version 2.3.11; using the parameter *-og*) to cluster the longest isoform of each protein-coding gene predicted from the genomes of 15 wild isolates and the N2 reference genome (WS270). We initially attempted to use the orthology assignments to identify alleles in the 15 isotypes for each region of interest. However, gene prediction errors were pervasive, with many fused and split gene models, which caused incorrect orthology assignment and/or spurious alignments. To ensure the differences between hyper-divergent haplotypes were real biological differences and not gene prediction artefacts, we used the AUGUSTUS gene predictions and orthology assignments to identify coordinates of hyper-divergent haplotypes only. For a given region of interest, we identified genes in the reference genome that flanked the hyper-divergent regions and used the orthology assignments to identify the corresponding alleles in all 15 wild isolates. Using the coordinates of these genes, we extracted the sequences of intervening hyper-divergent haplotypes using BEDtools (version 2.29.2; using the parameters *getfasta -bed*)^98^. To identify genes that were conserved in the reference haplotype, we extracted the longest isoform for each protein-coding gene in this region from the N2 reference gene set and used exonerate^110^ (version 2.4.0; using the parameters *--model p2g --showvulgar no --showalignment no --showquerygff no --showtargetgff yes*) to infer gene models directly for each of the 15 isotypes. To predict genes that were not present in the reference haplotype, we first used BEDtools (version v2.29.2; with the parameter *maskfasta*) to mask the coordinates of the exonerate-predicted genes with Ns. We then used AUGUSTUS (version 3.3.3; using the parameters *--genemodel=partial --gff3=on --species=caenorhabditis --codingseq=on*) to predict genes in the unmasked regions. To remove spurious gene predictions or transposable element loci, we searched the protein sequence of each AUGUSTUS-predicted gene against the N2 reference genome using BLASTP (version 2.9.0+; using the parameters *-max_target_seqs 1 -max_hsps 1 -evalue 1e-5*) and against the Pfam database^111^ using InterProScan (version 5.35-74.0; using the parameters *-dp -t p --goterms -appl Pfam -f tsv*)^112^. Predicted genes that contained Pfam domains associated with transposable elements, or those genes that lacked sequence similarity to a known *C. elegans* protein sequence and a conserved protein domain were discarded (Supplementary Data 4). The coordinates of all curated predicted protein-coding genes were used to generate gene content plots (Fig 5a; Extended Data Fig. 8a,9a) using the ggplot R package^113^. To generate amino acid identity heatmaps and gene trees, we aligned the protein sequences of all conserved genes (those predicted with exonerate) using FSA (version 1.15.9)^114^. We inferred gene trees using IQ-TREE^115^ (version 2.0.3), allowing the best-fitting substitution model to be automatically selected for each alignment^116^. Gene trees were visualised using the ggtree R package. We calculated percentage identity matrices from the protein alignments using a custom Python script (available at https://github.com/AndersenLab/Ce-328pop-div). We then generated heatmaps, ordered by the strain relatedness tree inferred for the region, using the ggplot R package^113^. Files containing gene coordinates, protein alignments, gene trees, and percentage identity matrices for each characterised region are available at https://github.com/AndersenLab/Ce-328pop-div. To construct dendrograms of hyper-divergent regions, we first converted the hard-filtered VCF to a gds object using the *snpgdsVCF2GDS* function in the SNPrelate package^117^. Next, we calculated the pairwise identity-by-descent fraction for all strains using the *snpgdsIBS* function in the *SNPrelate* R package, and performing hierarchical clustering on the matrix using the *snpgdsHCluster* function in the *SNPrelate* package. We visualized the strain relatedness using ggtree^80^.

### Comparing divergence between *C. briggsae* and *C. nigoni*

To compare the divergence between hyper-divergent haplotypes with closely related *Caenorhabditis* species, we downloaded the protein sequences predicted from the genomes of *Caenorhabditis briggsae*^118^ and *Caenorhabditis nigoni*^119^ from WormBase (WS275). We clustered the longest isoform of each protein-coding gene for both species with the longest isoform of each protein-coding gene in all 16 *C. elegans* isotypes using OrthoFinder (version 2.3.11; using the parameter *-og*). We identified 8,741 orthogroups containing protein sequences that were present and single-copy in all 18 genomes and aligned their sequences using FSA (version 1.15.9). We calculated a percentage identity matrix for each protein alignment using a custom Python script (available at https://github.com/AndersenLab/Ce-328pop-div). For each of the 15 isotypes, we partitioned the 8,741 genes into those that were classified as being divergent and those that were classified as being non-divergent by our short-read classification approach. We then extracted the percentage identity of the allele of interest and the N2 alleles, and also the percentage identity between the corresponding ortholog in *C. briggsae* and *C. nigoni*. For each gene, we calculated mean identity between all non-divergent alleles and the correspond N2 alleles (and between their orthologs in *C. briggsae* and *C. nigoni*) and the mean identity between all hyper-divergent alleles and the corresponding N2 allele (and between their orthologs in *C. briggsae* and *C. nigoni*).

## Supporting information

Supplementary Data Guide

## Data availability

The raw short-read sequencing reads for strains used in this project are available from the NCBI Sequence Read Archive (Project PRJNA549503). The raw PacBio long-read data, along with the *de novo* assemblies and gene predictions, are available from the NCBI Sequence Read Archive (Project PRJNA692613). Strain information and short-read genomic variation data are available at the *C. elegans* Natural Diversity Resource (www.elegansvariation.org)^69^.

## Code availability

All data sets and code for generating figures and tables are available on GitHub (https://github.com/AndersenLab/Ce-328pop-div).

## Acknowledgments

We would like to thank members of the Andersen lab for providing comments on this manuscript. We especially thank Michael Ailion, Jean David, Robert Luallen, Nathalie Pujol, and citizen-scientists for their contributions of wild *C. elegans* strains to CeNDR. We also thank the Duke University School of Medicine for the use of the Sequencing and Genomic Technologies Shared Resource, which provided Pacific Biosystems long-read sequencing.

## Author contributions

D.L., S.Z., and E.C.A. conceived and designed the study. D.L., S.Z., L.S., and E.C.A. analyzed the data and wrote the manuscript. Y.W., R.E.T., and D.E.C. performed whole-genome sequencing and isotype characterization for 609 wild *C. elegans* strains. R.E.T. performed long-read sequencing for 11 *C. elegans* wild isolates. R.C., A.K.W., and L.R.B. performed long-read sequencing for three *C. elegans* wild isolates. M.G.S., C.B., M.V.R., and M.-A.F. contributed wild isolates to the *C. elegans* strain collection and edited the manuscript. T.A.C. edited the manuscript.

## Additional information

**Supplementary information** is available for this paper.

**Correspondence and requests for materials** should be addressed to E.C.A.

## Extended Data Figures

**Extended Data Fig. 1.**
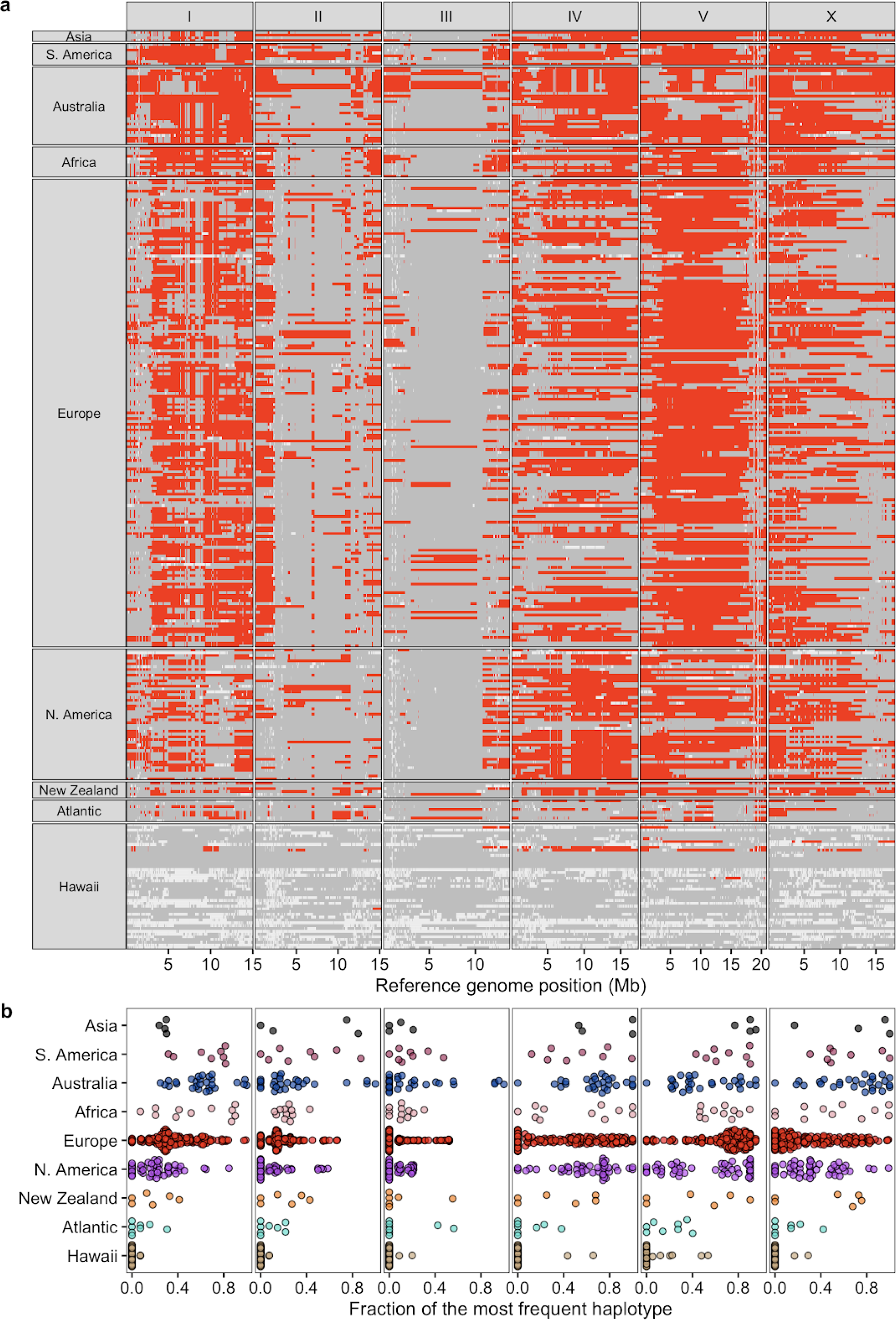
Chromosome-scale selective sweeps across wild *C. elegans* isotypes. (a) The genome-wide distribution of the most frequent haplotype (red) among 324 wild isotypes with known geographic origin is shown. Gray genomic regions represent other haplotypes, and white represents unclassified haplotypes. Each row is one of the 324 isotypes, grouped by the geographic origin. The genomic position in Mb is plotted on the x-axis, and each tick mark represents 5 Mb of the chromosome. (b) Beeswarm plots of the proportion of the most frequent haplotype for each chromosome from (a) for 324 isotypes with known geographic origins are shown. Wild isotypes are grouped by geographic origin. Each point corresponds to one of the 324 isotypes, and geographic origins are shown on the y-axis.

**Extended Data Fig. 2.**
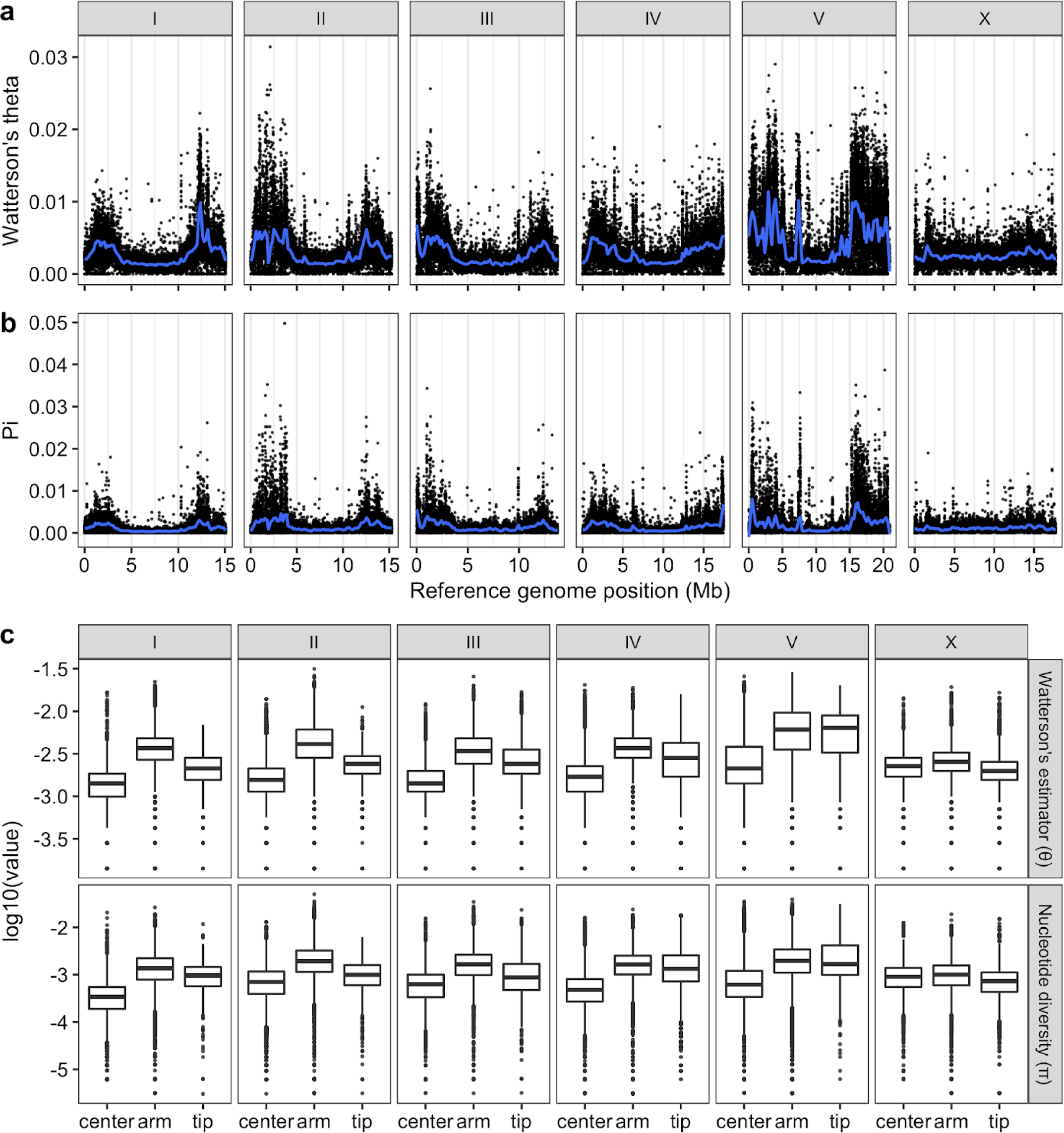
Patterns of molecular diversity across the *C. elegans* genome. The chromosomal patterns (a) Watterson’s theta (θ) and (b) nucleotide diversity (pi) for non-overlapping 1 kb windows are shown. Each dot corresponds to the calculated value for a particular window. The genomic position in Mb is plotted on the x-axis. Diversity statistic values are shown on the y-axis. Smoothed lines (blue) are LOESS fits. (c) Tukey box plots of genetic diversity statistics from (a) are shown with outlier data points plotted. Genetic diversity statistics for each sliding window are grouped by the chromosomal region defined previously (Rockman and Kruglyak, 2009). Genetic diversity statistic values are shown on the y-axis. The horizontal line in the middle of the box is the median, and the box denotes the 25th to 75th quantiles of the data. The vertical line represents the 1.5x interquartile range.

**Extended Data Fig. 3.**
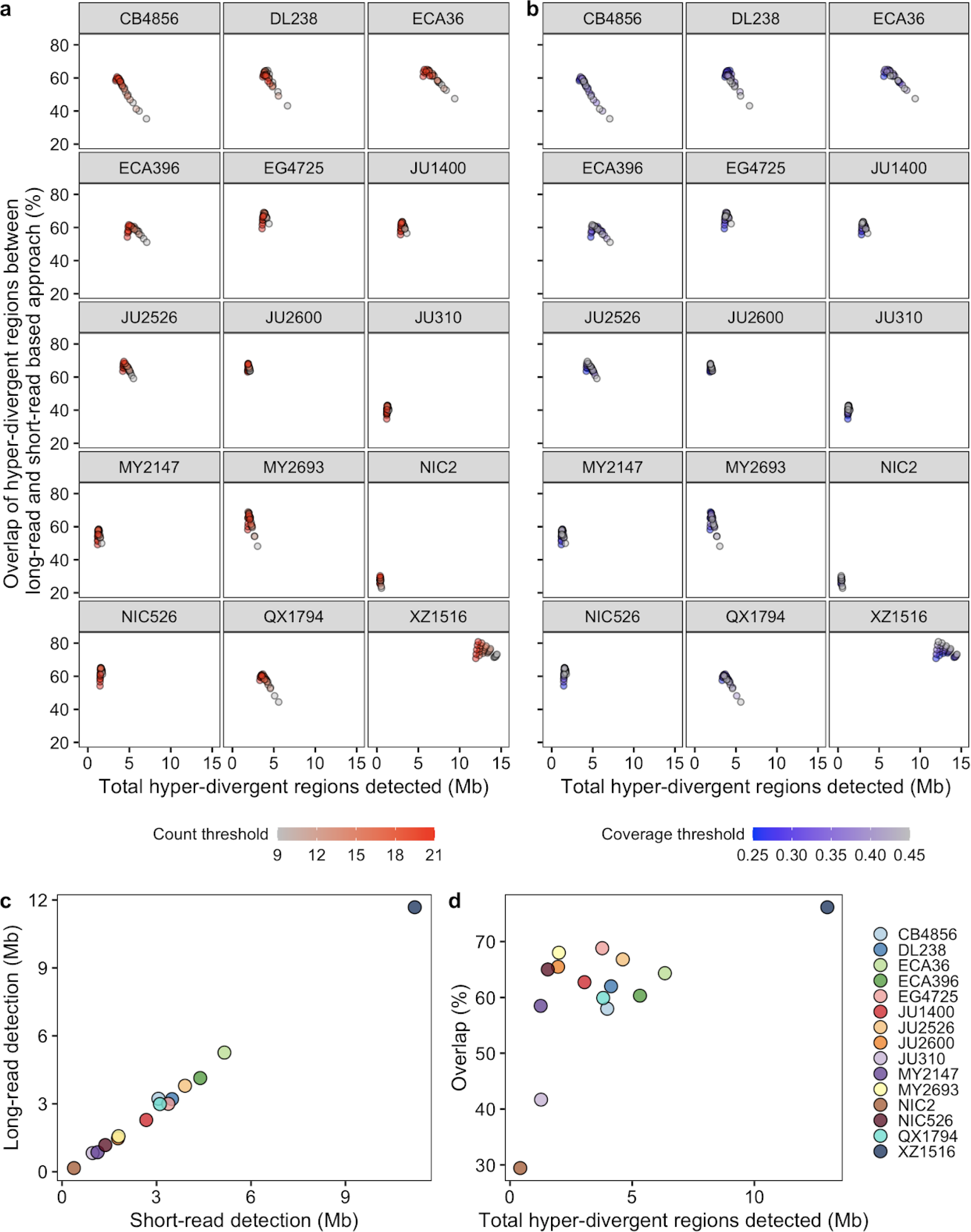
Optimization of parameters for the characterization of hyper-divergent regions. (a,b) The total detected hyper-divergent regions in Mb (x-axis) and the percent overlap of long-read and short-read hyper-divergent classification (y-axis) are shown (Methods). Each point corresponds to one of the combination of threshold parameters for the variant count and coverage fraction of 1 kb bin to be classified as hyper-divergent. Each point is colored by the variant count threshold (a) or the coverage fraction threshold (b). (c) The relationship between the total size of hyper-divergent regions detected by the optimized short-read or long-read based approach is shown. Each point corresponds to one of the 15 long-read sequenced isotypes. Total sizes of hyper-divergent regions detected by the short-read based approach are shown on the x-axis, and total sizes of hyper-divergent regions detected by the long-read based approach are shown on the y-axis. (d) The overlap between hyper-divergent regions defined by the optimized short-read based approach and long-read based approach is shown. Each point corresponds to one of the 15 long-read sequenced isotypes. Total sizes of hyper-divergent regions detected by either short-read or long-read based approach are shown on the x-axis, and the percentages of hyper-divergent regions detected by both approaches are shown on the y-axis.

**Extended Data Fig. 4.**
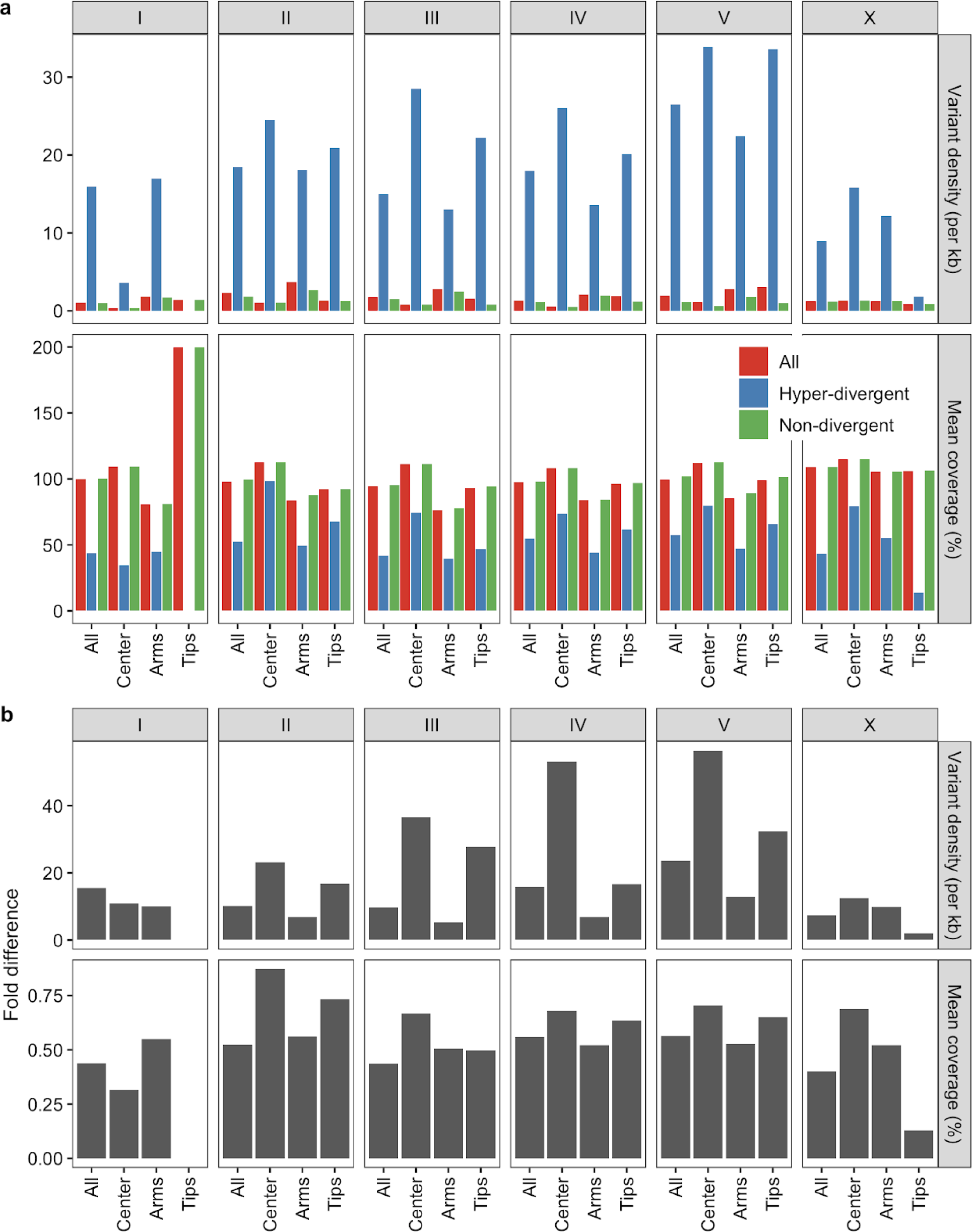
Summary statistics for hyper-divergent regions across six chromosomes. (a) Bar plots for the comparisons of variant (SNV/indel) density (top) and coverage fraction (bottom) between hyper-divergent regions (red) and the rest of the regions (blue) in each chromosomal region are shown. Note that no hyper-divergent region was found on the tips of chromosome I. (b) Fold differences between hyper-divergent regions and the rest of the regions from (a) are shown.

**Extended Data Fig. 5.**
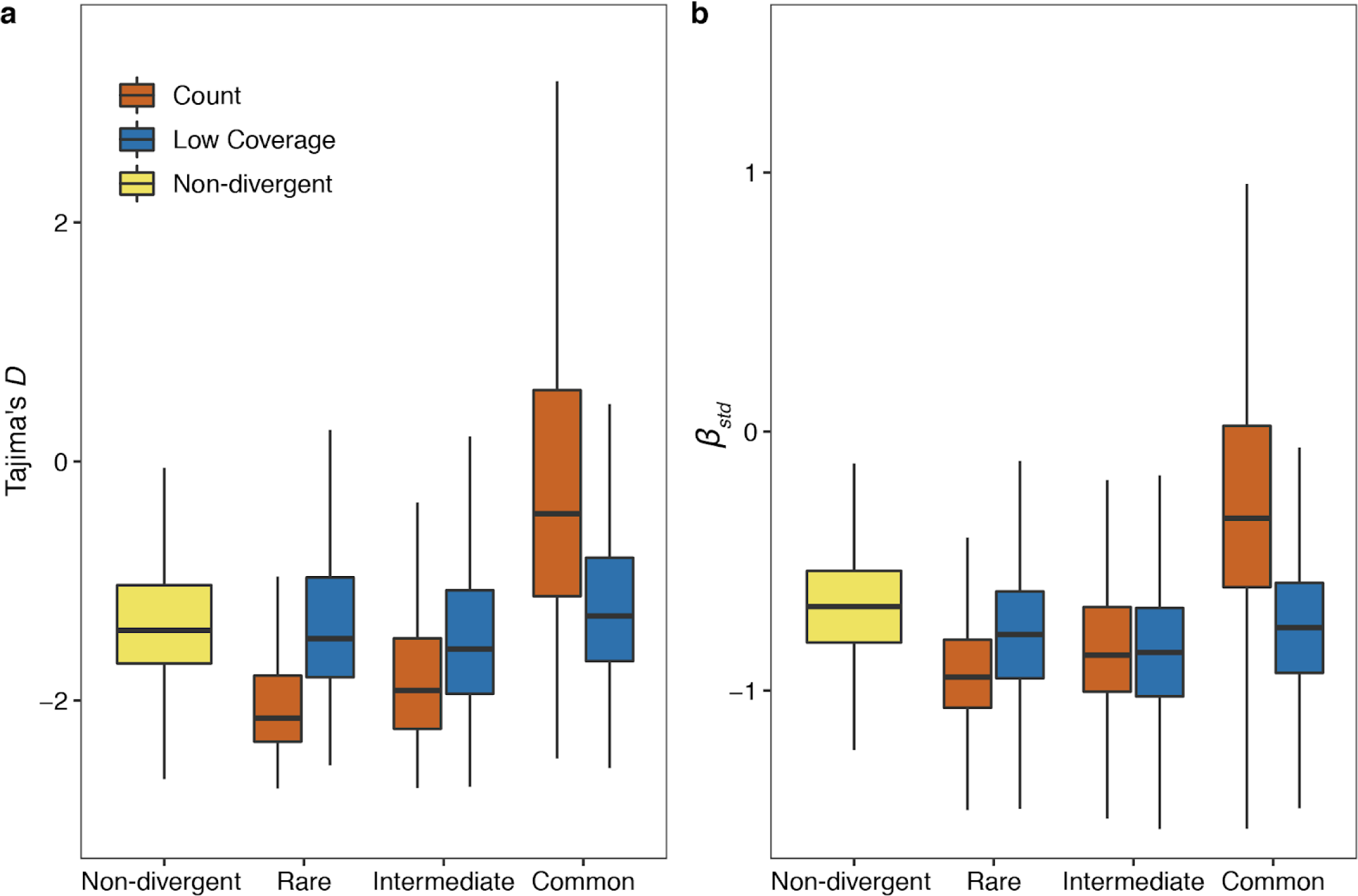
Genomic signatures of balancing selection in non-divergent regions and hyper-divergent regions. Tukey box plots of Tajima’s *D* (a) and standardized beta (b) are shown. Genomic bins (1kb) (a) or variants (b) are grouped and colored by their classification: (1) non-divergent bins (yellow), (2) hyper-divergent bins with high variant density (> 16 SNVs/indels, red), (3) hyper-divergent bins with low read depth (< 35%, blue). Hyper-divergent bins are grouped by their species-wide frequencies: rare (<1%), intermediate (≥ 1% and < 5%), or common (≥ 5%). The horizontal line in the middle of the box is the median, and the box denotes the 25th to 75th quantiles of the data. The vertical line represents the 1.5x interquartile range.

**Extended Data Fig. 6.**
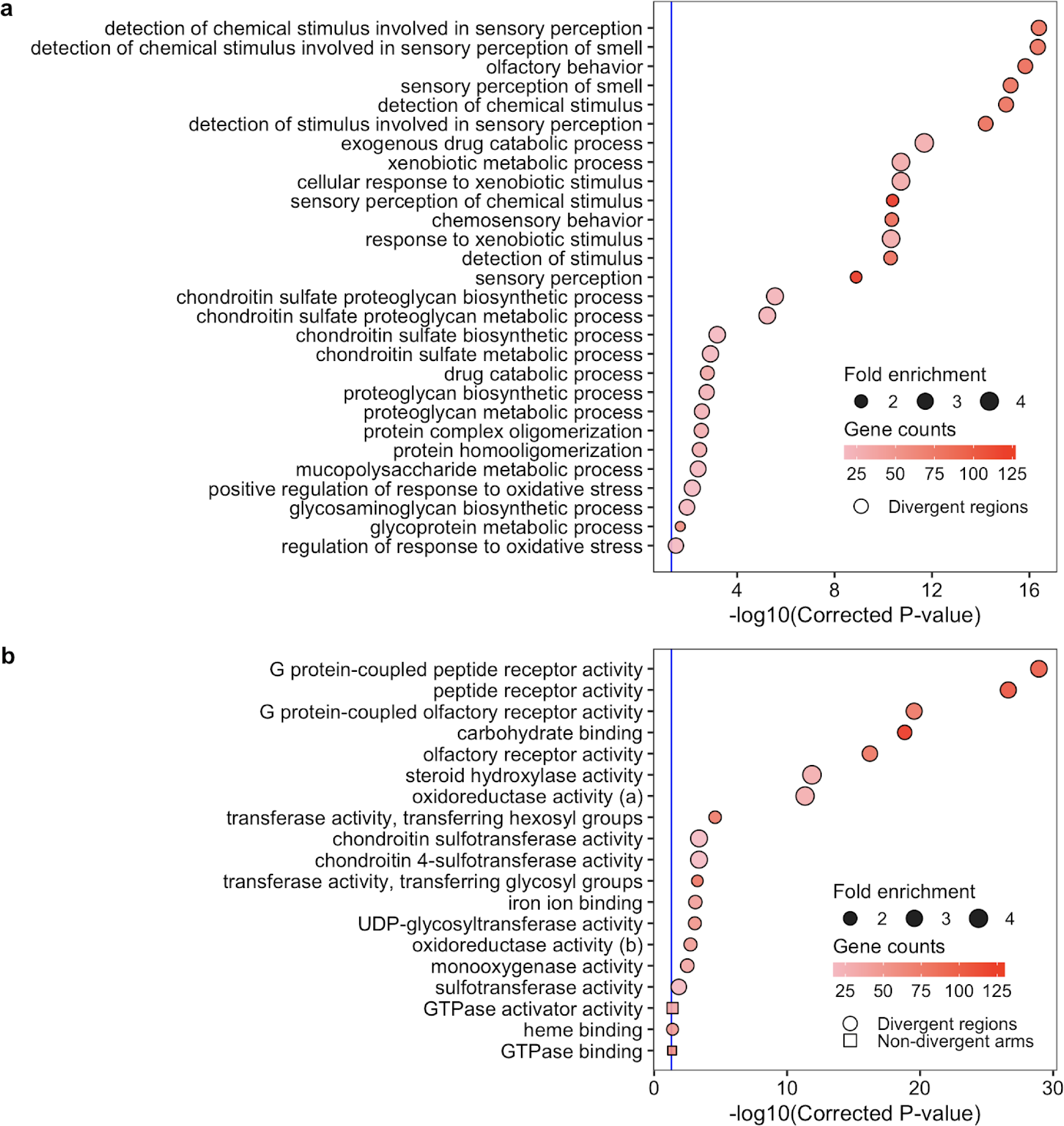
Gene ontology (GO) enrichment for hyper-divergent regions. Gene ontology (GO) enrichment for the biological process category (a) and the molecular function category (b) for non-divergent chromosomal arms (square) and hyper-divergent regions (circle) are shown. Significantly enriched GO terms in control regions or hyper-divergent regions or both are shown on the y-axis. Bonferroni-corrected significance values for GO enrichment are shown on the x-axis. Sizes of squares and circles correspond to the fold enrichment of the annotation, and colors of square and circle correspond to the gene counts of the annotation. The blue line shows the Bonferroni-corrected significance threshold (corrected p-value = 0.05). Note, we did not detect any GO-term enrichment of genes in non-divergent chromosomal arms for the biological process category.

**Extended Data Fig. 7.**
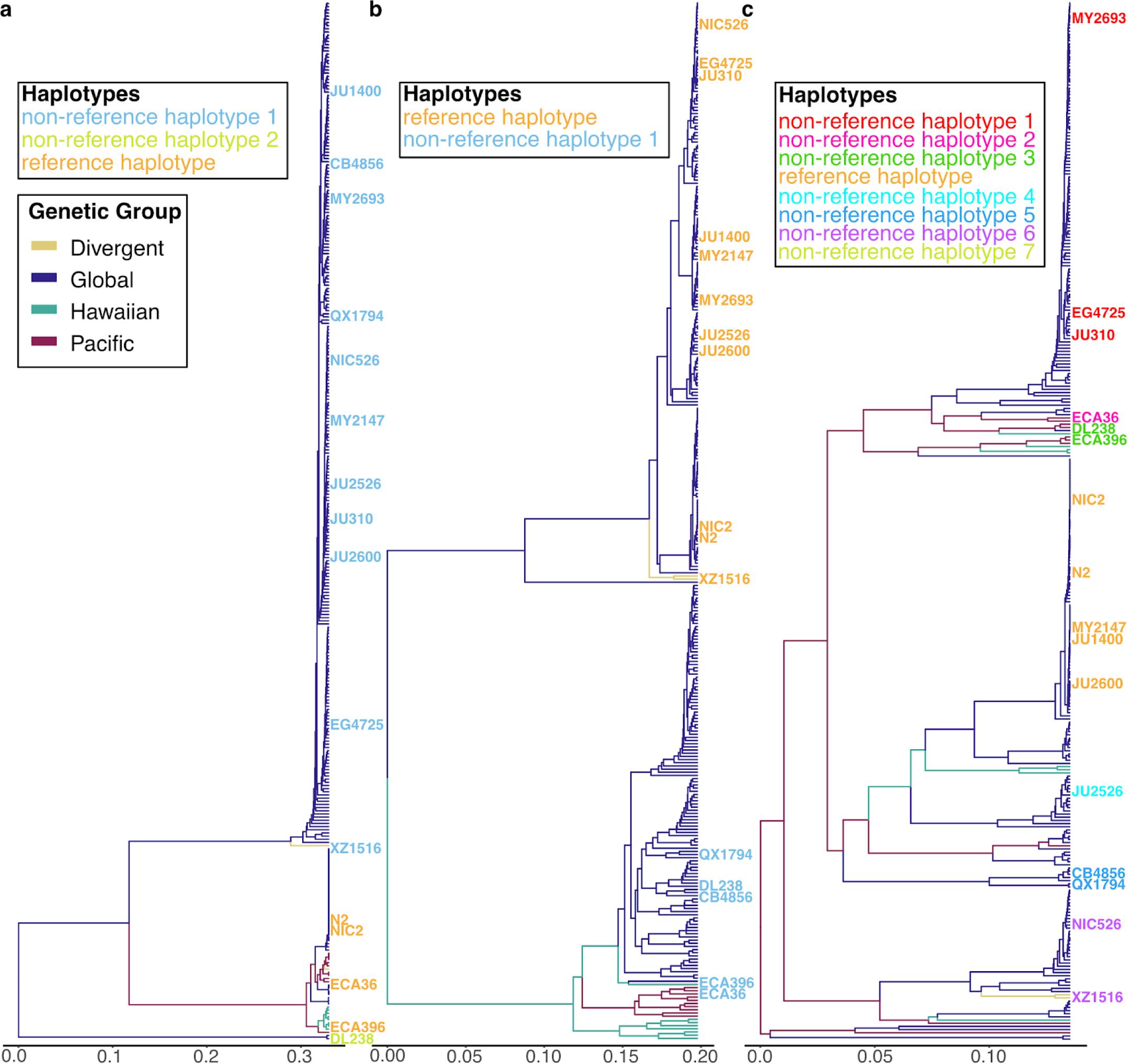
Species-wide SNP-based relatedness of divergent regions is in agreement with long-read sequencing results. The inferred for the *C. elegans* species-wide relatedness for the hyper-divergent regions that span (a) II:3,667,179-3,701,405, (b) I:2,318,291-2,381,851, and (c) V:20,193,463-20,267,244 are shown. The x-axis represents the dissimilarity of the fraction of identity-by-state in the region. For a-c, the isotype names are colored to match the haplotypes defined by long-read sequence data in Fig. 5 and Extended Fig. 8,9, respectively. The branch colors correspond to the species-wide genetic groups identified by PCA analysis in Fig. 1c.

**Extended Data Fig. 8.**
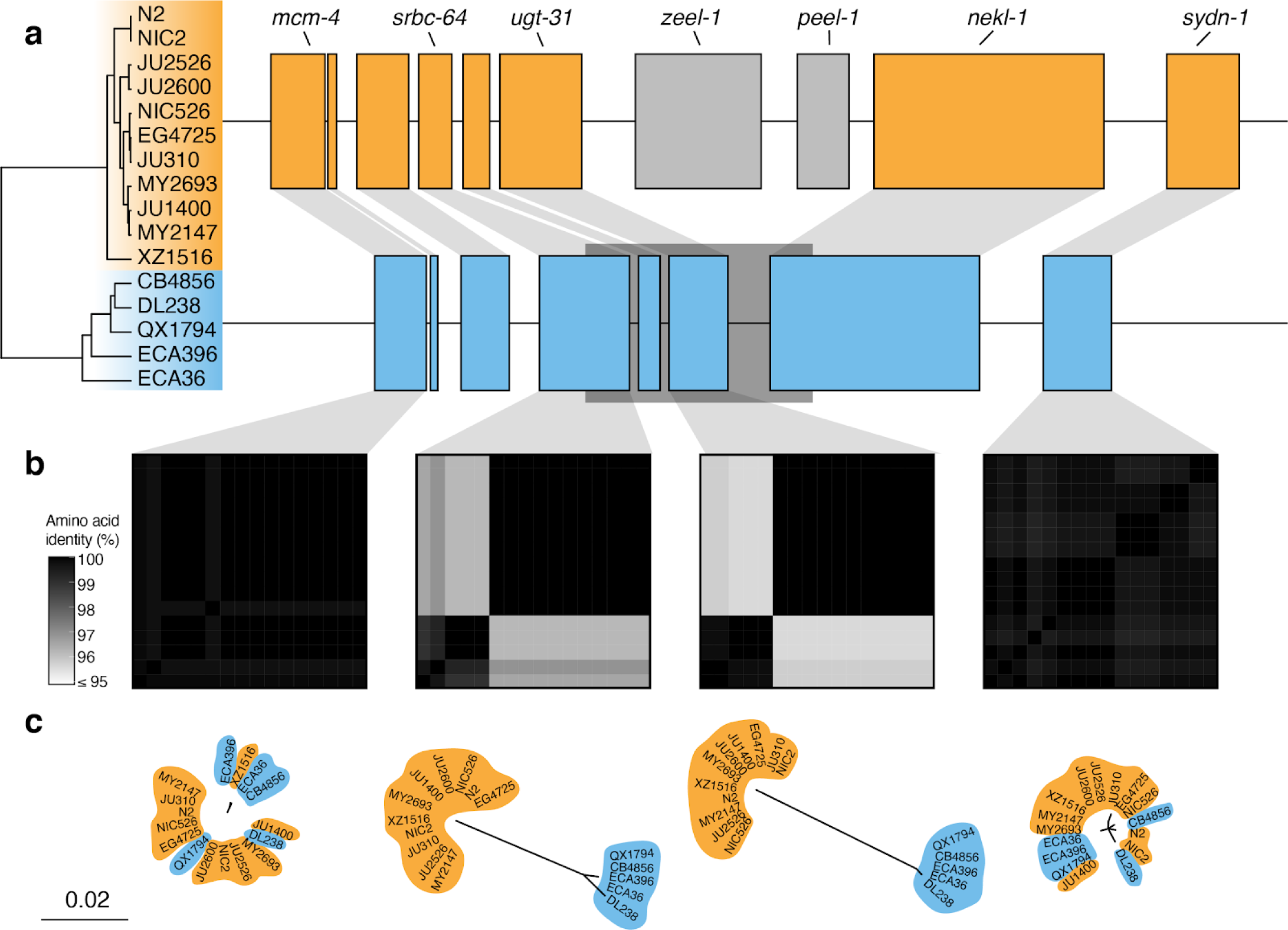
Two hyper-divergent haplotypes at the *peel-1 zeel-1* incompatibility locus. (a) The protein-coding gene contents of the two hyper-divergent haplotypes at the *peel-1 zeel-1* incompatibility locus on the left arm of chromosome I (I:2,318,291-2,381,851 of the N2 reference genome). The tree was inferred using SNVs and colored by inferred haplotypes. For each distinct haplotype, we chose a single isotype as a haplotype representative (orange haplotype: N2, blue haplotype: CB4856) and predicted protein-coding genes using both protein-based alignments and *ab initio* approaches. Protein-coding genes are shown as boxes; those genes that are conserved in all haplotypes are colored based on their haplotype, and those genes that are not are colored light gray. Dark gray boxes behind genes indicate coordinates of divergent regions. Genes with locus names in N2 are highlighted. (b) Heatmaps showing amino acid identity for alleles of four genes (*mcm-4*, *srbc-64*, *ugt-31*, and *sydn-1*). The percentage identity was calculated using alignments of protein sequences from all 16 isotypes. Heatmaps are ordered by the SNV tree shown in (a). (c) Maximum-likelihood gene trees of four genes (*mcm-4*, *srbc-64*, *ugt-31*, and *sydn-1*) inferred using amino acid alignments. Trees are plotted on the same scale (scale shown; scale is in substitutions per site). Strain names are colored by their haplotype.

**Extended Data Fig. 9.**
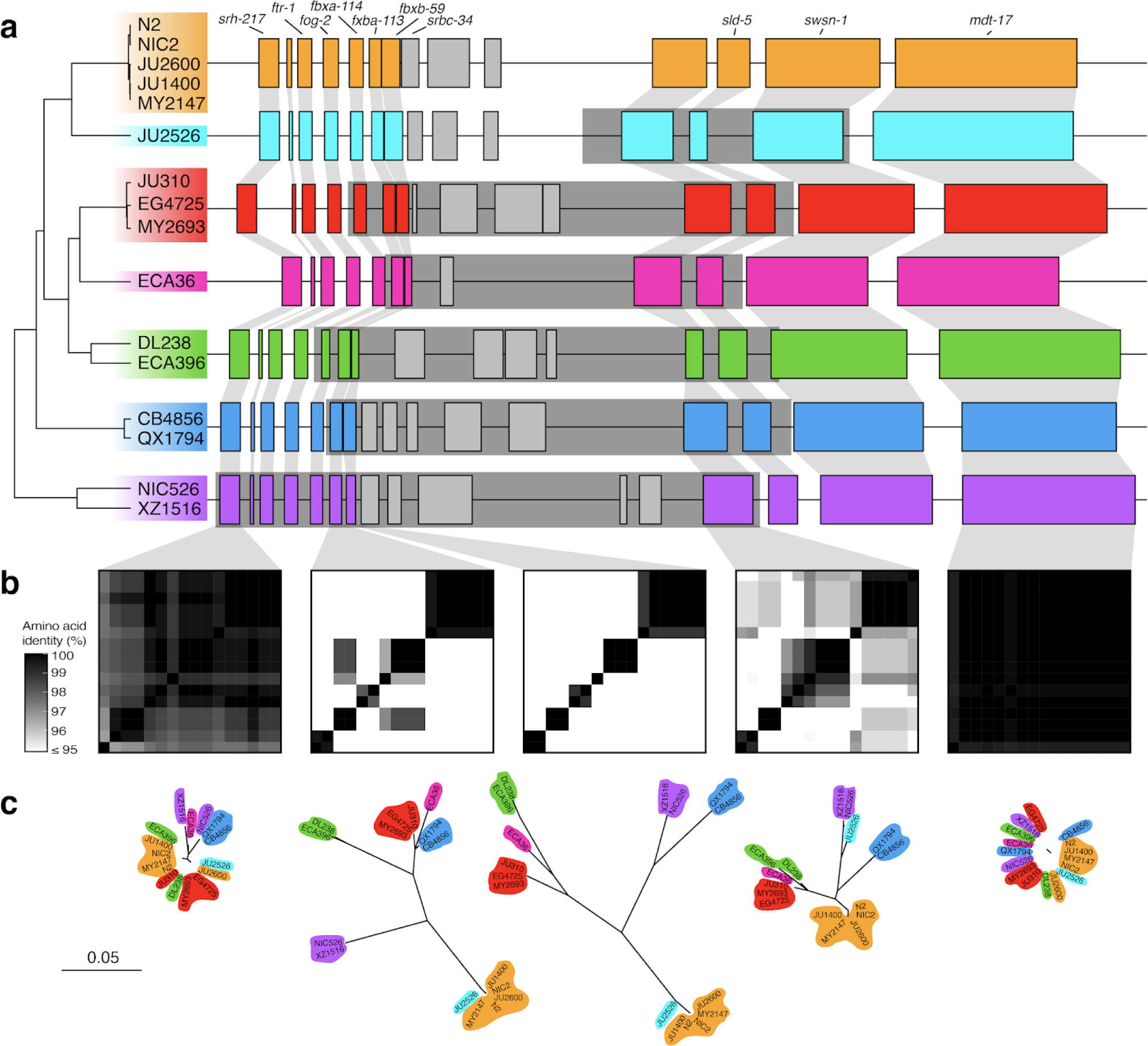
Hyper-divergent haplotypes at a region on the right arm of chromosome V. (a) The protein-coding gene contents of the seven hyper-divergent haplotypes at a region on the right arm of chromosome V (V:20,193,463-20,267,244 of the N2 reference genome). The tree was inferred using SNVs and colored by inferred haplotypes. For each distinct haplotype, we chose a single isotype as a haplotype representative (orange haplotype: N2, light blue haplotype: JU2526, red haplotype: EG4725, pink haplotype: ECA36, green haplotype: DL238, dark blue haplotype: QX1794, purple haplotype: NIC526) and predicted protein-coding genes using both protein-based alignments and *ab initio* approaches. JU2526 shares the reference haplotype at *fbxa-113* and *fbxb-59* (six hyper-divergent haplotypes at these loci) but is divergent at *Y113G7B.15* (seven hyper-divergent haplotypes at this locus). Protein-coding genes are shown as boxes; those genes that are conserved in all haplotypes are colored based on their haplotypes, and those genes that are not are colored light gray. Dark gray boxes behind genes indicate coordinates of divergent regions. Genes with locus names in N2 are highlighted. Of the 25 genes that are not conserved in all haplotypes (light gray boxes), ten are alleles of the three reference haplotype (N2) loci colored in light gray. The remaining 15 do not have a clear one-to-one relationship with a gene in the reference haplotype. Seven of these 15 have homology to *F54E12.2* (present in the reference haplotype) and are likely the product of duplication and diversification. Six have homology to either *M04C3.1, F19B2.5*, or *F54E12.2*, all of which are genes with SNF2 family N-terminal domains and which exist elsewhere in the N2 reference genome. Of the remaining two genes, one has homology to *Y113G7B.15*, which is present in the reference haplotype, and the other has homology to *W09C3.8*, a gene on chromosome I in the reference genome. Functional annotations of all unconserved loci (including BLAST hits and Pfam domains identified by InterProScan) can be found in Supplementary Data 4. (b) Heatmaps showing amino acid identity for between alleles of five genes (*srh-217*, *fbxb-113*, *fbxb-59*, *Y113G7B.15*, and *mdt-17*). The percentage identity was calculated using alignments of proteins sequences from all 16 isotypes. Heatmaps are ordered by the SNV tree shown in (a). (c) Maximum-likelihood gene trees of five genes (*srh-217*, *fbxb-113*, *fbxb-59*, *Y113G7B.15*, and *mdt-17*) inferred using amino acid alignments. Trees are plotted on the same scale (scale shown; scale is in substitutions per site). Strain names are colored by their haplotype.

**Extended Data Fig. 10.**
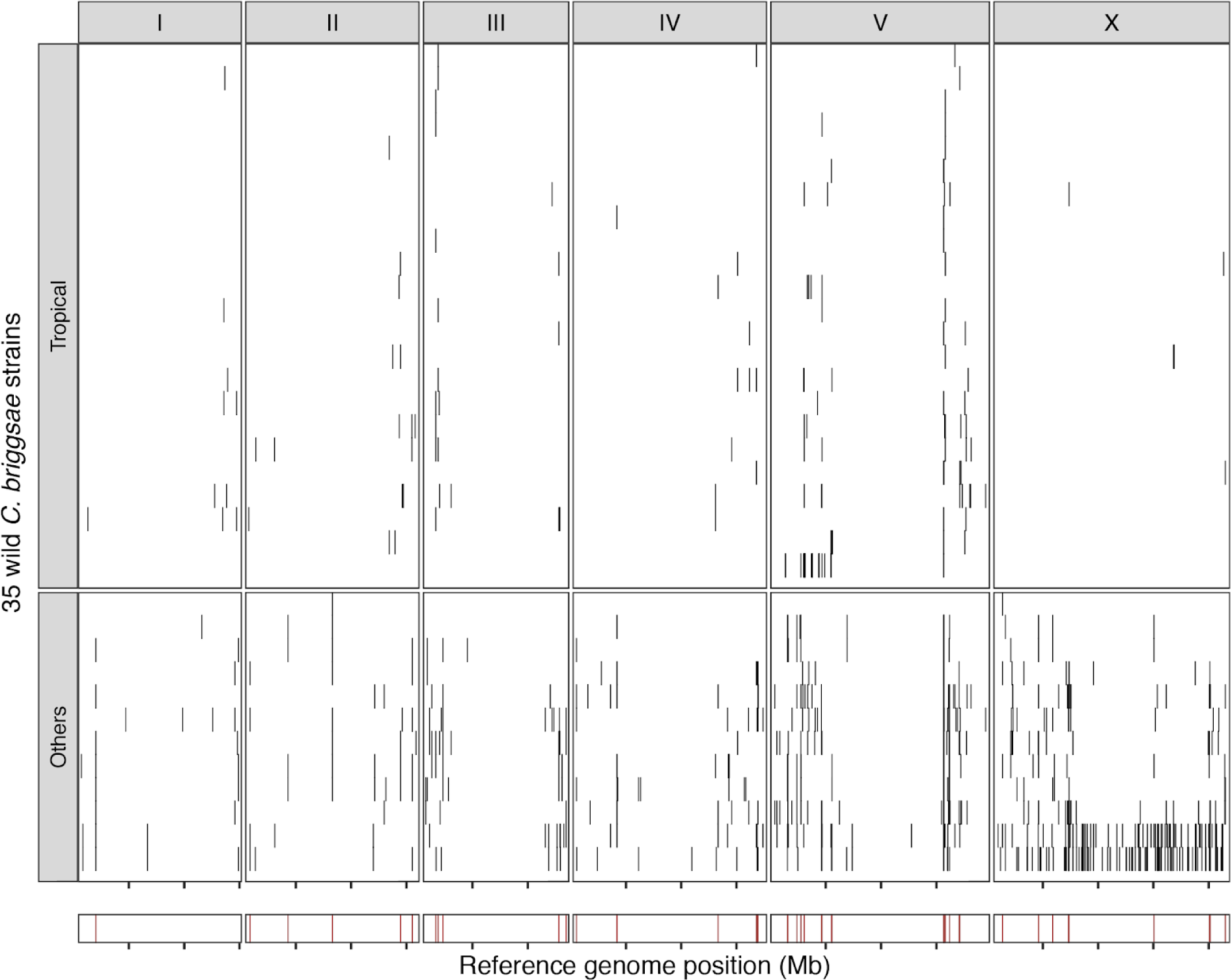
Hyper-divergent regions in *C. briggsae*. The genome-wide distribution of hyper-divergent regions across 35 non-reference wild *C. briggsae* strains is shown. In the top panel, each row is one of the 35 strains, grouped by previously defined clades (tropical or others) ordered by the total amount of genome covered by hyper-divergent regions (black). In the bottom panel, brown bars indicate genomic positions in which more than 10% of strains are classified as hyper-divergent at the locus. The genomic position in Mb is plotted on the x-axis, and each tick represents 5 Mb of the chromosome.

