## Supplementary Information for "Balancing selection maintains hyper-divergent haplotypes in *C. elegans*"

Daehan Lee[#], Stefan Zdraljevic[#], Lewis Stevens[#], Ye Wang, Robyn E. Tanny, Timothy A. Crombie, Daniel E. Cook, Amy K. Webster, Rojin Chirakar, L. Ryan Baugh, Mark G. Sterken, Christian Braendle, Marie-Anne Félix, Matthew V. Rockman, Erik C. Andersen[*]

### These authors contributed equally to this work.

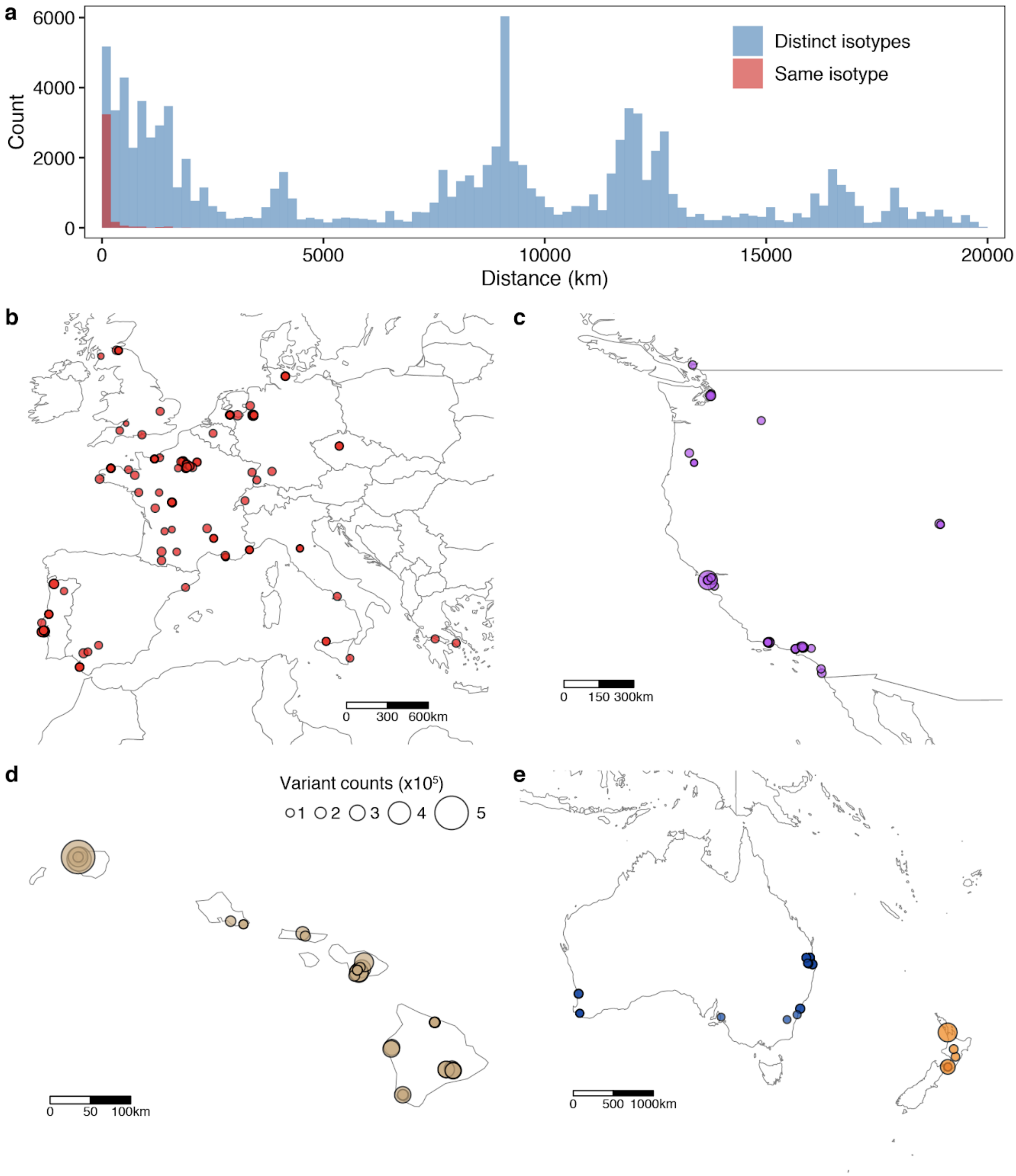
a
6000
4000
2000
0
Count
Distinct isotypes
Same isotype
0
5000
10000
15000
20000
Distance (km)
b
0 300 600km
c
0 150 300km
d
Variant counts (x10^5)
1 2 3 4 5
0 50 100km
e
0 500 1000km

**Supplementary Fig. 1 | Geographic distribution of *C. elegans* isotypes**

(a) A histogram for the pairwise distances between locations of wild strain isolations with the same isotype (red) or distinct isotypes (blue). Distances between two isolations are shown on the x-axis, and counts for the wild strain pair are shown on the y-axis.

(b-e) The geographic distributions of isotypes from (b) Europe, (c) Pacific coast of North America, (d) Hawaii Islands, and (e) Australia and New Zealand are shown. Each circle corresponds to one of the wild isotypes, and the size of each circle corresponds to the number of non-reference homozygous alleles.

a

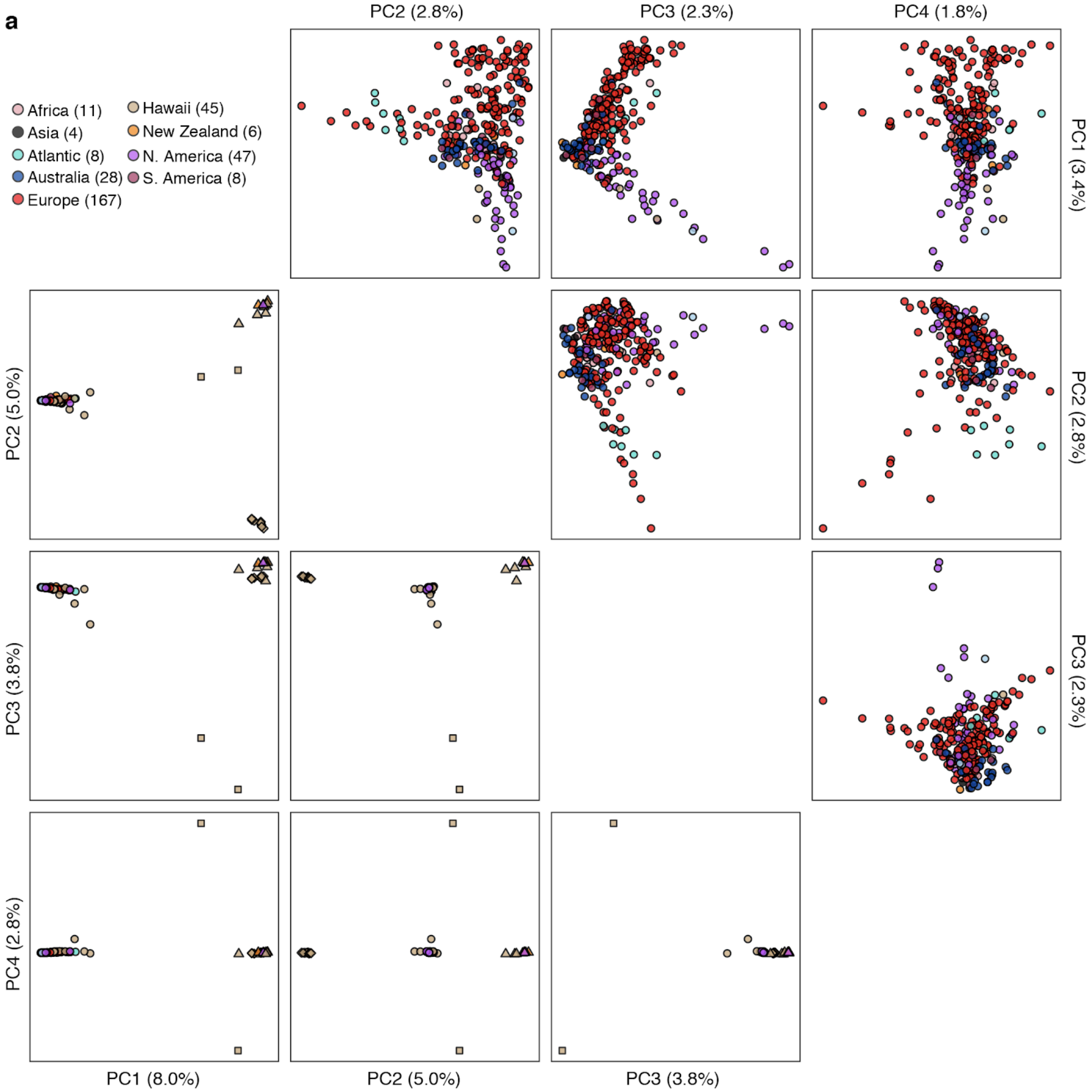

Africa (11)
Asia (4)
Atlantic (8)
Australia (28)
Europe (167)
Hawaii (45)
New Zealand (6)
N. America (47)
S. America (8)
PC2 (2.8%)
PC3 (2.3%)
PC4 (1.8%)
PC1 (3.4%)
PC2 (2.8%)
PC3 (2.3%)
PC2 (5.0%)
PC3 (3.8%)
PC4 (2.8%)
PC1 (8.0%)
PC2 (5.0%)
PC3 (3.8%)

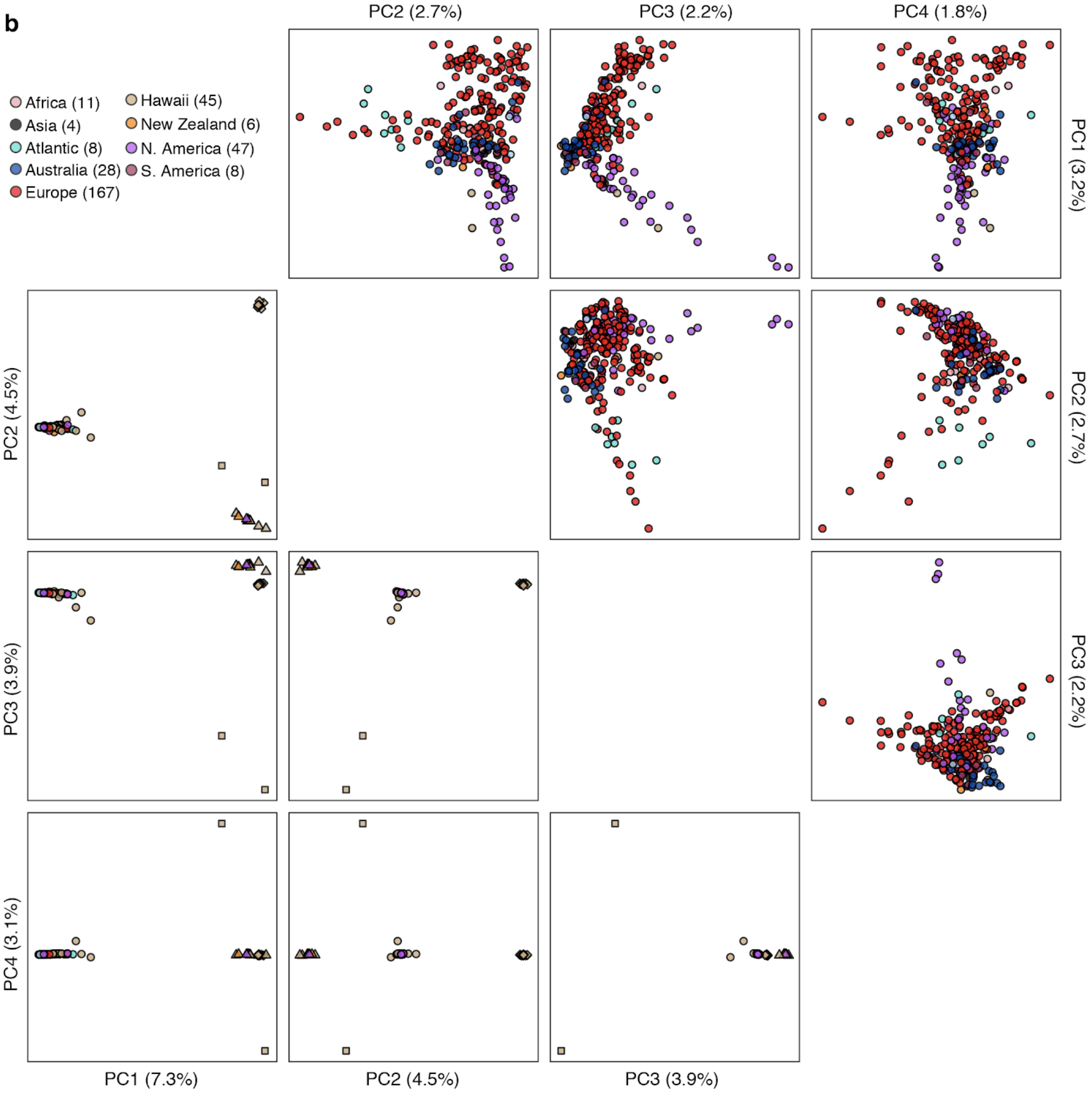

b
Africa (11)
Asia (4)
Atlantic (8)
Australia (28)
Europe (167)
Hawaii (45)
New Zealand (6)
N. America (47)
S. America (8)
PC2 (2.7%)
PC3 (2.2%)
PC4 (1.8%)
PC1 (3.2%)
PC2 (2.7%)
PC3 (2.2%)
PC2 (4.5%)
PC3 (3.9%)
PC4 (3.1%)
PC1 (7.3%)
PC2 (4.5%)
PC3 (3.9%)

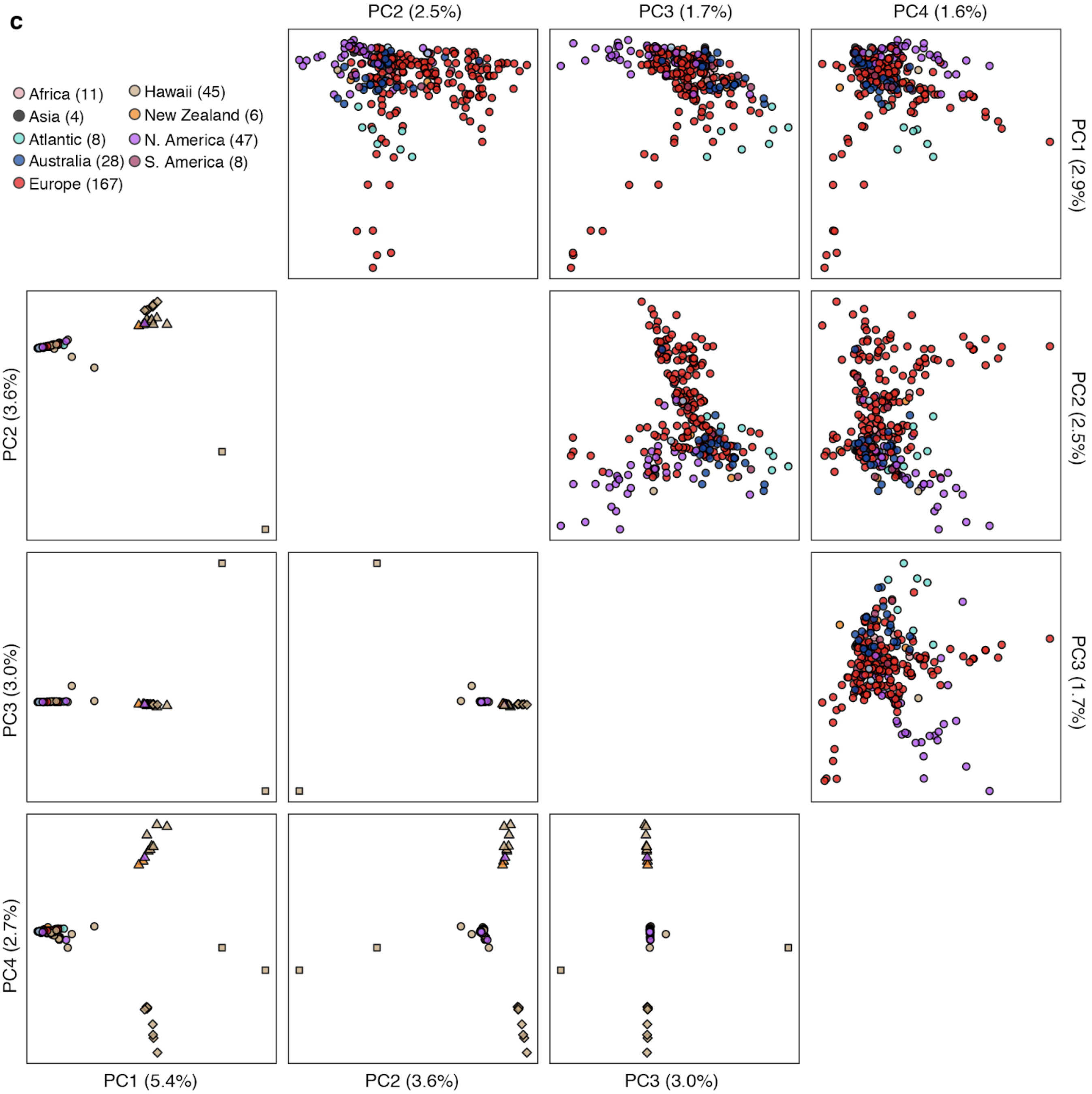


**Supplementary Fig. 2 | The population structure of 328 wild *C. elegans* isotypes**

Plots of the 324 isotypes from Global (circle), Pacific (triangle), and Hawaii (diamond) and two divergent isotypes (XZ1516 and ECA701, square) according to their values for each of the four significant axes of variation, as determined by principal component analysis (PCA) of the genotype covariances from LD-pruned genotypes matrices (LD threshold: 0.8 (a), 0.6 (b), 0.2 (c)). (Lower triangle (bottom left plots): without outlier removal, upper triangle (top right plots): with outlier removal. See Methods.) Each point is one of the 328 isotypes, which is colored by the geographic origin of the isotype.

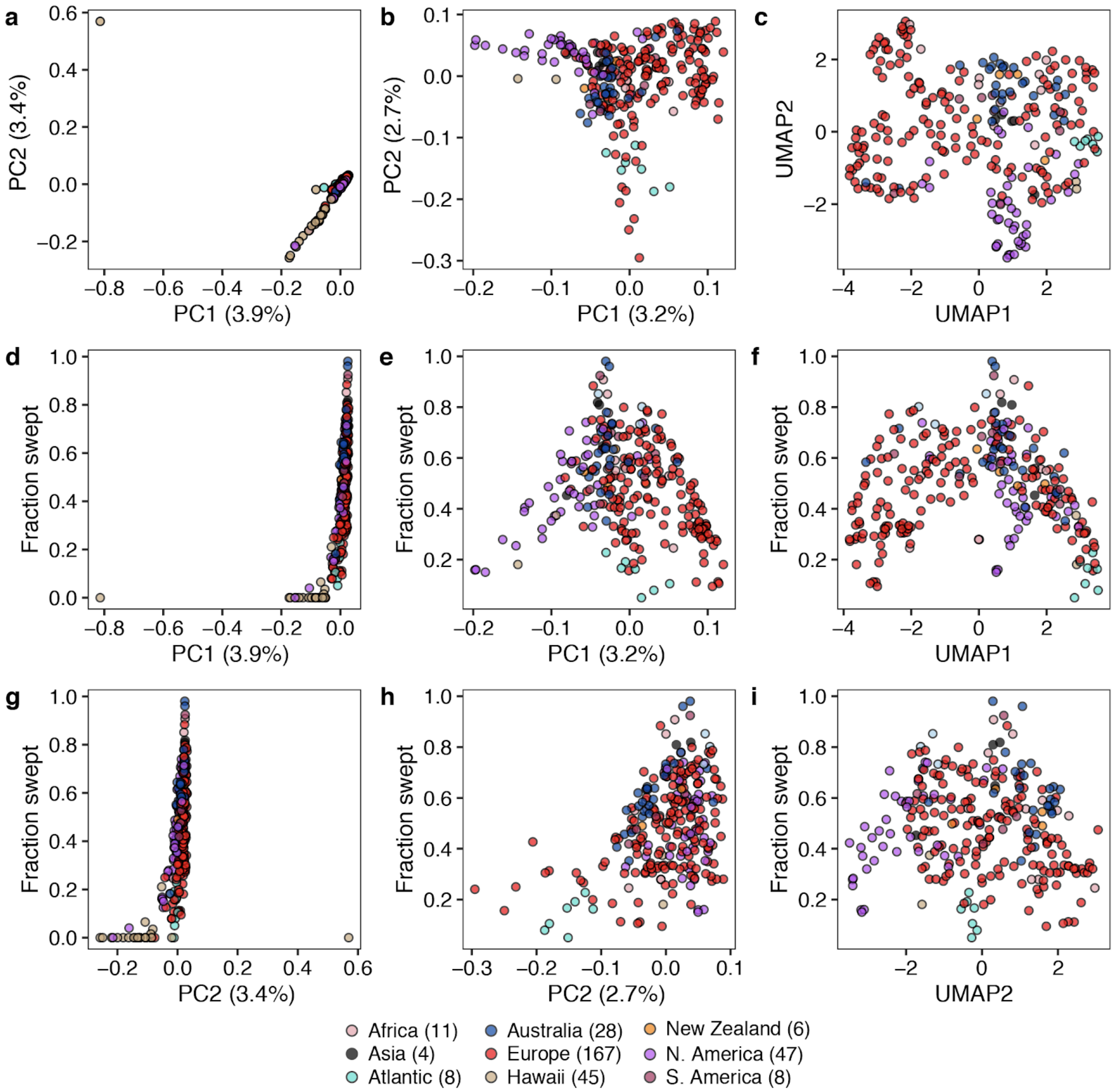


**Supplementary Fig. 3 | Selective sweeps shape population structure of the Global group**
(a) Plot of 308 isotypes that belong to the Global group according to their values for each of the two significant axes of variation, as determined by PCA of the genotype covariances. Each point is one of the 308 isotypes, which is colored by their geographic origins.
(b) Plot of 251 isotypes from the Global group according to the values for each of the two significant axes of variation, as determined by PCA of the genotype covariances with five iterations of outlier removal. Each point is one of the 251 isotypes, which is colored by the geographic origin.
(c) Plot of 251 isotypes from the Global group according to the values for each of the two significant axes of variation, as determined by Uniform Manifold Approximation and Projection (UMAP) analysis using five principal components from PCA of the genotype covariances with five iterations of outlier removal. Each point is one of the 251 isotypes, which is colored by the geographic origin.
(d-i) Scatter plots for the fraction of swept chromosomes for the isotypes in (a-c), respectively. The first (d-f) (d-f) or second significant axis (g-i) of variation defined in (a-c) is shown on the x-axis, and the fraction of swept chromosomes is shown on the y-axis. Each point corresponds to one of 308 (d, g) or 251 (e-f, h-i) isotypes and is colored by the geographic origin.

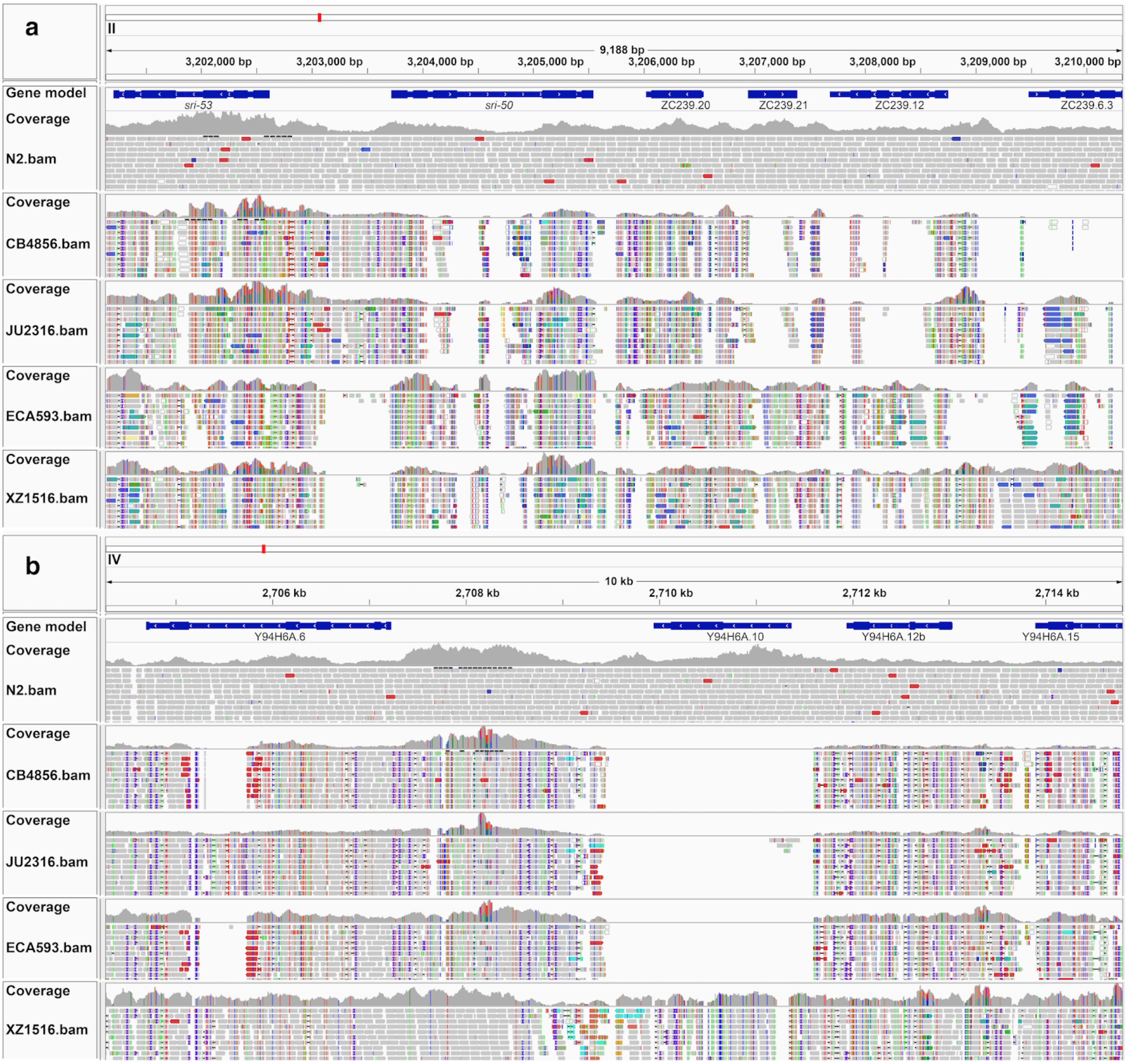


**Supplementary Fig. 4 | Short-read alignments of wild isotypes in hyper-divergent regions**
Short-read alignments from five isotypes (N2, CB4856, JU2316, ECA593, and XZ1516) to the N2 reference genome (WS245) for (a) a region of chromosome II (II:3,200,000-3,211,000) and (b) a region of chromosome IV (IV:2,704,000-2,715,000) plotted by IGV 2.8.0 are shown. For each plot, genes in the interval are shown at the top. For each isotype, the top panel shows the coverage of genomic positions, and the bottom panel shows aligned short-reads at genomic positions (gray: normal reads, red: reads with putative deletion, blue: reads with putative insertion, navy and turquoise: reads with putative inversion, green: reads with putative duplication or translocation). Colored vertical lines indicate mismatched bases at the position.

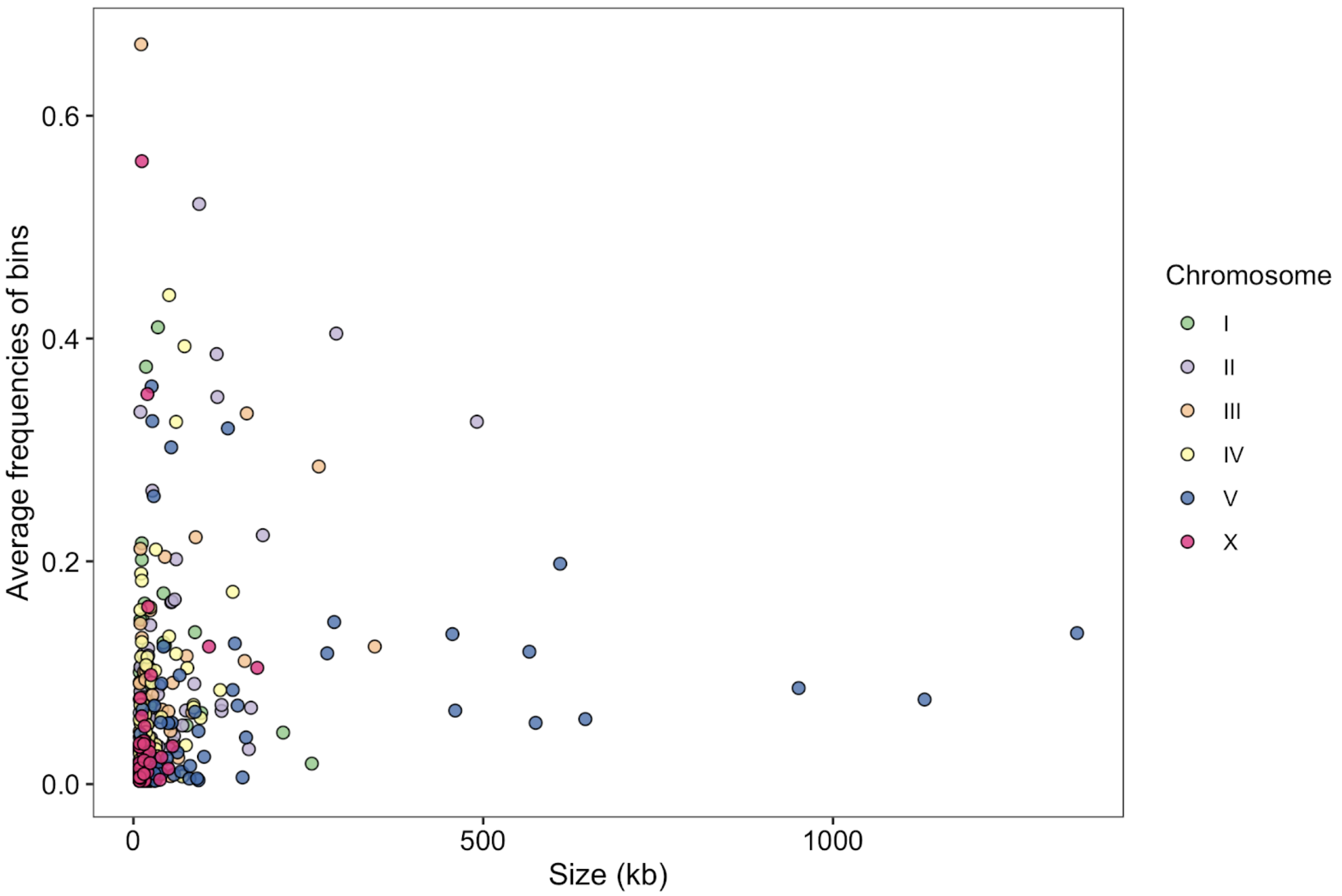


**Supplementary Fig. 5 | Sizes and frequencies of population-wide hyper-divergent blocks**
The population-wide hyper-divergent block sizes with respect to the N2 reference genome (x-axis) and the average frequencies of 1 kb bins that are classified as hyper-divergent across 327 non-reference isotypes (y-axis) is shown. Each point corresponds to one of the 366 population-wide hyper-divergent blocks.

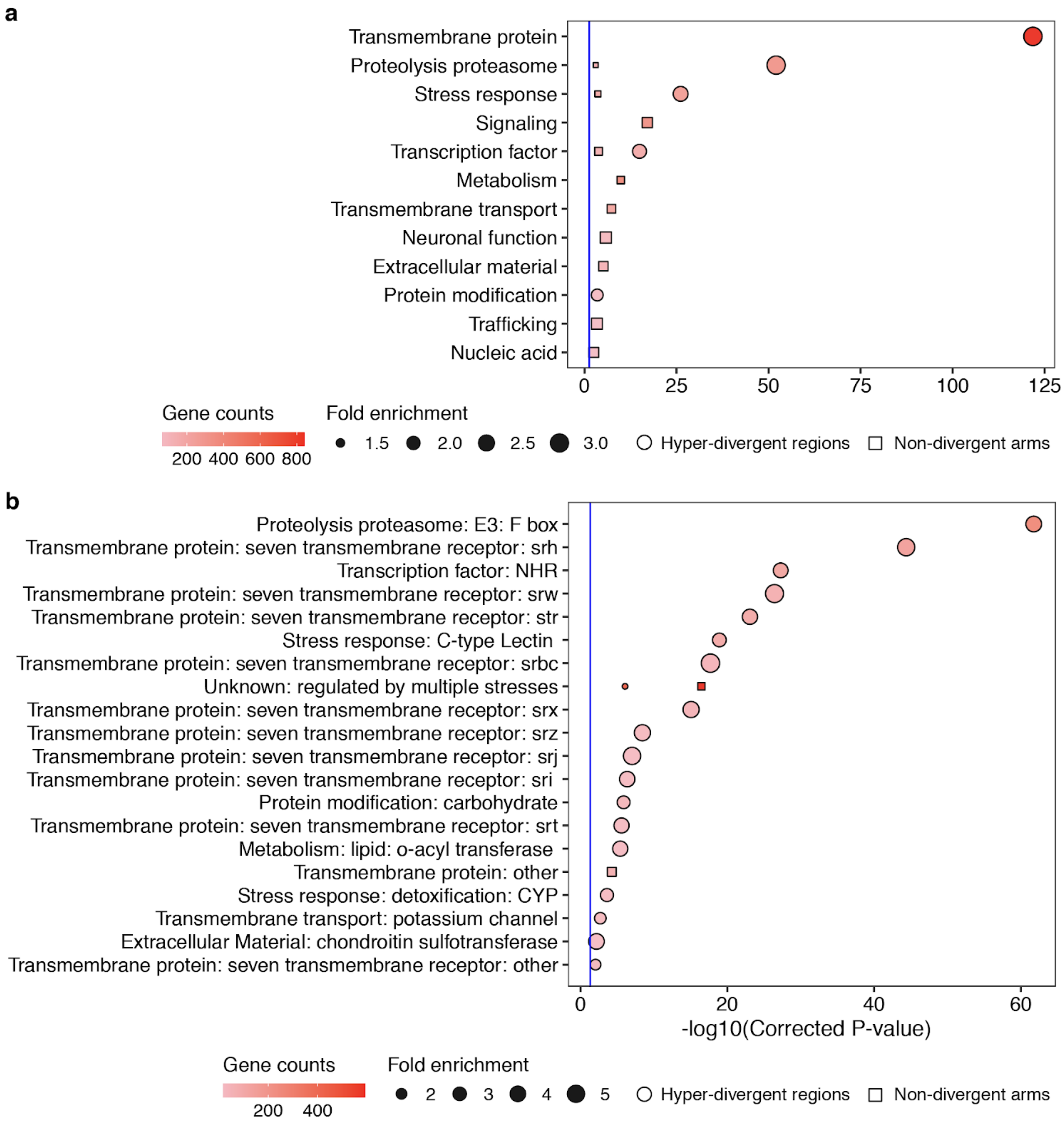


**Supplementary Fig. 6 | Gene-set enrichment (Wormcat) for hyper-divergent regions**

Gene-set enrichment for non-divergent chromosomal arms (square) and hyper-divergent regions (circle) are shown. Annotations in WormCat category 1 (a) and 3 (b) (see Methods) that are significantly enriched in non-divergent regions, hyper-divergent regions, or both are shown on the y-axis. Bonferroni-corrected significance values for gene-set enrichment analysis are shown on the x-axis. Sizes of squares and circles correspond to the fold enrichment of the annotation, and colors of square and circle correspond to the gene counts of the annotation. The blue line shows the Bonferroni-corrected significance threshold (corrected p-value = 0.05).

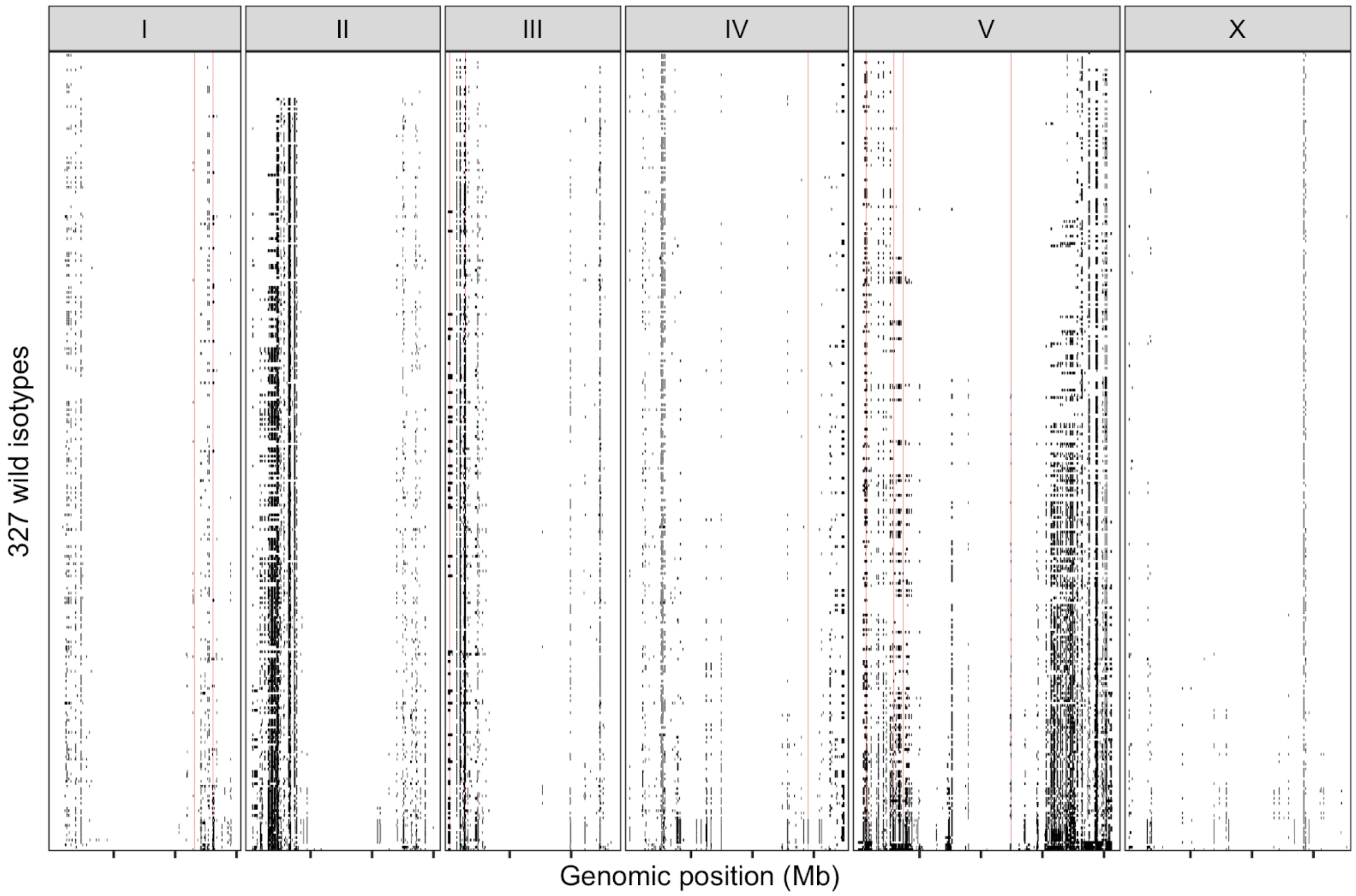


**Supplementary Fig. 7 | Co-localization of *pals* genes and hyper-divergent regions**
The genome-wide distribution of *pals* loci (red line) and hyper-divergent regions across 327 non-reference wild *C. elegans* isotypes is shown. Each row is one of the 327 isotypes, ordered by the total amount of genome covered by hyper-divergent regions (black). The genomic position in Mb is plotted on the x-axis, and each tick represents 5 Mb of the chromosome.

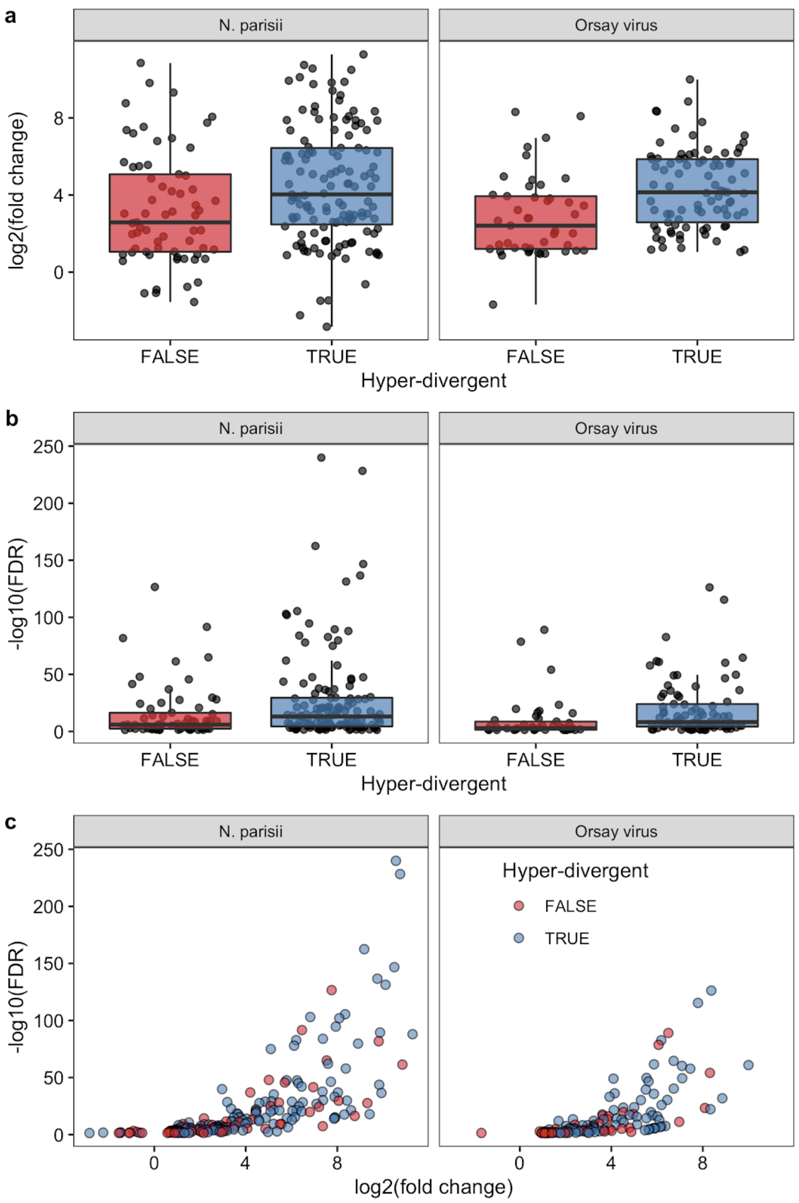


**Supplementary Fig. 8 | Induction of genes in hyper-divergent regions in responses to natural pathogens**

(a) Tukey box plots of fold changes in gene expression level upon natural pathogen treatment are shown. Each point corresponds to a differentially expressed gene (DEG) upon treatment of *N. parisii* (left, n = 196) and Orsay virus (right, n = 130), which are grouped by their hyper-divergent status. Gene expression changes ($log_2$(fold change) in pathogen response are shown on the y-axis.

(b) Tukey box plots of significance in gene expression changes upon natural pathogen treatment are shown with data points plotted behind. Each point corresponds to DEGs upon treatment of *N. parisii* (left, n = 196) and Orsay virus (right, n = 130), which are grouped by their location. The significance of differences in pathogen response are shown on the y-axis. (a,b) The horizontal line in the middle of the box is the median, and the box denotes the 25th to 75th quantiles of the data. The vertical line represents the 1.5x interquartile range.

(c) A scatter plot for the gene expression changes and significance of DEGs in (a) and (b). Each point corresponds to differentially expressed genes (DEGs) upon treatment of *N. parisii* (left, n = 196) and Orsay virus (right, n = 130), which are colored by their location (blue: within hyper-divergent regions, red: within the rest of the genome). Gene expression changes in pathogen response are shown on the x-axis, and significance of differences are shown on the y-axis.

**Supplementary Table 1 | Sampling information for 609 wild strains**
The sampling location and dates for all 611 *C. elegans* strains. The 328 reference isotypes are listed in the second column named “Isotype”. The GPS coordinates are in the third and fourth columns, the person who sampled the strain is listed in the fifth column, and the isolation date is in the sixth column.

| Strain | Isotype | Latitude | Longitude | Sampled by | isolation date |
|---|---|---|---|---|---|
| AB1 | AB1 | -34.93 | 138.59 | D. Riddle and A. Bird | 1983 |
| AB4 | CB4858 | -34.93 | 138.59 | D. Riddle and A. Bird | 1983 |
| BRC20067 | BRC20067 | 24.072954 | 121.169736 | J. Wang | 2012-01-13 |
| BRC20113 | BRC20067 | 24.124167 | 121.283056 | C. Lee and N. Chang | 2014-05-09 |
| BRC20231 | MY23 | 23.541517 | 120.908112 | T. Volans | 2014-06-15 |
| BRC20263 | BRC20263 | -33.8674861 | 151.206989 | C. Lee and N. Chang | 2014-08-04 |
| CB4852 | CB4852 | NA | NA | Unknown | pre-1966 |
| CB4854 | CB4854 | 34.189 | -118.131 | C. Johnson | 1973-12-01 |
| CB4856 | CB4856 | 21.33 | -157.86 | L. Hollen | 1972-08-01 |
| CB4932 | CB4932 | 51.02 | -3.1 | P. Grewal | pre-1991 |
| CX11254 | CX11254 | 34.130138 | -118.11484 | A. Sivasundar | 2003-09-01 |
| CX11262 | CX11262 | 34.130138 | -118.11484 | A. Sivasundar | 2003-09-01 |
| CX11264 | CX11264 | 34.130138 | -118.11484 | A. Sivasundar | 2003-09-01 |
| CX11271 | CX11271 | 34.13712 | -118.12532 | A. Sivasundar | 2003-09-01 |
| CX11276 | CX11276 | 34.20111 | -118.21198 | A. Sivasundar | 2003-09-01 |
| CX11285 | CX11285 | 34.14331 | -118.05496 | A. Sivasundar | 2003-09-01 |
| CX11292 | CX11292 | 34.13531 | -118.30582 | A. Sivasundar | 2004-02-01 |
| CX11307 | CX11307 | 34.12946 | -118.10987 | A. Sivasundar | 2013-09-03 |
| CX11314 | CX11314 | 34.12946 | -118.10987 | A. Sivasundar | 2003-09-01 |
| CX11315 | CX11315 | 34.12946 | -118.10987 | A. Sivasundar | 2013-09-03 |
| DL200 | DL200 | 9.03 | 38.74 | D. Denver | 2007-12-01 |
| DL226 | DL226 | 44.5633 | -123.2821 | C. Hilburn | 2008 |
| DL238 | DL238 | 19.11 | -155.81 | J. Knapp | 2008-07-15 |
| ECA189 | ECA189 | 19.423889 | -155.2225 | T. Blanche | 2014-07-09 |
| ECA190 | ECA189 | 19.423889 | -155.2225 | T. Blanche | 2014-07-09 |
| ECA191 | ECA191 | 19.424167 | -155.22194 | T. Blanche | 2014-07-09 |
| ECA192 | ECA189 | 19.424444 | -155.22194 | T. Blanche | 2014-07-09 |
| ECA193 | ECA189 | 19.424444 | -155.22194 | T. Blanche | 2014-07-09 |
| ECA243 | CB4851 | 44.85 | 0.48 | V. Nigon | pre-1949 |
| ECA245 | CB4853 | 34.171 | -118.131 | C. Johnson | 1974-05-01 |
| ECA246 | CB4853 | 34.189 | -118.131 | C. Johnson | 1974-05-01 |
| ECA248 | CB4855 | 37.44 | -122.14 | T. Doniach | 1982 |
| ECA249 | CB4857 | 34.096 | -117.719 | E. Hedgecock | 1972-11-01 |

| | | | | | |
|---|---|---|---|---|---|
| ECA250 | CB4857 | 34.096 | -117.719 | E. Hedgecock | 1972-11-01 |
| ECA251 | CB4858 | 34.1 | -118.1 | E. Hedgecock | 1973-06-01 |
| ECA259 | PB306 | NA | NA | S. Baird | 1998-11-28 |
| ECA347 | ECA347 | 19.438889 | -155.30361 | C. Andersen and M. Andersen | 2016-02-07 |
| ECA348 | ECA348 | 37.775334 | -122.25389 | S. Zdraljevic | 2015-12-30 |
| ECA349 | ECA349 | 32.765758 | -117.23674 | E. Andersen | 2016-02-09 |
| ECA350 | ECA349 | 32.765353 | -117.23544 | E. Andersen | 2016-02-09 |
| ECA36 | ECA36 | -36.893333 | 174.745529 | I. Ly | 2013-07-27 |
| ECA363 | ECA363 | 20.723611 | -156.30444 | C. Andersen and M. Andersen | 2016-02-19 |
| ECA369 | ECA369 | 21.175556 | -157.00639 | C. Andersen and M. Andersen | 2016-02-20 |
| ECA372 | ECA372 | 22.121667 | -159.66444 | C. Andersen and M. Andersen | 2016-02-27 |
| ECA393 | ECA189 | 19.449831 | -155.23549 | G. Chavez | 2016-05-17 |
| ECA394 | ECA189 | 19.449831 | -155.23549 | G. Chavez | 2016-05-17 |
| ECA395 | ECA189 | 19.449831 | -155.23549 | G. Chavez | 2016-05-17 |
| ECA396 | ECA396 | 19.449831 | -155.23549 | G. Chavez | 2016-05-17 |
| ECA397 | ECA396 | 19.449831 | -155.23549 | G. Chavez | 2016-05-17 |
| ECA398 | ECA396 | 19.449831 | -155.23549 | G. Chavez | 2016-05-17 |
| ECA399 | ECA396 | 19.449831 | -155.23549 | G. Chavez | 2016-05-17 |
| ECA551 | JU311 | 44.42 | 4.4 | M.-A. Félix | 2002-09-08 |
| ECA552 | MY2573 | 43.8658 | 5.0628 | M.-A. Félix | 2013-11-17 |
| ECA571 | ED3073 | -1.05 | 36.39 | E. Dolgin | 2006-05-17 |
| ECA572 | ED3073 | -1.05 | 36.39 | E. Dolgin | 2006-05-17 |
| ECA589 | JU1395 | 47.2199 | 0.04619 | M.-A. Félix | 2008-03-01 |
| ECA592 | ECA592 | 34.417128 | -119.64092 | E. Andersen | 2017-03-07 |
| ECA593 | ECA593 | 34.418632 | -119.64216 | E. Andersen | 2017-03-07 |
| ECA594 | ECA594 | 34.417532 | -119.64067 | E. Andersen | 2017-03-07 |
| ECA615 | JU1581 | 48.7015 | 2.1725 | M.-A. Félix | 2008-09-08 |
| ECA616 | JU1530 | 48.7015 | 2.1725 | M.-A. Félix | 2008-09-09 |
| ECA640 | ECA640 | 34.065378 | -118.44116 | E. Andersen | 2017-06-22 |
| ECA701 | ECA701 | 22.123283 | -159.65643 | D. Cook | 2017-08-11 |
| ECA702 | ECA701 | 22.123283 | -159.65643 | D. Cook | 2017-08-11 |
| ECA703 | ECA703 | 20.038831 | -155.43892 | S. Zdraljevic | 2017-08-13 |
| ECA704 | ECA812 | 20.039783 | -155.44152 | D. Lee | 2017-08-13 |
| ECA705 | ECA705 | 20.038831 | -155.43892 | S. Zdraljevic | 2017-08-13 |
| ECA706 | ECA706 | 20.038831 | -155.43892 | S. Zdraljevic | 2017-08-13 |
| ECA707 | ECA778 | 20.039783 | -155.44152 | D. Lee | 2017-08-13 |
| ECA708 | ECA778 | 20.039783 | -155.44152 | D. Lee | 2017-08-13 |

| | | | | | |
|---|---|---|---|---|---|
| ECA709 | ECA778 | 20.039783 | -155.44152 | D. Lee | 2017-08-13 |
| ECA710 | ECA710 | 20.039656 | -155.4381 | D. Lee | 2017-08-13 |
| ECA711 | ECA712 | 19.423864 | -155.22248 | S. Zdraljevic | 2017-08-14 |
| ECA712 | ECA712 | 19.423864 | -155.22248 | S. Zdraljevic | 2017-08-14 |
| ECA713 | ECA712 | 19.423864 | -155.22248 | S. Zdraljevic | 2017-08-14 |
| ECA714 | ECA712 | 19.423864 | -155.22248 | S. Zdraljevic | 2017-08-14 |
| ECA715 | ECA712 | 19.423864 | -155.22248 | S. Zdraljevic | 2017-08-14 |
| ECA716 | ECA807 | 20.039844 | -155.44141 | D. Lee | 2017-08-13 |
| ECA717 | ECA807 | 20.039844 | -155.44141 | D. Lee | 2017-08-13 |
| ECA718 | ECA807 | 20.039844 | -155.44141 | D. Lee | 2017-08-13 |
| ECA719 | ECA812 | 20.039858 | -155.44142 | S. Zdraljevic | 2017-08-13 |
| ECA720 | ECA807 | 20.039858 | -155.44142 | S. Zdraljevic | 2017-08-13 |
| ECA721 | ECA778 | 20.039858 | -155.44142 | S. Zdraljevic | 2017-08-13 |
| ECA722 | ECA722 | 19.423975 | -155.22263 | S. Zdraljevic | 2017-08-14 |
| ECA723 | ECA723 | 19.424272 | -155.22166 | D. Lee | 2017-08-14 |
| ECA724 | ECA724 | 19.439144 | -155.30383 | S. Zdraljevic | 2017-08-14 |
| ECA725 | ECA724 | 19.439144 | -155.30383 | S. Zdraljevic | 2017-08-14 |
| ECA726 | ECA724 | 19.439144 | -155.30383 | S. Zdraljevic | 2017-08-14 |
| ECA727 | ECA724 | 19.439144 | -155.30383 | S. Zdraljevic | 2017-08-14 |
| ECA728 | ECA723 | 19.424272 | -155.22166 | D. Lee | 2017-08-14 |
| ECA729 | ECA723 | 19.424272 | -155.22166 | D. Lee | 2017-08-14 |
| ECA730 | ECA730 | 19.423869 | -155.22248 | S. Zdraljevic | 2017-08-14 |
| ECA731 | ECA733 | 19.115886 | -155.81839 | S. Zdraljevic | 2017-08-15 |
| ECA732 | ECA732 | 19.115403 | -155.81845 | E. Andersen | 2017-08-15 |
| ECA733 | ECA733 | 19.115886 | -155.81839 | S. Zdraljevic | 2017-08-15 |
| ECA734 | ECA738 | 21.13927631 | -156.97406 | S. Brady | 2017-08-12 |
| ECA735 | ECA738 | 21.13927631 | -156.97406 | S. Brady | 2017-08-12 |
| ECA736 | ECA738 | 21.13927631 | -156.97406 | S. Brady | 2017-08-12 |
| ECA737 | ECA738 | 21.13927631 | -156.97406 | S. Brady | 2017-08-12 |
| ECA738 | ECA738 | 21.13927631 | -156.97406 | S. Brady | 2017-08-12 |
| ECA739 | ECA760 | 20.039869 | -155.44162 | D. Cook | 2017-08-17 |
| ECA740 | ECA740 | 20.670022 | -156.33919 | B. Rodriguez | 2017-08-14 |
| ECA741 | ECA741 | 20.6762 | -156.33832 | B. Rodriguez | 2017-08-14 |
| ECA742 | ECA742 | 20.80569 | -156.2781 | K. Evans | 2017-08-16 |
| ECA743 | ECA743 | 20.741871 | -156.32317 | K. Evans | 2017-08-16 |
| ECA744 | ECA744 | 19.719231 | -155.94957 | S. Zdraljevic | 2017-08-15 |
| ECA745 | ECA745 | 19.700464 | -155.95584 | E. Andersen | 2017-08-15 |

| | | | | | |
|---|---|---|---|---|---|
| ECA746 | ECA746 | 19.115292 | -155.81934 | S. Zdraljevic | 2017-08-15 |
| ECA747 | ECA745 | 19.700464 | -155.95584 | E. Andersen | 2017-08-15 |
| ECA748 | ECA812 | 20.039847 | -155.44149 | D. Cook | 2017-08-17 |
| ECA749 | ECA760 | 20.039875 | -155.44161 | D. Cook | 2017-08-17 |
| ECA750 | ECA812 | 20.039847 | -155.44149 | D. Cook | 2017-08-17 |
| ECA751 | ECA812 | 20.039847 | -155.44149 | D. Cook | 2017-08-17 |
| ECA752 | ECA760 | 20.039847 | -155.44149 | D. Cook | 2017-08-17 |
| ECA753 | ECA760 | 20.039847 | -155.44149 | D. Cook | 2017-08-17 |
| ECA754 | ECA760 | 20.039869 | -155.44146 | D. Lee | 2017-08-17 |
| ECA755 | ECA812 | 20.039869 | -155.44146 | D. Lee | 2017-08-17 |
| ECA756 | ECA778 | 20.039822 | -155.44153 | D. Cook | 2017-08-17 |
| ECA757 | ECA778 | 20.039822 | -155.44153 | D. Cook | 2017-08-17 |
| ECA758 | ECA760 | 20.039886 | -155.44153 | D. Lee | 2017-08-17 |
| ECA759 | ECA812 | 20.039886 | -155.44153 | D. Lee | 2017-08-17 |
| ECA760 | ECA760 | 20.039869 | -155.44162 | D. Cook | 2017-08-17 |
| ECA761 | ECA812 | 20.039847 | -155.44149 | D. Cook | 2017-08-17 |
| ECA762 | ECA812 | 20.039822 | -155.44153 | D. Cook | 2017-08-17 |
| ECA763 | ECA778 | 20.039878 | -155.44161 | D. Cook | 2017-08-17 |
| ECA764 | ECA778 | 20.039878 | -155.44161 | D. Cook | 2017-08-17 |
| ECA765 | ECA778 | 20.039878 | -155.44161 | D. Cook | 2017-08-17 |
| ECA766 | ECA778 | 20.039878 | -155.44161 | D. Cook | 2017-08-17 |
| ECA767 | ECA778 | 20.039878 | -155.44161 | D. Cook | 2017-08-17 |
| ECA768 | ECA768 | 20.039914 | -155.44156 | D. Cook | 2017-08-17 |
| ECA769 | ECA760 | 20.039869 | -155.44162 | D. Cook | 2017-08-17 |
| ECA770 | ECA807 | 20.039869 | -155.44153 | D. Lee | 2017-08-17 |
| ECA771 | ECA812 | 20.039864 | -155.44164 | D. Cook | 2017-08-17 |
| ECA772 | ECA768 | 20.039908 | -155.4415 | D. Lee | 2017-08-17 |
| ECA773 | ECA812 | 20.039864 | -155.44164 | D. Cook | 2017-08-17 |
| ECA774 | ECA812 | 20.039869 | -155.44153 | D. Lee | 2017-08-17 |
| ECA775 | ECA812 | 20.039869 | -155.44153 | D. Lee | 2017-08-17 |
| ECA776 | ECA812 | 20.039869 | -155.44153 | D. Lee | 2017-08-17 |
| ECA777 | ECA777 | 20.039908 | -155.4415 | D. Lee | 2017-08-17 |
| ECA778 | ECA778 | 20.039908 | -155.4415 | D. Lee | 2017-08-17 |
| ECA779 | ECA760 | 20.039858 | -155.44161 | D. Cook | 2017-08-17 |
| ECA780 | ECA760 | 20.039858 | -155.44161 | D. Cook | 2017-08-17 |
| ECA781 | ECA778 | 20.039858 | -155.44161 | D. Cook | 2017-08-17 |
| ECA782 | ECA812 | 20.039875 | -155.44161 | D. Cook | 2017-08-17 |

| | | | | | |
|---|---|---|---|---|---|
| ECA783 | ECA812 | 20.039783 | -155.44148 | D. Lee | 2017-08-17 |
| ECA784 | ECA760 | 20.039783 | -155.44148 | D. Lee | 2017-08-17 |
| ECA785 | ECA760 | 20.039783 | -155.44148 | D. Lee | 2017-08-17 |
| ECA786 | ECA812 | 20.039875 | -155.44161 | D. Cook | 2017-08-17 |
| ECA787 | ECA812 | 20.039783 | -155.44148 | D. Lee | 2017-08-17 |
| ECA807 | ECA807 | 20.039869 | -155.44153 | D. Lee | 2017-08-17 |
| ECA808 | ECA760 | 20.039858 | -155.44161 | D. Cook | 2017-08-17 |
| ECA809 | ECA812 | 20.039783 | -155.44148 | D. Lee | 2017-08-17 |
| ECA810 | ECA778 | 20.039897 | -155.44153 | D. Lee | 2017-08-17 |
| ECA811 | ECA778 | 20.039897 | -155.44153 | D. Lee | 2017-08-17 |
| ECA812 | ECA812 | 20.039917 | -155.44154 | D. Lee | 2017-08-17 |
| ECA813 | ECA812 | 20.039917 | -155.44154 | D. Lee | 2017-08-17 |
| ECA822 | ECA705 | 20.038831 | -155.43892 | S. Zdraljevic | 2017-08-13 |
| ECA922 | ECA923 | 21.290067 | -157.70792 | N. Singh | 2017-12-01 |
| ECA923 | ECA923 | 21.290067 | -157.70792 | N. Singh | 2017-12-01 |
| ECA924 | ECA923 | 21.290067 | -157.70792 | N. Singh | 2017-12-01 |
| ECA925 | ECA923 | 21.290067 | -157.70792 | N. Singh | 2017-12-01 |
| ECA926 | ECA928 | 21.290067 | -157.70792 | N. Singh | 2017-12-01 |
| ECA927 | ECA923 | 21.290067 | -157.70792 | N. Singh | 2017-12-01 |
| ECA928 | ECA928 | 21.290067 | -157.70792 | N. Singh | 2017-12-01 |
| ECA930 | ECA930 | 31.778363 | 35.197931 | E. Bokman, A. Topper, C. Pritz | 2017-12-14 |
| ED3005 | ED3005 | 55.94 | -3.36 | A. Cutter | 2004-10-25 |
| ED3011 | ED3011 | 55.92 | -3.19 | A. Cutter | 2004-11-11 |
| ED3012 | ED3012 | 55.92 | -3.19 | A. Cutter | 2004-11-26 |
| ED3017 | ED3017 | 55.92 | -3.19 | A. Cutter | 2004-12-03 |
| ED3040 | ED3040 | -26.166667 | 28.016667 | E. Dolgin | 2006-05-05 |
| ED3046 | ED3046 | -33.366667 | 19.316667 | E. Dolgin | 2006-05-05 |
| ED3048 | ED3048 | -33.366667 | 19.316667 | E. Dolgin | 2006-04-02 |
| ED3049 | ED3049 | -33.366667 | 19.316667 | E. Dolgin | 2006-04-02 |
| ED3052 | ED3052 | -33.366667 | 19.316667 | E. Dolgin | 2006-05-05 |
| ED3073 | ED3073 | -1.083333 | 36.65 | E. Dolgin | 2006-05-17 |
| ED3077 | ED3077 | -1.316667 | 36.8 | E. Dolgin | 2006-05-17 |
| EG4347 | EG4347 | 44.04789 | -123.07108 | M. Ailion | 2006-10-09 |
| EG4349 | EG4349 | 40.771467 | -111.87316 | M. Ailion | 2006-10-02 |
| EG4724 | EG4724 | 41.6288 | -8.3476 | M. Ailion | 2007-03-28 |
| EG4725 | EG4725 | 41.6288 | -8.3476 | M. Ailion | 2007-03-28 |
| EG4946 | EG4946 | 40.72596 | -111.82184 | G. Hollopeter | 2007-09-27 |

| | | | | | |
|---|---|---|---|---|---|
| GXW1 | GXW1 | 30.542889 | 114.419828 | G. Wang | 2010-11-05 |
| JT11398 | JT11398 | 47.763944 | -122.27548 | J. Kemner | 2003-12-01 |
| JU1088 | JU1088 | 34.7613 | 138.0149 | M.-A. Félix | 2007-03-14 |
| JU1172 | JU1172 | -36.87 | -73.04 | M.-A. Félix | 2007-04-01 |
| JU1200 | JU1200 | 55.577 | -4.6 | T. Page | 2007-08-01 |
| JU1212 | JU1212 | 48.71 | -3.81 | M.-A. Félix | 2007-09-24 |
| JU1213 | JU1213 | 48.71 | -3.81 | M.-A. Félix | 2007-09-24 |
| JU1242 | JU1242 | 49.1269 | 1.9595 | M.-A. Félix | 2007-10-14 |
| JU1246 | JU1246 | 49.12618 | 1.96152 | M.-A. Félix | 2007-10-14 |
| JU1249 | JU1249 | 49.126 | 1.951 | M.-A. Félix | 2007-10-14 |
| JU1395 | JU1395 | 47.2199 | 0.04619 | M.-A. Félix | 2008-03-01 |
| JU1400 | JU1400 | 37.3845 | -5.988 | M.-A. Félix | 2008-03-29 |
| JU1409 | JU1409 | 37.468 | -5.637 | M.-A. Félix | 2008-03-31 |
| JU1440 | JU1440 | 41.41307 | 2.15231 | M.-A. Félix | 2008-06-09 |
| JU1491 | JU1491 | 46.63 | 1.06 | M.-A. Félix | 2008-08-17 |
| JU1516 | JU1581 | 48.7015 | 2.1725 | M.-A. Félix | 2008-09-09 |
| JU1530 | JU1530 | 48.7015 | 2.1725 | M.-A. Félix | 2008-09-09 |
| JU1543 | JU1543 | 48.7015 | 2.1725 | M.-A. Félix | 2008-09-08 |
| JU1568 | JU1568 | 48.8092 | 2.3862 | M.-A. Félix | 2008-10-05 |
| JU1580 | JU1580 | 48.7015 | 2.1725 | M.-A. Félix | 2008-10-06 |
| JU1581 | JU1581 | 48.7015 | 2.1725 | M.-A. Félix | 2008-10-23 |
| JU1586 | JU1586 | 46.63 | 1.06 | M.-A. Félix | 2008-11-03 |
| JU1652 | JU1652 | -34.86 | -56.19 | R. Giordano | 2009 |
| JU1656 | JU1213 | 48.7049 | -3.7941 | M.-A. Félix | 2009-07-14 |
| JU1666 | JU1666 | 48.7049 | -3.7941 | M.-A. Félix | 2009-07-14 |
| JU1762 | JU1213 | 48.695548 | -3.790033 | M.-A. Félix | 2009-08-16 |
| JU1770 | JU1213 | 48.695548 | -3.790033 | M.-A. Félix | 2009-08-16 |
| JU1792 | JU1792 | 46.63278 | 1.064289 | M.-A. Félix | 2009-09-27 |
| JU1793 | JU1793 | 49.12604 | 1.95114 | M.-A. Félix | 2009-10-10 |
| JU1807 | JU1581 | 48.662134 | 2.2131 | M.-A. Félix | 2008-10-23 |
| JU1808 | JU1808 | 48.81257 | 2.395372 | M.-A. Félix | 2009-10-14 |
| JU1896 | JU1896 | 37.999722 | 23.749673 | M. Barkoulas | 2010-01-02 |
| JU1920 | JU1793 | 49.12604 | 1.95114 | M.-A. Félix | 2009-10-25 |
| JU1922 | JU1793 | 49.12604 | 1.95114 | M.-A. Félix | 2009-10-25 |
| JU1924 | JU2600 | 49.12604 | 1.95114 | M.-A. Félix | 2009-10-25 |
| JU1929 | JU1793 | 49.12604 | 1.95114 | M.-A. Félix | 2009-10-25 |
| JU1934 | JU1934 | 49.12604 | 1.95114 | M.-A. Félix | 2009-10-25 |

| | | | | | |
|---|---|---|---|---|---|
| JU1960 | CB4858 | 34.18973 | -118.13132 | J. DeModena | 2008-11-30 |
| JU2001 | JU2001 | -21.1 | 55.5 | T. Bélicard | 2010-09-01 |
| JU2007 | JU2007 | 50.7608 | -1.32413 | M. Barkoulas | 2010-10-16 |
| JU2016 | JU2016 | 37.757593 | -122.46318 | C. Nelson | 2010-11-01 |
| JU2017 | JU2017 | 37.757593 | -122.46318 | C. Nelson | 2010-11-01 |
| JU2106 | JU2106 | 43.06 | 0.24 | M.-A. Félix | 2011-07-10 |
| JU2131 | JU2131 | 48.0497 | -4.705 | M.-A. Félix | 2011-08-09 |
| JU2139 | JU1793 | 46.63278 | 1.064289 | M.-A. Félix | 2011-10-23 |
| JU2141 | JU2141 | 46.6293 | 1.0512 | M.-A. Félix | 2011-10-23 |
| JU2151 | JU2600 | 49.121 | 1.951 | M.-A. Félix | 2011-10-31 |
| JU2234 | JU2234 | 47.227 | -1.583 | L. Frézal | 2012-11-11 |
| JU2250 | JU2250 | 48.285278 | -1.891389 | L. Frézal | 2012-11-15 |
| JU2257 | JU2257 | 46.6 | 1.1 | M.-A. Félix | 2013-08-15 |
| JU2287 | JU2257 | 46.6 | 1.1 | M.-A. Félix | 2013-08-15 |
| JU2316 | JU2316 | 38.7175 | -9.1486 | M.-A. Félix | 2013-08-25 |
| JU2460 | JU792 | 48.6934 | -3.9251 | M.-A. Félix | 2012-12-28 |
| JU2464 | JU2464 | -13.155 | -72.525 | L. Lokmane | 2013-01 |
| JU2466 | JU2466 | -13.257 | -72.266 | L. Lokmane | 2013-01 |
| JU2467 | JU2464 | -13.155 | -72.525 | L. Lokmane | 2013-01 |
| JU2468 | JU2464 | -13.155 | -72.525 | L. Lokmane | 2013-01 |
| JU2478 | JU2478 | 50.85 | 4.351 | D. Thieffry | 2013-02-01 |
| JU2513 | JU2513 | 38.26016 | 22.0736 | M.-A. Félix | 2013-05-07 |
| JU2519 | JU2519 | 38.7175 | -9.1486 | M.-A. Félix | 2013-08-25 |
| JU2522 | JU2522 | 38.7175 | -9.1486 | M.-A. Félix | 2013-08-25 |
| JU2526 | JU2526 | 38.7175 | -9.1486 | M.-A. Félix | 2013-08-25 |
| JU2527 | JU792 | 47.8797 | -3.5999 | F. Besnard | 2013-08-19 |
| JU2534 | JU2534 | 49.357921 | 0.097087 | S. Marsh | 2013-06-09 |
| JU2565 | JU2565 | 44.958394 | 1.066213 | T. Bélicard | 2013-10-05 |
| JU2566 | JU2566 | 48.83886 | 2.44599 | M.-A. Félix | 2013-10-02 |
| JU2570 | JU2570 | 46.2657 | -0.2685 | F. Besnard | 2013-11-13 |
| JU2572 | JU2572 | 48.81257 | 2.395372 | S. Marsh | 2013-10-01 |
| JU2575 | JU2575 | 49.0857 | 3.0707 | M.-A. Félix and S. Marsh | 2013-10-26 |
| JU2576 | JU2576 | 49.0861 | 3.072 | M.-A. Félix and S. Marsh | 2013-10-26 |
| JU2578 | JU2578 | 48.87613 | 2.4315 | M.-A. Félix | 2013-10-27 |
| JU258 | JU258 | 32.73 | -16.89 | M.-A. Félix | 2001-10-01 |
| JU2581 | JU2581 | 46.6 | 1.1 | M.-A. Félix | 2013-11-03 |
| JU2586 | JU2586 | 48.897 | 2.159 | A. Zalmanski | 2013-11-11 |

| | | | | | |
|---|---|---|---|---|---|
| JU2587 | JU2587 | 45.026713 | 3.872016 | A. Richaud | 2013-11-10 |
| JU2592 | JU2592 | 48.628558 | -2.403836 | L. Frézal | 2013-11-11 |
| JU2593 | JU2593 | 48.822556 | 2.222334 | O. Bensaude | 2013 |
| JU2600 | JU2600 | 49.1242 | 1.9496 | M.-A. Félix and S. Marsh | 2013-11-20 |
| JU2604 | JU1793 | 49.16477 | 1.86175 | M.-A. Félix and S. Marsh | 2013-11-20 |
| JU2605 | JU1793 | 49.166 | 1.86 | M.-A. Félix and S. Marsh | 2013-11-20 |
| JU2610 | JU2610 | 49.1075 | 1.7915 | M.-A. Félix and S. Marsh | 2013-11-20 |
| JU2619 | JU2619 | 32.998352 | -117.27543 | S. Marsh | 2014-01 |
| JU2800 | JU2800 | 48.79 | 2.331 | M. Casado | 2014-09-01 |
| JU2802 | JU2800 | 48.9067 | 2.14167 | F. Besnard | 2014-09-14 |
| JU2811 | JU2811 | -33.8632 | 151.219453 | S. Marsh | 2014-09-24 |
| JU2825 | JU2825 | 48.8092 | 2.3862 | R. Luallen | 2014-11-14 |
| JU2828 | JU2829 | 46.632 | 1.064 | M.-A. Félix | 2010-10-03 |
| JU2829 | JU2829 | 46.632 | 1.064 | M.-A. Félix | 2010-10-03 |
| JU2830 | JU2829 | 46.632 | 1.064 | M.-A. Félix | 2010-10-03 |
| JU2838 | JU2838 | 20.205961 | -97.98832 | A. Vargas Velazquez | 2015 |
| JU2841 | JU2841 | -40 | 176 | M. Wilson | 2015-03-01 |
| JU2853 | JU2853 | 41.19 | -7.545 | M. Félix and B. Félix | 2015-05-01 |
| JU2860 | JU1200 | 52.19842 | 0.115365 | M.-A. Félix | 2015-06-06 |
| JU2862 | JU2862 | 52.1944 | 0.1264 | M.-A. Félix | 2015-06-06 |
| JU2866 | JU2866 | 34.0751 | -118.4409 | M.-A. Félix | 2015-06-28 |
| JU2878 | JU2878 | 19.29732 | -99.098769 | A. Vargas Velazquez | 2015-07-02 |
| JU2879 | JU2879 | 19.29732 | -99.098769 | A. Vargas Velazquez | 2015-07-02 |
| JU2906 | JU2906 | 43.8023 | 11.2806 | M.-A. Félix | 2015-12-13 |
| JU2907 | JU2907 | 43.8021 | 11.2906 | M.-A. Félix | 2015-12-13 |
| JU2908 | JU2907 | 43.8063 | 11.2839 | M.-A. Félix | 2015-12-13 |
| JU310 | JU310 | 46.63 | 1.06 | M.-A. Félix | 2002-08-25 |
| JU311 | JU311 | 44.42 | 4.4 | M.-A. Félix | 2002-09-08 |
| JU312 | JU311 | 44.42 | 4.4 | M.-A. Félix | 2002-09-08 |
| JU3125 | JU3125 | 38.1127 | 13.3722 | M.-A. Félix | 2016-04-20 |
| JU3127 | JU3127 | 38.11196 | 13.37274 | M.-A. Félix | 2016-04-20 |
| JU3128 | JU3128 | 37.076 | 15.2763 | M.-A. Félix | 2016-04-23 |
| JU3131 | JU323 | 48.7015 | 2.1725 | M.-A. Félix | 2008-10-23 |
| JU3132 | JU3132 | 48.7015 | 2.1725 | M.-A. Félix | 2011-10-10 |
| JU3133 | JU1440 | 48.7015 | 2.1725 | M.-A. Félix | 2011-11-08 |
| JU3134 | JU3134 | 48.7015 | 2.1725 | M.-A. Félix | 2011-11-11 |
| JU3135 | JU3135 | 48.7015 | 2.1725 | M.-A. Félix | 2013-10-29 |

| | | | | | |
|---|---|---|---|---|---|
| JU3136 | MY2453 | 48.7015 | 2.1725 | M.-A. Félix | 2013-11-13 |
| JU3137 | JU3137 | 48.7015 | 2.1725 | M.-A. Félix | 2014-10-20 |
| JU3138 | JU2600 | 49.1269 | 1.9595 | M.-A. Félix | 2011-11-07 |
| JU3139 | JU1666 | 48.71 | -3.81 | M.-A. Félix | 2009-08-09 |
| JU3140 | JU3140 | 48.71 | -3.81 | M.-A. Félix | 2009-08-09 |
| JU3141 | JU3141 | 48.71 | -3.81 | M.-A. Félix | 2009-08-09 |
| JU3142 | JU792 | 48.71 | -3.81 | M.-A. Félix | 2009-08-09 |
| JU3144 | JU3144 | 6.18 | 10.52 | J. David | 2016-05-01 |
| JU315 | JU311 | 44.2 | 4.4 | M.-A. Félix | 2002-09-08 |
| JU3166 | JU3166 | 0.28 | 6.59837 | M.-A. Félix | 2016-09-19 |
| JU3167 | JU3167 | 0.289 | 6.612 | J. David | 2016-09-21 |
| JU3169 | JU3169 | 0.289 | 6.612 | J. David | 2016-09-21 |
| JU3224 | JU3224 | -41.296186 | 174.784857 | J. Ewbank | 2017-04-01 |
| JU3225 | JU3225 | -41.296186 | 174.784857 | N. Pujol | 2017-04-02 |
| JU3226 | JU3226 | -41.29085 | 174.76872 | N. Pujol | 2017-04-03 |
| JU3227 | JU3228 | -38.64844 | 176.08947 | N. Pujol | 2017-04-04 |
| JU3228 | JU3228 | -39.02256 | 175.71458 | N. Pujol | 2017-04-05 |
| JU323 | JU323 | 44.42 | 4.4 | M.-A. Félix | 2002-09-08 |
| JU3271 | JU1793 | 49.11 | 1.93 | M.-A. Félix and E. Troemel | NA |
| JU3280 | JU3280 | 50.07182 | 14.42333 | M.-A. Félix | 2017-09-24 |
| JU3282 | JU3282 | 50.0711 | 14.42028 | M.-A. Félix | 2017-09-24 |
| JU3291 | JU3291 | 48.7679 | 2.70719 | M.-A. Félix | 2017-10-08 |
| JU345 | JU346 | 44.42 | 4.4 | M.-A. Félix | 2002-09-08 |
| JU346 | JU346 | 44.42 | 4.4 | M.-A. Félix | 2002-09-08 |
| JU360 | JU360 | 48.98 | 2.23 | C. Pieau | 2002-09-16 |
| JU363 | JU360 | 48.98 | 2.23 | C. Pieau | 2002-09-16 |
| JU367 | JU367 | 48.98 | 2.23 | C. Pieau | 2002-09-16 |
| JU393 | JU393 | 49.28 | -0.32 | A. Barrière | 2002-09-01 |
| JU394 | JU394 | 49.28 | -0.32 | A. Barrière | 2002-09-01 |
| JU397 | JU397 | 49.28 | -0.32 | A. Barrière | 2002-09-01 |
| JU406 | JU406 | 49.28 | -0.32 | A. Barrière | 2002-12-30 |
| JU440 | JU440 | 48.715 | 1.56 | D. Baïlle | 2003-09-12 |
| JU561 | JU561 | 48.71 | -3.81 | M.-A. Félix | 2004-10-03 |
| JU642 | JU642 | 48.84 | 2.5 | J.-A. Lepesant | 2004-12-14 |
| JU751 | JU751 | 48.84 | 2.5 | J.-A. Lepesant | 2005-06-08 |
| JU774 | JU774 | 38.683 | -9.34 | M.-A. Félix | 2005-07-10 |
| JU775 | JU775 | 38.7175 | -9.1486 | M.-A. Félix | 2005-07-10 |

| | | | | | |
|---|---|---|---|---|---|
| JU778 | JU778 | 38.719 | -9.1491 | M.-A. Félix | 2005-07-10 |
| JU782 | JU782 | 38.7191 | -9.1503 | M.-A. Félix | 2005-07-10 |
| JU792 | JU792 | 43.06 | 0.24 | M.-A. Félix | 2005-08-31 |
| JU830 | JU830 | 48.52 | 9.05 | R. Hong | 2005-09-28 |
| JU847 | JU847 | 48.46 | 7.461 | M.-A. Félix | 2005-10-03 |
| KR314 | KR314 | 49.28 | -123.13 | F. Dill | 1984-05-01 |
| LKC34 | LKC34 | -18 | 46 | V. Stowell | 2005-06-17 |
| MY1 | MY1 | 52.54 | 7.31 | H. Schulenburg | 2002-07-01 |
| MY10 | MY10 | 51.96 | 7.53 | H. Schulenburg | 2002-07-01 |
| MY16 | MY16 | 51.93 | 7.57 | H. Schulenburg | 2002-07-01 |
| MY18 | MY18 | 51.96 | 7.53 | H. Schulenburg | 2002-07-01 |
| MY2001 | MY2453 | 51.950777 | 7.536628 | C. Petersen | 2012-06-20 |
| MY2004 | MY2453 | 51.950777 | 7.536628 | C. Petersen | 2012-06-20 |
| MY2011 | MY2453 | 51.950777 | 7.536628 | C. Petersen | 2012-07-04 |
| MY2014 | MY2453 | 51.950777 | 7.536628 | C. Petersen | 2012-07-04 |
| MY2022 | RC301 | 54.346355 | 10.117737 | C. Petersen | 2012-07-04 |
| MY2024 | RC301 | 54.346355 | 10.117737 | C. Petersen | 2012-07-04 |
| MY2042 | MY920 | 54.346355 | 10.117737 | C. Petersen | 2012-07-18 |
| MY2050 | MY2453 | 51.950777 | 7.536628 | C. Petersen | 2012-07-18 |
| MY2051 | MY2453 | 51.950777 | 7.536628 | C. Petersen | 2012-07-18 |
| MY2054 | MY2453 | 51.950777 | 7.536628 | C. Petersen | 2012-07-18 |
| MY2078 | RC301 | 54.346355 | 10.117737 | C. Petersen | 2012-08-01 |
| MY2097 | MY920 | 54.346355 | 10.117737 | C. Petersen | 2012-08-15 |
| MY2099 | MY795 | 54.346355 | 10.117737 | C. Petersen | 2012-08-15 |
| MY2109 | MY2741 | 54.346355 | 10.117737 | C. Petersen | 2012-08-15 |
| MY2121 | MY2453 | 51.950777 | 7.536628 | C. Petersen | 2012-08-15 |
| MY2137 | MY2453 | 51.950777 | 7.536628 | C. Petersen | 2012-09-17 |
| MY2138 | MY920 | 54.346355 | 10.117737 | C. Petersen | 2012-09-17 |
| MY2142 | MY2573 | 51.950723 | 7.599099 | C. Petersen | 2012-09-17 |
| MY2143 | MY2147 | 54.346355 | 10.117737 | C. Petersen | 2012-09-17 |
| MY2144 | MY2573 | 51.950723 | 7.599099 | C. Petersen | 2012-09-17 |
| MY2147 | MY2147 | 54.346355 | 10.117737 | C. Petersen | 2012-09-17 |
| MY2198 | MY2573 | 51.950723 | 7.599099 | C. Petersen | 2012-09-26 |
| MY2199 | MY2453 | 51.950777 | 7.536628 | C. Petersen | 2012-09-26 |
| MY2208 | MY2573 | 54.346355 | 10.117737 | C. Petersen | 2012-09-26 |
| MY2212 | MY2212 | 54.346355 | 10.117737 | C. Petersen | 2012-09-26 |
| MY2224 | MY2573 | 54.346355 | 10.117737 | C. Petersen | 2012-09-26 |

| | | | | | |
|---|---|---|---|---|---|
| MY2239 | MY2573 | 54.346355 | 10.117737 | C. Petersen | 2012-09-26 |
| MY2282 | MY2573 | 51.950723 | 7.599099 | C. Petersen | 2012-10-10 |
| MY2288 | MY2453 | 51.950777 | 7.536628 | C. Petersen | 2012-10-10 |
| MY2291 | RC301 | 54.346355 | 10.117737 | C. Petersen | 2012-10-10 |
| MY2294 | MY2573 | 51.950723 | 7.599099 | C. Petersen | 2012-10-10 |
| MY23 | MY23 | 51.96 | 7.53 | H. Schulenburg | 2002-07-01 |
| MY2338 | MY2573 | 51.950723 | 7.599099 | C. Petersen | 2012-10-24 |
| MY2339 | MY2453 | 51.950777 | 7.536628 | C. Petersen | 2012-10-24 |
| MY2344 | MY2535 | 51.950723 | 7.599099 | C. Petersen | 2012-10-24 |
| MY2347 | MY920 | 54.346355 | 10.117737 | C. Petersen | 2012-10-24 |
| MY2373 | MY920 | 54.346355 | 10.117737 | C. Petersen | 2012-10-24 |
| MY2406 | MY920 | 54.346355 | 10.117737 | C. Petersen | 2012-10-24 |
| MY2434 | MY2573 | 51.950723 | 7.599099 | C. Petersen | 2012-11-07 |
| MY2443 | MY2573 | 51.950723 | 7.599099 | C. Petersen | 2012-11-07 |
| MY2453 | MY2453 | 51.950777 | 7.536628 | C. Petersen | 2012-11-07 |
| MY2479 | MY920 | 54.346355 | 10.117737 | C. Petersen | 2012-11-21 |
| MY2481 | JU311 | 54.346355 | 10.117737 | C. Petersen | 2012-11-21 |
| MY2491 | MY2573 | 51.950723 | 7.599099 | C. Petersen | 2012-11-21 |
| MY2502 | MY2453 | 51.950777 | 7.536628 | C. Petersen | 2012-11-21 |
| MY2530 | MY2530 | 54.346355 | 10.117737 | C. Petersen | 2012-12-05 |
| MY2532 | MY2535 | 54.346355 | 10.117737 | C. Petersen | 2012-12-05 |
| MY2535 | MY2535 | 51.950723 | 7.599099 | C. Petersen | 2012-12-05 |
| MY2541 | MY2453 | 51.950777 | 7.536628 | C. Petersen | 2012-12-05 |
| MY2573 | MY2573 | 51.950723 | 7.599099 | C. Petersen | 2012-12-10 |
| MY2579 | MY2453 | 51.950777 | 7.536628 | C. Petersen | 2012-12-10 |
| MY2585 | MY2585 | 54.346355 | 10.117737 | C. Petersen | 2012-12-10 |
| MY2622 | MY2453 | 54.3491 | 10.11505 | R. Hermann | 2013-08-09 |
| MY2623 | MY2585 | 54.3491 | 10.11505 | C. Petersen | 2013-08-09 |
| MY2630 | RC301 | 54.3491 | 10.11505 | R. Hermann | 2013-08-09 |
| MY2635 | MY2713 | 54.3491 | 10.11505 | R. Hermann | 2013-08-23 |
| MY2636 | RC301 | 54.3491 | 10.11505 | R. Hermann | 2013-08-23 |
| MY2640 | RC301 | 54.3491 | 10.11505 | R. Hermann | 2013-08-23 |
| MY2679 | MY2713 | 54.3491 | 10.11505 | R. Hermann | 2013-09-04 |
| MY2681 | RC301 | 54.3491 | 10.11505 | C. Petersen | 2013-09-04 |
| MY2684 | RC301 | 54.3491 | 10.11505 | M. Barg | 2013-08-19 |
| MY2685 | MY2693 | 54.3491 | 10.11505 | M. Barg | 2013-08-19 |
| MY2688 | MY920 | 54.3491 | 10.11505 | M. Barg | 2013-08-21 |

| | | | | | |
|---|---|---|---|---|---|
| MY2689 | MY2713 | 54.3491 | 10.11505 | M. Barg | 2013-08-21 |
| MY2691 | RC301 | 54.3491 | 10.11505 | R. Hermann | 2013-07-31 |
| MY2692 | MY2741 | 54.3491 | 10.11505 | R. Hermann | 2013-08-16 |
| MY2693 | MY2693 | 54.3491 | 10.11505 | R. Hermann | 2013-08-15 |
| MY2713 | MY2713 | 54.3491 | 10.11505 | M. Barg | 2013-09-03 |
| MY2719 | MY2693 | 54.3491 | 10.11505 | M. Barg | 2013-09-05 |
| MY2741 | MY2741 | 54.3491 | 10.11505 | C. Petersen | 2014-09-01 |
| MY508 | MY2453 | 51.950777 | 7.536628 | C. Petersen | 2011-07-08 |
| MY518 | MY518 | 51.964154 | 7.611719 | C. Petersen | 2011-07-08 |
| MY524 | MY2453 | 51.950777 | 7.536628 | C. Petersen | 2011-11-28 |
| MY538 | MY795 | 54.346355 | 10.117737 | C. Petersen | 2011-08-25 |
| MY559 | MY2573 | 51.950723 | 7.599099 | C. Petersen | 2011-09-09 |
| MY561 | MY2453 | 51.950777 | 7.536628 | C. Petersen | 2011-09-08 |
| MY564 | MY2453 | 51.950777 | 7.536628 | C. Petersen | 2011-09-08 |
| MY570 | MY2453 | 51.950777 | 7.536628 | C. Petersen | 2011-09-08 |
| MY579 | MY2573 | 51.950723 | 7.599099 | C. Petersen | 2011-09-09 |
| MY589 | MY920 | 54.346355 | 10.117737 | C. Petersen | 2011-08-25 |
| MY673 | MY795 | 54.346355 | 10.117737 | C. Petersen | 2011-09-27 |
| MY679 | MY679 | 54.346355 | 10.117737 | C. Petersen | 2011-09-27 |
| MY684 | RC301 | 54.346355 | 10.117737 | C. Petersen | 2011-09-27 |
| MY710 | MY2573 | 51.950723 | 7.599099 | C. Petersen | 2011-10-12 |
| MY713 | MY2453 | 51.950777 | 7.536628 | C. Petersen | 2011-10-12 |
| MY741 | MY920 | 54.346355 | 10.117737 | C. Petersen | 2011-10-10 |
| MY772 | MY772 | 54.346355 | 10.117737 | C. Petersen | 2011-09-27 |
| MY792 | MY2453 | 51.950777 | 7.536628 | C. Petersen | 2011-10-24 |
| MY795 | MY795 | 54.346355 | 10.117737 | C. Petersen | 2011-10-26 |
| MY803 | MY2713 | 54.346355 | 10.117737 | C. Petersen | 2011-11-03 |
| MY804 | MY2573 | 51.950723 | 7.599099 | C. Petersen | 2011-11-02 |
| MY819 | MY2453 | 51.950777 | 7.536628 | C. Petersen | 2011-11-02 |
| MY864 | MY2453 | 51.950723 | 7.599099 | C. Petersen | 2011-11-14 |
| MY881 | MY2535 | 51.950723 | 7.599099 | C. Petersen | 2011-12-01 |
| MY882 | RC301 | 54.346355 | 10.117737 | C. Petersen | 2011-12-01 |
| MY887 | MY920 | 54.346355 | 10.117737 | C. Petersen | 2011-12-08 |
| MY904 | MY2453 | 51.950777 | 7.536628 | C. Petersen | 2011-12-08 |
| MY920 | MY920 | 54.346355 | 10.117737 | C. Petersen | 2012-01-18 |
| MY934 | MY2573 | 51.950723 | 7.599099 | C. Petersen | 2012-01-25 |
| MY965 | MY2453 | 51.950777 | 7.536628 | C. Petersen | 2012-05-09 |

| | | | | | |
|---|---|---|---|---|---|
| MY990 | MY920 | 54.346355 | 10.117737 | C. Petersen | 2012-05-23 |
| MY991 | MY679 | 54.346355 | 10.117737 | C. Petersen | 2012-06-20 |
| N2 | N2 | 51.45 | -2.59 | W. Nicholas | 1951 |
| NIC1 | NIC1 | 43.279 | 5.3543 | C. Braendle | 2008-09-14 |
| NIC1049 | NIC1049 | 43.593561 | 1.45032 | C. Braendle | 2014-10-14 |
| NIC1107 | NIC1107 | 43.716278 | 7.266935 | A. Vielle and P. Vigne | 2014-11-25 |
| NIC1119 | NIC1119 | -13.421743 | -71.849508 | C. Braendle | 2015-06-04 |
| NIC1604 | MY2573 | 38.596 | -0.0859 | C. Braendle | 2017-05-01 |
| NIC166 | NIC166 | 46.72722 | 6.89775 | C. Braendle | 2007-07-27 |
| NIC195 | NIC195 | 38.545874 | -28.37322 | D. Bourc'his and J. Dumont | 2011-08-22 |
| NIC196 | NIC195 | 38.4716405 | -28.20149 | D. Bourc'his and J. Dumont | 2011-08-23 |
| NIC197 | NIC195 | 38.39682 | -28.25165 | D. Bourc'his and J. Dumont | 2011-08-23 |
| NIC198 | NIC195 | 39.4364016 | -31.23849 | D. Bourc'his and J. Dumont | 2011-08-29 |
| NIC199 | NIC199 | 37.745428 | -25.19929 | D. Bourc'his and J. Dumont | 2011-03-09 |
| NIC2 | NIC2 | 43.279 | 5.3543 | C. Braendle | 2008-09-14 |
| NIC200 | NIC199 | 37.74942 | -25.66489 | D. Bourc'his and J. Dumont | 2011-05-09 |
| NIC207 | NIC207 | 43.720505 | 7.24045 | C. Braendle | 2011-09-21 |
| NIC231 | NIC231 | 43.720505 | 7.24045 | C. Braendle | 2011-11-14 |
| NIC232 | NIC231 | 43.720505 | 7.24045 | C. Braendle | 2011-11-14 |
| NIC236 | NIC236 | 39.242006 | -9.313417 | C. Braendle | 2011-11-26 |
| NIC237 | NIC236 | 39.242006 | -9.313417 | C. Braendle | 2011-11-26 |
| NIC242 | NIC242 | 38.691765 | -9.31609 | C. Braendle | 2008-08-17 |
| NIC251 | NIC251 | 38.66621 | -28.151364 | S. Carvalho | 2012-08-01 |
| NIC252 | NIC252 | 38.661313 | -28.147949 | S. Carvalho | 2012-08-01 |
| NIC255 | NIC255 | 43.714834 | 7.266779 | C. Braendle | 2012-11-18 |
| NIC256 | NIC256 | 38.71829 | -9.14875 | C. Braendle | 2012-12-02 |
| NIC258 | NIC258 | 38.71829 | -9.14875 | C. Braendle | 2012-12-02 |
| NIC259 | NIC259 | 38.71829 | -9.14875 | C. Braendle | 2012-12-02 |
| NIC260 | NIC260 | 38.71829 | -9.14875 | C. Braendle | 2012-12-02 |
| NIC261 | NIC261 | 38.71829 | -9.14875 | C. Braendle | 2012-12-02 |
| NIC262 | NIC262 | 38.71829 | -9.14875 | C. Braendle | 2012-12-02 |
| NIC263 | NIC256 | 38.71829 | -9.14875 | C. Braendle | 2012-12-02 |
| NIC265 | NIC265 | 38.71829 | -9.14875 | C. Braendle | 2012-12-02 |
| NIC266 | NIC266 | 38.69237 | -9.31592 | C. Braendle | 2012-12-07 |
| NIC267 | NIC267 | 38.69237 | -9.31592 | C. Braendle | 2012-12-07 |
| NIC268 | NIC268 | 38.69237 | -9.31592 | C. Braendle | 2012-12-07 |
| NIC269 | NIC269 | 38.69237 | -9.31592 | C. Braendle | 2012-12-07 |

| | | | | | |
|---|---|---|---|---|---|
| NIC270 | NIC270 | 38.69237 | -9.31592 | C. Braendle | 2012-12-07 |
| NIC271 | NIC271 | 38.69237 | -9.31592 | C. Braendle | 2012-12-07 |
| NIC272 | NIC272 | 39.769459 | -8.756356 | C. Braendle | 2012-12-09 |
| NIC273 | NIC272 | 39.769459 | -8.756356 | C. Braendle | 2012-12-09 |
| NIC274 | NIC274 | 39.769459 | -8.756356 | C. Braendle | 2012-12-09 |
| NIC275 | NIC275 | 39.769459 | -8.756356 | C. Braendle | 2012-12-09 |
| NIC276 | NIC276 | 39.769459 | -8.756356 | C. Braendle | 2012-12-09 |
| NIC277 | NIC277 | 43.716441 | 7.266138 | C. Braendle | 2012-12-15 |
| NIC3 | NIC3 | 43.3636 | 5.3215 | C. Braendle | 2008-09-28 |
| NIC4 | NIC3 | 43.3636 | 5.3215 | C. Braendle | 2008-09-28 |
| NIC501 | NIC501 | 48.77302 | 2.2675 | C. Braendle | 2013-09-14 |
| NIC508 | MY2535 | 40.86183 | 14.261756 | C. Braendle | 2013-11-14 |
| NIC511 | NIC511 | 40.8625 | 14.2623855 | C. Braendle | 2013-11-14 |
| NIC512 | MY2535 | 40.8625 | 14.2623855 | C. Braendle | 2013-11-14 |
| NIC513 | NIC513 | 38.776038 | -9.167136 | C. Braendle | 2013-11-30 |
| NIC514 | NIC514 | 38.776038 | -9.167136 | C. Braendle | 2013-11-30 |
| NIC515 | NIC515 | 38.776775 | -9.165884 | C. Braendle | 2013-11-30 |
| NIC521 | NIC527 | 36.535679 | -6.304441 | C. Braendle | 2014-04-01 |
| NIC522 | NIC522 | 36.535679 | -6.304441 | C. Braendle | 2014-04-01 |
| NIC523 | NIC523 | 36.535679 | -6.29831 | C. Braendle | 2014-04-01 |
| NIC526 | NIC526 | 36.535679 | -6.304441 | C. Braendle | 2014-04-01 |
| NIC527 | NIC527 | 36.535679 | -6.304441 | C. Braendle | 2014-04-01 |
| NIC528 | NIC528 | 36.535679 | -6.29831 | C. Braendle | 2014-04-01 |
| NIC529 | NIC529 | 37.870754 | -4.784967 | C. Braendle | 2014-05-01 |
| PB303 | PB303 | NA | NA | S. Baird | 1998-11-14 |
| PS2025 | PS2025 | 34.19 | -118.13 | J. DeModena | NA |
| PX179 | PX179 | 44.035 | -123.058 | B. White | 2001-10-02 |
| QG2075 | QG2075 | 33.951 | -83.376 | M. Rockman | 2013-02-20 |
| QG2810 | QG2811 | -35.2542 | 149.1151 | M. Rockman | 2017-04-02 |
| QG2811 | QG2811 | -35.2542 | 149.1151 | M. Rockman | 2017-04-02 |
| QG2812 | QG2813 | -34.6342 | 150.7279 | M. Rockman | 2017-03-30 |
| QG2813 | QG2813 | -34.6342 | 150.7279 | M. Rockman | 2017-03-28 |
| QG2818 | QG2818 | -28.0474 | 152.3936 | M. Rockman | 2017-04-12 |
| QG2823 | QG2823 | -34.4511 | 116.033 | M. Rockman | 2017-05-23 |
| QG2824 | QG2824 | -34.4511 | 116.033 | M. Rockman | 2017-05-23 |
| QG2825 | QG2825 | -34.4511 | 116.0328 | M. Rockman | 2017-05-23 |
| QG2826 | QG2823 | -34.4511 | 116.0328 | M. Rockman | 2017-05-23 |

| | | | | | |
|---|---|---|---|---|---|
| QG2827 | QG2827 | -31.9547 | 115.8446 | M. Rockman | 2017-05-24 |
| QG2828 | QG2828 | -31.9547 | 115.8446 | M. Rockman | 2017-05-24 |
| QG2829 | QG2827 | -31.9547 | 115.8446 | M. Rockman | 2017-05-24 |
| QG2830 | QG2832 | -27.3311 | 152.7636 | M. Rockman | 2017-05-31 |
| QG2831 | QG2832 | -27.3311 | 152.7636 | M. Rockman | 2017-05-31 |
| QG2832 | QG2832 | -27.3311 | 152.7636 | M. Rockman | 2017-05-30 |
| QG2833 | QG2832 | -27.3311 | 152.7636 | M. Rockman | 2017-05-30 |
| QG2834 | QG2835 | -27.3308 | 152.7592 | M. Rockman | 2017-05-30 |
| QG2835 | QG2835 | -27.3308 | 152.7592 | M. Rockman | 2017-05-30 |
| QG2836 | QG2836 | -27.3308 | 152.7592 | M. Rockman | 2017-06-01 |
| QG2837 | QG2837 | -27.3308 | 152.7592 | M. Rockman | 2017-06-02 |
| QG2838 | QG2838 | -27.3311 | 152.7636 | M. Rockman | 2017-06-02 |
| QG2839 | QG2841 | -28.2446 | 153.2089 | M. Rockman | 2017-06-06 |
| QG2840 | QG2841 | -28.2446 | 153.2089 | M. Rockman | 2017-06-06 |
| QG2841 | QG2841 | -28.2446 | 153.2063 | M. Rockman | 2017-06-06 |
| QG2842 | QG2841 | -28.2446 | 153.2063 | M. Rockman | 2017-06-06 |
| QG2843 | QG2843 | -28.2402 | 153.195 | M. Rockman | 2017-06-09 |
| QG2844 | QG2846 | -28.2043 | 153.1907 | M. Rockman | 2017-06-06 |
| QG2845 | QG2846 | -28.2043 | 153.1907 | M. Rockman | 2017-06-06 |
| QG2846 | QG2846 | -28.2043 | 153.1907 | M. Rockman | 2017-06-06 |
| QG2850 | QG2850 | -28.0845 | 152.5074 | M. Rockman | 2017-06-22 |
| QG2851 | QG2850 | -28.0845 | 152.5074 | M. Rockman | 2017-06-22 |
| QG2852 | QG2854 | -28.0826 | 152.5066 | M. Rockman | 2017-06-25 |
| QG2853 | QG2854 | -28.0826 | 152.5066 | M. Rockman | 2017-06-25 |
| QG2854 | QG2854 | -28.0806 | 152.5046 | M. Rockman | 2017-06-22 |
| QG2855 | QG2855 | -28.0806 | 152.5046 | M. Rockman | 2017-06-22 |
| QG2856 | QG2854 | -28.0806 | 152.5044 | M. Rockman | 2017-06-20 |
| QG2857 | QG2857 | -28.0806 | 152.5044 | M. Rockman | 2017-06-20 |
| QG2858 | QG2854 | -28.0801 | 152.5041 | M. Rockman | 2017-06-20 |
| QG2859 | QG2854 | -28.0801 | 152.5041 | M. Rockman | 2017-06-20 |
| QG2872 | QG2873 | -27.369 | 152.1831 | M. Rockman | 2017-06-25 |
| QG2873 | QG2873 | -27.369 | 152.1831 | M. Rockman | 2017-06-26 |
| QG2874 | QG2874 | -27.3686 | 152.1838 | M. Rockman | 2017-06-28 |
| QG2875 | QG2875 | -27.3672 | 152.1834 | M. Rockman | 2017-06-27 |
| QG2876 | QG2875 | -27.3672 | 152.1834 | M. Rockman | 2017-06-27 |
| QG2877 | QG2877 | -27.367 | 152.183 | M. Rockman | 2017-06-28 |
| QG2878 | QG2877 | -27.367 | 152.183 | M. Rockman | 2017-06-28 |

| | | | | | |
|---|---|---|---|---|---|
| QG2927 | QG2932 | -28.0538 | 152.3952 | M. Rockman | 2017-07-26 |
| QG2928 | QG2932 | -28.0538 | 152.3952 | M. Rockman | 2017-07-26 |
| QG2931 | QG2932 | -28.0538 | 152.3952 | M. Rockman | 2017-07-26 |
| QG2932 | QG2932 | -28.0538 | 152.3952 | M. Rockman | 2017-07-26 |
| QG536 | QG536 | 37.7679 | -122.4415 | M. Rockman | 2010-12-30 |
| QG537 | QG536 | 37.7679 | -122.4415 | M. Rockman | 2010-12-30 |
| QG538 | QG536 | 37.7679 | -122.4415 | M. Rockman | 2010-12-30 |
| QG556 | QG556 | 34.421629 | -119.70202 | A. Paaby | 2011-06-27 |
| QG557 | QG557 | 34.421629 | -119.70202 | A. Paaby | 2011-06-27 |
| QG558 | QG556 | 34.421629 | -119.70202 | A. Paaby | 2011-06-27 |
| QW947 | QW947 | -33.4213 | -70.6106 | M. Alkema | 2013-04-12 |
| QX1211 | QX1211 | 37.7502 | -122.4331 | M. Rockman | 2007-11-26 |
| QX1212 | QX1212 | 37.7502 | -122.4331 | M. Rockman | 2007-11-26 |
| QX1213 | QX1212 | 37.7502 | -122.4331 | M. Rockman | 2007-11-26 |
| QX1214 | QX1212 | 37.7502 | -122.4331 | M. Rockman | 2007-11-26 |
| QX1215 | QX1211 | 37.7502 | -122.4331 | M. Rockman | 2007-11-26 |
| QX1216 | QX1211 | 37.7502 | -122.4331 | M. Rockman | 2007-11-26 |
| QX1233 | QX1233 | 37.8804 | -122.2838 | M. Rockman | 2007-11-24 |
| QX1791 | QX1791 | 20.6344 | -156.3935 | E. Andersen | 2011-01-27 |
| QX1792 | QX1792 | 20.70554 | -156.35475 | E. Andersen | 2011-01-28 |
| QX1793 | QX1793 | 20.70559 | -156.35678 | E. Andersen | 2011-01-28 |
| QX1794 | QX1794 | 20.70559 | -156.35678 | E. Andersen | 2011-01-28 |
| RC301 | RC301 | 47.99 | 7.84 | R. Cassada | 1983 |
| WN2001 | WN2001 | 51.954422 | 6.303406 | J. Riksen | 2008-06-26 |
| WN2002 | WN2002 | 51.975285 | 5.694834 | J. Riksen | 2007-11-20 |
| WN2010 | WN2001 | 51.954422 | 6.303406 | J. Riksen | 2008-06-26 |
| WN2011 | WN2001 | 51.954422 | 6.303406 | J. Riksen | 2008-06-26 |
| WN2013 | WN2001 | 51.954422 | 6.303406 | J. Riksen | 2008-06-26 |
| WN2014 | WN2001 | 51.954422 | 6.303406 | J. Riksen | 2008-06-26 |
| WN2016 | WN2001 | 51.954422 | 6.303406 | J. Riksen | 2008-06-26 |
| WN2017 | WN2001 | 51.954422 | 6.303406 | J. Riksen | 2008-06-26 |
| WN2018 | WN2001 | 51.954422 | 6.303406 | J. Riksen | 2008-06-26 |
| WN2019 | WN2001 | 51.954422 | 6.303406 | J. Riksen | 2008-06-26 |
| WN2020 | WN2001 | 51.954422 | 6.303406 | J. Riksen | 2008-06-26 |
| WN2021 | WN2001 | 51.954422 | 6.303406 | J. Riksen | 2008-06-26 |
| WN2033 | WN2033 | 51.975 | 5.694 | J. Riksen | 2014-11-05 |
| WN2035 | WN2033 | 51.975 | 5.694 | J. Riksen | 2014-11-05 |

| | | | | | |
|---|---|---|---|---|---|
| WN2039 | WN2033 | 51.975 | 5.694 | J. Riksen | 2014-11-05 |
| WN2050 | WN2050 | 51.975 | 5.717 | J. Riksen | 2014-11-11 |
| WN2056 | WN2050 | 51.975 | 5.717 | J. Riksen | 2014-11-11 |
| WN2059 | WN2066 | 51.975306 | 5.694904 | M. Sterken | 2016-10-11 |
| WN2060 | WN2066 | 51.975306 | 5.694904 | M. Sterken | 2016-10-11 |
| WN2062 | WN2050 | 51.975196 | 5.718727 | M. Sterken | 2016-10-11 |
| WN2063 | WN2063 | 51.974975 | 5.651316 | M. Gultom | 2016-10-11 |
| WN2064 | WN2064 | 51.974703 | 5.68639 | M. van Wijk | 2016-10-11 |
| WN2065 | WN2066 | 51.975306 | 5.694904 | Y. Huang | 2016-10-11 |
| WN2066 | WN2066 | 51.975306 | 5.694904 | Y. Huang | 2016-10-11 |
| XZ1513 | XZ1513 | 22.14787 | -159.63105 | M. Ailion | 2014-10-08 |
| XZ1514 | XZ1514 | 22.149 | -159.668 | M. Ailion | 2014-10-15 |
| XZ1515 | XZ1515 | 22.149 | -159.668 | M. Ailion | 2014-10-15 |
| XZ1516 | XZ1516 | 22.149 | -159.668 | M. Ailion | 2014-10-15 |
| XZ1672 | XZ1672 | 47.687741 | -122.30051 | B. Coleman | 2015-08-01 |
| XZ1734 | XZ1734 | -23.5 | -46.7 | M. Ailion | 2015-11-15 |
| XZ1735 | XZ1735 | -22.9 | -46.7 | M. Ailion | 2015-11-07 |
| XZ1756 | XZ1756 | 47.623232 | -122.30732 | P. Lamelza | 2015-10-24 |
| XZ2018 | XZ2018 | NA | | M. Ailion | 2016-10-01 |
| XZ2019 | XZ2019 | 47.636483 | -122.30589 | M. Ailion | 2016-10-25 |
| XZ2020 | XZ2020 | 46.3 | -120 | A. Nassar | 2016-10-01 |

**Supplementary Table 2 | Distances among wild strains that belong to the same isotype**
The geographic information of 14 isotypes that were sampled multiple times from locations at least 50 km apart is shown.

| Strain | Isotype | Latitude | Longitude | Isotype strain counts | Is reference | Distance to reference strain (km) |
|---|---|---|---|---|---|---|
| AB4 | ECA251 | -34.93 | 138.59 | 2 | FALSE | 13170.5973987643 |
| ECA251 | ECA251 | 34.1 | -118.1 | 2 | TRUE | NA |
| JU1200 | JU1200 | 55.577 | -4.6 | 2 | TRUE | NA |
| JU2860 | JU1200 | 52.19842 | 0.115365 | 2 | FALSE | 487.152613837801 |
| JU1440 | JU1440 | 41.41307 | 2.15231 | 2 | TRUE | NA |
| JU3133 | JU1440 | 48.7015 | 2.1725 | 2 | FALSE | 809.98596260867 |
| JU1793 | JU1793 | 49.12604 | 1.95114 | 2 | TRUE | NA |
| JU2139 | JU1793 | 46.63278 | 1.064289 | 2 | FALSE | 285.040917449512 |
| JU311 | JU311 | 44.42 | 4.4 | 2 | TRUE | NA |
| MY2481 | JU311 | 54.346355 | 10.117737 | 2 | FALSE | 1178.49124674857 |
| JU3227 | JU3228 | -38.64844 | 176.08947 | 2 | FALSE | 52.767605399004 |
| JU3228 | JU3228 | -39.02256 | 175.71458 | 2 | TRUE | NA |
| JU3131 | JU323 | 48.7015 | 2.1725 | 2 | FALSE | 505.599584406246 |
| JU323 | JU323 | 44.42 | 4.4 | 2 | TRUE | NA |
| JU2527 | JU792 | 47.8797 | -3.5999 | 3 | FALSE | 613.897518387652 |
| JU3142 | JU792 | 48.71 | -3.81 | 3 | FALSE | 702.031724447556 |
| JU792 | JU792 | 43.06 | 0.24 | 3 | TRUE | NA |
| BRC20231 | MY23 | 23.541517 | 120.908112 | 2 | FALSE | 9447.77982000758 |
| MY23 | MY23 | 51.96 | 7.53 | 2 | TRUE | NA |
| JU3136 | MY2453 | 48.7015 | 2.1725 | 3 | FALSE | 525.612240370477 |
| MY2453 | MY2453 | 51.950777 | 7.536628 | 3 | TRUE | NA |
| MY2622 | MY2453 | 54.3491 | 10.11505 | 3 | FALSE | 317.753858523063 |
| MY2532 | MY2535 | 54.346355 | 10.117737 | 3 | FALSE | 315.352444531987 |
| MY2535 | MY2535 | 51.950723 | 7.599099 | 3 | TRUE | NA |
| NIC508 | MY2535 | 40.86183 | 14.261756 | 3 | FALSE | 1333.4739542811 |
| ECA552 | MY2573 | 43.8658 | 5.0628 | 4 | FALSE | 918.573851024173 |
| MY2208 | MY2573 | 54.346355 | 10.117737 | 4 | FALSE | 315.352444531987 |
| MY2573 | MY2573 | 51.950723 | 7.599099 | 4 | TRUE | NA |
| NIC1604 | MY2573 | 38.596 | -0.0859 | 4 | FALSE | 1599.85744272238 |
| NIC195 | NIC195 | 38.545874 | -28.37322 | 2 | TRUE | NA |
| NIC198 | NIC195 | 39.4364016 | -31.23849 | 2 | FALSE | 267.18451568098 |
| MY2024 | RC301 | 54.346355 | 10.117737 | 2 | FALSE | 724.773320793207 |
| RC301 | RC301 | 47.99 | 7.84 | 2 | TRUE | NA |

**Supplementary Table 3 | Distribution of swept haplotype and hyper-divergent genome across wild *C. elegans* isotypes**

The variant distribution for the 328 *C. elegans* isotypes is shown. The geographic origin of the isotype is shown in the second column. The fraction swept columns refer to the fraction of chromosomes I, IV, V, and X that each strain shares with the most abundant haplotype (Methods). The hyper-divergent genome column refers to the amount of genome (kb) that was classified as hyper-divergent and the fraction hyper-divergent fraction column is the fraction of the genome that is classified as hyper-divergent. The genome-wide variants column represents the total number of ALT variant calls for each isotype, and the divergent-region variants column shows the number of variants localized to hyper-divergent regions. Finally, the fraction divergent-region variants column shows the fraction of total variants that are localized to the hyper-divergent regions.

| Isotype reference strain | Origin | Fraction swept (I, IV, V, X) | | | | Hyper-divergent genome (kb) | Fraction hyper-divergent | Genome-wide variants | Divergent-region variants | Fraction divergent-region variants |
|---|---|---|---|---|---|---|---|---|---|---|
| AB1 | Australia | 0.695 | 0.601 | 0.805 | 0.761 | 1162 | 0.012 | 92517 | 23363 | 0.253 |
| BRC20067 | Asia | 0.300 | 1.000 | 0.904 | 1.000 | 826 | 0.008 | 89685 | 10905 | 0.122 |
| BRC20263 | Australia | 0.323 | 0.735 | 0.367 | 0.690 | 2000 | 0.020 | 153922 | 39114 | 0.254 |
| CB4852 | Unknown | 0.817 | 0.904 | 0.904 | 0.588 | 528 | 0.005 | 67966 | 10002 | 0.147 |
| CB4854 | N. America | 0.407 | 0.853 | 0.903 | 0.653 | 980 | 0.010 | 98343 | 14154 | 0.144 |
| CB4856 | Hawaii | 0.000 | 0.000 | 0.000 | 0.000 | 3224 | 0.032 | 242577 | 58514 | 0.241 |
| CB4932 | Europe | 0.262 | 0.000 | 0.768 | 0.058 | 710 | 0.007 | 88756 | 13888 | 0.156 |
| CX11254 | N. America | 0.000 | 0.870 | 0.313 | 0.272 | 2001 | 0.020 | 136736 | 35033 | 0.256 |
| CX11262 | N. America | 0.000 | 0.655 | 0.263 | 0.170 | 2137 | 0.021 | 165389 | 41765 | 0.253 |
| CX11264 | N. America | 0.237 | 0.561 | 0.234 | 0.000 | 2112 | 0.021 | 177791 | 43398 | 0.244 |
| CX11271 | N. America | 0.237 | 0.565 | 0.792 | 0.306 | 667 | 0.007 | 109851 | 12986 | 0.118 |
| CX11276 | N. America | 0.646 | 0.742 | 0.717 | 0.327 | 1804 | 0.018 | 117544 | 26202 | 0.223 |
| CX11285 | N. America | 0.096 | 0.555 | 0.129 | 0.361 | 2173 | 0.022 | 169882 | 35749 | 0.210 |
| CX11292 | N. America | 0.318 | 0.743 | 0.904 | 0.306 | 682 | 0.007 | 97849 | 12768 | 0.130 |
| CX11307 | N. America | 0.222 | 0.973 | 0.241 | 0.217 | 2026 | 0.020 | 149073 | 43265 | 0.290 |
| CX11314 | N. America | 0.311 | 0.740 | 0.282 | 0.306 | 2139 | 0.021 | 147343 | 38304 | 0.260 |
| CX11315 | N. America | 0.096 | 0.561 | 0.663 | 0.553 | 2121 | 0.021 | 134796 | 37884 | 0.281 |
| DL200 | Africa | 0.693 | 0.759 | 0.682 | 0.000 | 1258 | 0.013 | 123175 | 17745 | 0.144 |
| DL226 | N. America | 0.187 | 0.768 | 0.805 | 0.618 | 2039 | 0.020 | 143436 | 32910 | 0.229 |
| DL238 | Hawaii | 0.000 | 0.000 | 0.000 | 0.000 | 3206 | 0.032 | 260360 | 69449 | 0.267 |
| ECA189 | Hawaii | 0.000 | 0.000 | 0.000 | 0.000 | 2987 | 0.030 | 231297 | 58893 | 0.255 |
| ECA191 | Hawaii | 0.000 | 0.000 | 0.000 | 0.000 | 3856 | 0.038 | 426268 | 80204 | 0.188 |
| ECA243 | Europe | 0.292 | 0.000 | 0.640 | 0.134 | 510 | 0.005 | 61864 | 13801 | 0.223 |
| ECA246 | N. America | 0.631 | 0.765 | 0.904 | 0.500 | 1387 | 0.014 | 108790 | 19480 | 0.179 |
| ECA248 | N. America | 0.237 | 0.381 | 0.655 | 0.222 | 1379 | 0.014 | 130786 | 21273 | 0.163 |
| ECA250 | N. America | 0.177 | 0.743 | 0.902 | 0.139 | 1516 | 0.015 | 102241 | 17874 | 0.175 |
| ECA251 | N. America | 0.318 | 0.743 | 0.904 | 0.306 | 1339 | 0.013 | 112507 | 19243 | 0.171 |

| | | | | | | | | | | |
|---|---|---|---|---|---|---|---|---|---|---|
| ECA259 | Unknown | 0.096 | 0.743 | 0.106 | 0.092 | 1739 | 0.017 | 159112 | 33239 | 0.209 |
| ECA347 | Hawaii | 0.000 | 0.000 | 0.000 | 0.000 | 4592 | 0.046 | 463961 | 99679 | 0.215 |
| ECA348 | N. America | 0.081 | 0.432 | 0.127 | 0.000 | 2417 | 0.024 | 202138 | 49636 | 0.246 |
| ECA349 | N. America | 0.318 | 0.593 | 0.682 | 0.136 | 1589 | 0.016 | 138595 | 27144 | 0.196 |
| ECA36 | New Zealand | 0.000 | 0.000 | 0.000 | 0.000 | 5262 | 0.052 | 483212 | 123370 | 0.255 |
| ECA363 | Hawaii | 0.000 | 0.000 | 0.000 | 0.000 | 4813 | 0.048 | 459879 | 105428 | 0.229 |
| ECA369 | Hawaii | 0.000 | 0.000 | 0.000 | 0.000 | 7670 | 0.076 | 386353 | 194678 | 0.504 |
| ECA372 | Hawaii | 0.000 | 0.000 | 0.000 | 0.000 | 2925 | 0.029 | 219397 | 53775 | 0.245 |
| ECA396 | Hawaii | 0.000 | 0.000 | 0.000 | 0.000 | 4138 | 0.041 | 445019 | 89316 | 0.201 |
| ECA592 | N. America | 0.000 | 0.000 | 0.286 | 0.171 | 2007 | 0.020 | 194179 | 36248 | 0.187 |
| ECA593 | N. America | 0.000 | 0.163 | 0.000 | 0.000 | 2723 | 0.027 | 238820 | 51048 | 0.214 |
| ECA594 | N. America | 0.154 | 0.102 | 0.097 | 0.329 | 2504 | 0.025 | 207667 | 48082 | 0.232 |
| ECA640 | N. America | 0.279 | 0.443 | 0.079 | 0.524 | 1857 | 0.019 | 168016 | 43341 | 0.258 |
| ECA701 | Hawaii | 0.000 | 0.000 | 0.000 | 0.000 | 9903 | 0.099 | 580298 | 258199 | 0.445 |
| ECA703 | Hawaii | 0.000 | 0.000 | 0.000 | 0.000 | 2809 | 0.028 | 257175 | 53718 | 0.209 |
| ECA705 | Hawaii | 0.000 | 0.000 | 0.000 | 0.000 | 3070 | 0.031 | 235110 | 55849 | 0.238 |
| ECA706 | Hawaii | 0.000 | 0.000 | 0.000 | 0.000 | 2441 | 0.024 | 243854 | 51853 | 0.213 |
| ECA710 | Hawaii | 0.000 | 0.000 | 0.000 | 0.000 | 2485 | 0.025 | 226762 | 50729 | 0.224 |
| ECA712 | Hawaii | 0.000 | 0.000 | 0.000 | 0.000 | 2884 | 0.029 | 225399 | 50786 | 0.225 |
| ECA722 | Hawaii | 0.000 | 0.000 | 0.000 | 0.000 | 3615 | 0.036 | 444068 | 79778 | 0.180 |
| ECA723 | Hawaii | 0.000 | 0.000 | 0.000 | 0.000 | 3366 | 0.034 | 431904 | 72976 | 0.169 |
| ECA724 | Hawaii | 0.000 | 0.000 | 0.000 | 0.000 | 4326 | 0.043 | 437044 | 91298 | 0.209 |
| ECA730 | Hawaii | 0.000 | 0.000 | 0.000 | 0.000 | 2966 | 0.030 | 420411 | 63875 | 0.152 |
| ECA732 | Hawaii | 0.000 | 0.000 | 0.000 | 0.000 | 3017 | 0.030 | 256960 | 66813 | 0.260 |
| ECA733 | Hawaii | 0.000 | 0.000 | 0.000 | 0.000 | 3001 | 0.030 | 250557 | 62046 | 0.248 |
| ECA738 | Hawaii | 0.000 | 0.000 | 0.000 | 0.000 | 2422 | 0.024 | 227308 | 48817 | 0.215 |
| ECA740 | Hawaii | 0.000 | 0.000 | 0.000 | 0.000 | 4939 | 0.049 | 486776 | 119954 | 0.246 |
| ECA741 | Hawaii | 0.000 | 0.000 | 0.000 | 0.000 | 4907 | 0.049 | 480385 | 110565 | 0.230 |
| ECA742 | Hawaii | 0.000 | 0.000 | 0.000 | 0.000 | 5538 | 0.055 | 498928 | 133335 | 0.267 |
| ECA743 | Hawaii | 0.000 | 0.000 | 0.121 | 0.000 | 2373 | 0.024 | 225101 | 48674 | 0.216 |
| ECA744 | Hawaii | 0.000 | 0.000 | 0.000 | 0.000 | 3377 | 0.034 | 426617 | 73389 | 0.172 |
| ECA745 | Hawaii | 0.000 | 0.000 | 0.000 | 0.000 | 3776 | 0.038 | 440983 | 83326 | 0.189 |
| ECA746 | Hawaii | 0.000 | 0.000 | 0.000 | 0.000 | 3446 | 0.034 | 434102 | 77631 | 0.179 |
| ECA760 | Hawaii | 0.000 | 0.000 | 0.000 | 0.000 | 2124 | 0.021 | 226120 | 44842 | 0.198 |
| ECA768 | Hawaii | 0.000 | 0.000 | 0.000 | 0.000 | 2391 | 0.024 | 236885 | 51418 | 0.217 |
| ECA777 | Hawaii | 0.000 | 0.000 | 0.000 | 0.000 | 2030 | 0.020 | 225133 | 43788 | 0.194 |
| ECA778 | Hawaii | 0.000 | 0.000 | 0.000 | 0.000 | 2287 | 0.023 | 223197 | 48527 | 0.217 |

| | | | | | | | | | | |
|---|---|---|---|---|---|---|---|---|---|---|
| ECA807 | Hawaii | 0.000 | 0.000 | 0.000 | 0.000 | 2737 | 0.027 | 241101 | 47559 | 0.197 |
| ECA812 | Hawaii | 0.000 | 0.000 | 0.000 | 0.000 | 2753 | 0.027 | 226067 | 48104 | 0.213 |
| ECA923 | Hawaii | 0.073 | 0.657 | 0.536 | 0.170 | 2563 | 0.026 | 166996 | 42958 | 0.257 |
| ECA928 | Hawaii | 0.073 | 0.434 | 0.204 | 0.000 | 2360 | 0.024 | 188816 | 43662 | 0.231 |
| ECA930 | Asia | 0.237 | 0.562 | 0.770 | 0.170 | 1970 | 0.020 | 132581 | 30552 | 0.230 |
| ED3005 | Europe | 0.351 | 0.562 | 0.770 | 0.170 | 2060 | 0.021 | 137514 | 32800 | 0.239 |
| ED3011 | Europe | 0.318 | 0.443 | 0.901 | 0.611 | 2088 | 0.021 | 113708 | 24193 | 0.213 |
| ED3012 | Europe | 0.292 | 0.803 | 0.904 | 0.616 | 841 | 0.008 | 47914 | 13894 | 0.290 |
| ED3017 | Europe | 0.749 | 0.825 | 0.710 | 0.485 | 1147 | 0.011 | 115601 | 20291 | 0.176 |
| ED3040 | Africa | 0.073 | 0.645 | 0.674 | 0.756 | 998 | 0.010 | 121638 | 14838 | 0.122 |
| ED3046 | Africa | 0.206 | 0.153 | 0.465 | 0.256 | 2045 | 0.020 | 152355 | 34163 | 0.224 |
| ED3048 | Africa | 0.874 | 0.882 | 0.901 | 0.992 | 875 | 0.009 | 91113 | 13063 | 0.143 |
| ED3049 | Africa | 0.206 | 0.153 | 0.465 | 0.256 | 2049 | 0.020 | 152132 | 34228 | 0.225 |
| ED3052 | Africa | 0.900 | 0.228 | 0.763 | 0.500 | 1035 | 0.010 | 119790 | 19286 | 0.161 |
| ED3073 | Africa | 0.384 | 1.000 | 0.811 | 0.871 | 1029 | 0.010 | 115599 | 15888 | 0.137 |
| ED3077 | Africa | 0.900 | 1.000 | 0.588 | 0.992 | 1500 | 0.015 | 130076 | 25604 | 0.197 |
| EG4347 | N. America | 0.197 | 0.765 | 0.904 | 0.471 | 650 | 0.006 | 48038 | 8338 | 0.174 |
| EG4349 | N. America | 0.410 | 0.072 | 0.394 | 0.611 | 1946 | 0.019 | 172747 | 37824 | 0.219 |
| EG4724 | Europe | 0.703 | 0.292 | 0.342 | 0.508 | 1900 | 0.019 | 163976 | 36931 | 0.225 |
| EG4725 | Europe | 0.277 | 0.164 | 0.218 | 0.421 | 3000 | 0.030 | 206758 | 61174 | 0.296 |
| EG4946 | N. America | 0.368 | 0.797 | 0.904 | 0.530 | 1178 | 0.012 | 89578 | 18468 | 0.206 |
| GXW1 | Asia | 0.287 | 0.532 | 0.953 | 0.727 | 960 | 0.010 | 118217 | 19965 | 0.169 |
| JT11398 | N. America | 0.193 | 0.738 | 0.868 | 0.489 | 566 | 0.006 | 85909 | 9837 | 0.115 |
| JU1088 | Asia | 0.302 | 1.000 | 0.904 | 0.959 | 1961 | 0.020 | 105264 | 28900 | 0.275 |
| JU1172 | S. America | 0.318 | 0.743 | 0.872 | 0.306 | 1342 | 0.013 | 112780 | 20476 | 0.182 |
| JU1200 | Europe | 0.292 | 0.863 | 0.904 | 0.710 | 61 | 0.001 | 21836 | 1155 | 0.053 |
| JU1212 | Europe | 0.098 | 0.000 | 0.438 | 0.134 | 1735 | 0.017 | 102086 | 31593 | 0.309 |
| JU1213 | Europe | 0.621 | 0.000 | 0.431 | 0.192 | 1896 | 0.019 | 134478 | 30788 | 0.229 |
| JU1242 | Europe | 0.532 | 0.604 | 0.740 | 0.000 | 892 | 0.009 | 117997 | 15018 | 0.127 |
| JU1246 | Europe | 0.194 | 0.000 | 0.644 | 0.117 | 1157 | 0.012 | 121086 | 19486 | 0.161 |
| JU1249 | Europe | 0.382 | 0.897 | 0.904 | 0.134 | 429 | 0.004 | 33832 | 8988 | 0.266 |
| JU1395 | Europe | 0.823 | 0.811 | 0.904 | 0.588 | 615 | 0.006 | 66203 | 9305 | 0.141 |
| JU1400 | Europe | 0.229 | 0.825 | 0.538 | 0.186 | 2282 | 0.023 | 175806 | 49100 | 0.279 |
| JU1409 | Europe | 0.413 | 0.461 | 0.868 | 0.545 | 810 | 0.008 | 105797 | 13291 | 0.126 |
| JU1440 | Europe | 0.629 | 0.538 | 0.800 | 0.667 | 1113 | 0.011 | 98731 | 21867 | 0.221 |
| JU1491 | Europe | 0.590 | 0.000 | 0.701 | 0.400 | 1285 | 0.013 | 137678 | 26518 | 0.193 |
| JU1530 | Europe | 0.402 | 0.885 | 0.774 | 0.202 | 780 | 0.008 | 108829 | 13397 | 0.123 |

| | | | | | | | | | | |
|---|---|---|---|---|---|---|---|---|---|---|
| JU1543 | Europe | 0.808 | 0.729 | 0.822 | 0.749 | 269 | 0.003 | 38858 | 6847 | 0.176 |
| JU1568 | Europe | 0.292 | 0.526 | 0.820 | 0.297 | 128 | 0.001 | 34873 | 1569 | 0.045 |
| JU1580 | Europe | 0.531 | 0.000 | 0.000 | 0.000 | 2548 | 0.025 | 209856 | 56063 | 0.267 |
| JU1581 | Europe | 0.000 | 0.000 | 0.768 | 0.000 | 1052 | 0.010 | 148575 | 21400 | 0.144 |
| JU1586 | Europe | 0.292 | 0.653 | 0.816 | 0.562 | 384 | 0.004 | 42094 | 7876 | 0.187 |
| JU1652 | S. America | 0.692 | 0.121 | 0.718 | 0.936 | 1848 | 0.018 | 121191 | 28738 | 0.237 |
| JU1666 | Europe | 0.627 | 0.000 | 0.652 | 0.000 | 1471 | 0.015 | 145236 | 23121 | 0.159 |
| JU1792 | Europe | 0.292 | 0.536 | 0.756 | 0.058 | 812 | 0.008 | 83277 | 16110 | 0.193 |
| JU1793 | Europe | 0.535 | 0.000 | 0.000 | 0.000 | 2749 | 0.027 | 206545 | 57040 | 0.276 |
| JU1808 | Europe | 0.278 | 0.443 | 0.808 | 0.149 | 984 | 0.010 | 111766 | 16386 | 0.147 |
| JU1896 | Europe | 0.000 | 0.000 | 0.868 | 0.791 | 1458 | 0.015 | 117869 | 25535 | 0.217 |
| JU1934 | Europe | 0.506 | 0.000 | 0.740 | 0.058 | 1010 | 0.010 | 121568 | 18708 | 0.154 |
| JU2001 | Africa | 0.516 | 0.000 | 0.473 | 0.000 | 2121 | 0.021 | 187815 | 43047 | 0.229 |
| JU2007 | Europe | 0.264 | 0.656 | 0.622 | 0.638 | 955 | 0.010 | 123027 | 19470 | 0.158 |
| JU2016 | N. America | 0.318 | 0.743 | 0.904 | 0.306 | 1220 | 0.012 | 105254 | 17098 | 0.162 |
| JU2017 | N. America | 0.215 | 0.306 | 0.682 | 0.516 | 1597 | 0.016 | 151825 | 27993 | 0.184 |
| JU2106 | Europe | 0.506 | 0.000 | 0.000 | 0.000 | 1973 | 0.020 | 165575 | 41744 | 0.252 |
| JU2131 | Europe | 0.627 | 0.000 | 0.600 | 0.000 | 1470 | 0.015 | 149803 | 24404 | 0.163 |
| JU2141 | Europe | 0.506 | 0.000 | 0.740 | 0.058 | 894 | 0.009 | 120359 | 15366 | 0.128 |
| JU2234 | Europe | 0.435 | 0.063 | 0.768 | 0.807 | 991 | 0.010 | 116614 | 20118 | 0.173 |
| JU2250 | Europe | 0.497 | 0.000 | 0.692 | 0.000 | 824 | 0.008 | 136220 | 15054 | 0.111 |
| JU2257 | Europe | 0.292 | 0.093 | 0.771 | 0.134 | 262 | 0.003 | 47142 | 6514 | 0.138 |
| JU2316 | Europe | 0.263 | 0.000 | 0.058 | 0.278 | 3271 | 0.033 | 257407 | 68021 | 0.264 |
| JU2464 | S. America | 0.775 | 0.563 | 0.904 | 0.488 | 1623 | 0.016 | 122126 | 22039 | 0.180 |
| JU2466 | S. America | 0.814 | 0.994 | 0.900 | 1.000 | 673 | 0.007 | 94560 | 10939 | 0.116 |
| JU2478 | Europe | 0.341 | 0.735 | 0.904 | 0.214 | 717 | 0.007 | 106652 | 11520 | 0.108 |
| JU2513 | Europe | 0.784 | 0.911 | 0.825 | 0.691 | 1605 | 0.016 | 116827 | 27635 | 0.237 |
| JU2519 | Europe | 0.433 | 0.199 | 0.058 | 0.000 | 2628 | 0.026 | 207698 | 49843 | 0.240 |
| JU2522 | Europe | 0.312 | 0.234 | 0.354 | 0.424 | 1639 | 0.016 | 154808 | 30249 | 0.195 |
| JU2526 | Europe | 0.219 | 0.069 | 0.058 | 0.000 | 3793 | 0.038 | 269650 | 72579 | 0.269 |
| JU2534 | Europe | 0.657 | 0.944 | 0.867 | 0.075 | 836 | 0.008 | 89081 | 12105 | 0.136 |
| JU2565 | Europe | 0.679 | 0.000 | 0.820 | 0.134 | 130 | 0.001 | 41682 | 1578 | 0.038 |
| JU2566 | Europe | 0.292 | 0.488 | 0.708 | 0.058 | 359 | 0.004 | 57003 | 8235 | 0.144 |
| JU2570 | Europe | 0.614 | 0.000 | 0.740 | 0.000 | 913 | 0.009 | 135159 | 16090 | 0.119 |
| JU2572 | Europe | 0.303 | 0.825 | 0.792 | 0.058 | 1036 | 0.010 | 79883 | 22719 | 0.284 |
| JU2575 | Europe | 0.292 | 0.824 | 0.723 | 0.076 | 489 | 0.005 | 59024 | 7854 | 0.133 |
| JU2576 | Europe | 0.000 | 0.000 | 0.762 | 0.166 | 1350 | 0.013 | 142266 | 22027 | 0.155 |

| | | | | | | | | | | |
|---|---|---|---|---|---|---|---|---|---|---|
| JU2578 | Europe | 0.176 | 0.760 | 0.868 | 0.812 | 974 | 0.010 | 118552 | 17081 | 0.144 |
| JU258 | Atlantic | 0.000 | 0.000 | 0.274 | 0.000 | 3509 | 0.035 | 206928 | 54206 | 0.262 |
| JU2581 | Europe | 0.141 | 0.184 | 0.482 | 0.464 | 1561 | 0.016 | 144025 | 24825 | 0.172 |
| JU2586 | Europe | 0.000 | 0.462 | 0.342 | 0.545 | 1898 | 0.019 | 169120 | 33087 | 0.196 |
| JU2587 | Europe | 0.610 | 0.239 | 0.666 | 0.605 | 2162 | 0.022 | 153502 | 39383 | 0.257 |
| JU2592 | Europe | 0.292 | 0.397 | 0.564 | 0.058 | 1455 | 0.015 | 98664 | 23338 | 0.237 |
| JU2593 | Europe | 0.305 | 0.068 | 0.758 | 0.354 | 1859 | 0.019 | 154999 | 30011 | 0.194 |
| JU2600 | Europe | 0.611 | 0.000 | 0.703 | 0.000 | 1479 | 0.015 | 139840 | 29012 | 0.207 |
| JU2610 | Europe | 0.567 | 0.000 | 0.703 | 0.000 | 1716 | 0.017 | 141986 | 30897 | 0.218 |
| JU2619 | N. America | 0.073 | 1.000 | 0.710 | 0.592 | 1904 | 0.019 | 139960 | 31258 | 0.223 |
| JU2800 | Europe | 0.452 | 0.000 | 0.902 | 0.149 | 1323 | 0.013 | 133149 | 21096 | 0.158 |
| JU2811 | Australia | 0.319 | 0.000 | 0.379 | 0.705 | 2177 | 0.022 | 182212 | 43729 | 0.240 |
| JU2825 | Europe | 0.292 | 0.000 | 0.613 | 0.134 | 1037 | 0.010 | 86020 | 25271 | 0.294 |
| JU2829 | Europe | 0.251 | 0.624 | 0.482 | 0.495 | 1495 | 0.015 | 153621 | 26792 | 0.174 |
| JU2838 | N. America | 0.376 | 0.627 | 0.219 | 0.131 | 1709 | 0.017 | 159833 | 33717 | 0.211 |
| JU2841 | New Zealand | 0.129 | 0.504 | 0.730 | 0.546 | 1639 | 0.016 | 146135 | 30703 | 0.210 |
| JU2853 | Europe | 0.303 | 0.874 | 0.820 | 0.692 | 338 | 0.003 | 68256 | 4556 | 0.067 |
| JU2862 | Europe | 0.373 | 0.639 | 0.868 | 0.145 | 870 | 0.009 | 112976 | 12541 | 0.111 |
| JU2866 | N. America | 0.161 | 0.706 | 0.772 | 0.160 | 1556 | 0.016 | 135682 | 29032 | 0.214 |
| JU2878 | N. America | 0.426 | 0.753 | 0.660 | 0.819 | 1743 | 0.017 | 150327 | 34140 | 0.227 |
| JU2879 | N. America | 0.000 | 0.557 | 0.332 | 0.167 | 2313 | 0.023 | 188605 | 49937 | 0.265 |
| JU2906 | Europe | 0.403 | 0.858 | 0.853 | 0.464 | 694 | 0.007 | 78203 | 10326 | 0.132 |
| JU2907 | Europe | 0.151 | 0.939 | 0.853 | 0.669 | 444 | 0.004 | 40348 | 5445 | 0.135 |
| JU310 | Europe | 0.495 | 0.885 | 0.774 | 0.202 | 827 | 0.008 | 110575 | 14902 | 0.135 |
| JU311 | Europe | 0.292 | 0.000 | 0.866 | 0.149 | 1176 | 0.012 | 84376 | 14319 | 0.170 |
| JU3125 | Europe | 0.605 | 0.737 | 0.458 | 0.632 | 1335 | 0.013 | 117986 | 25564 | 0.217 |
| JU3127 | Europe | 0.603 | 0.635 | 0.458 | 0.436 | 1387 | 0.014 | 124637 | 25540 | 0.205 |
| JU3128 | Europe | 0.376 | 0.778 | 0.853 | 0.276 | 702 | 0.007 | 93411 | 10329 | 0.111 |
| JU3132 | Europe | 0.402 | 0.000 | 0.712 | 0.066 | 999 | 0.010 | 134611 | 15550 | 0.116 |
| JU3134 | Europe | 0.151 | 0.309 | 0.868 | 0.902 | 1727 | 0.017 | 119861 | 27718 | 0.231 |
| JU3135 | Europe | 0.092 | 0.950 | 0.904 | 0.873 | 1240 | 0.012 | 104695 | 21465 | 0.205 |
| JU3137 | Europe | 0.482 | 0.000 | 0.768 | 0.701 | 1239 | 0.012 | 135633 | 22552 | 0.166 |
| JU3140 | Europe | 0.614 | 0.000 | 0.685 | 0.000 | 1130 | 0.011 | 144483 | 17782 | 0.123 |
| JU3141 | Europe | 0.292 | 0.000 | 0.756 | 0.058 | 772 | 0.008 | 96197 | 15055 | 0.157 |
| JU3144 | Africa | 0.410 | 0.727 | 0.512 | 0.587 | 1822 | 0.018 | 148206 | 35131 | 0.237 |
| JU3166 | Atlantic | 0.155 | 0.230 | 0.354 | 0.000 | 2466 | 0.025 | 196219 | 44301 | 0.226 |
| JU3167 | Atlantic | 0.073 | 0.000 | 0.000 | 0.140 | 2796 | 0.028 | 233185 | 60051 | 0.258 |

| | | | | | | | | | | |
|---|---|---|---|---|---|---|---|---|---|---|
| JU3169 | Atlantic | 0.073 | 0.000 | 0.403 | 0.140 | 1874 | 0.019 | 194808 | 31708 | 0.163 |
| JU3224 | New Zealand | 0.413 | 0.678 | 0.383 | 0.730 | 1999 | 0.020 | 150166 | 34389 | 0.229 |
| JU3225 | New Zealand | 0.181 | 0.679 | 0.904 | 0.678 | 1008 | 0.010 | 99429 | 17591 | 0.177 |
| JU3226 | New Zealand | 0.000 | 0.000 | 0.000 | 0.000 | 5767 | 0.058 | 451149 | 120800 | 0.268 |
| JU3228 | New Zealand | 0.328 | 0.252 | 0.590 | 0.756 | 2237 | 0.022 | 158485 | 36471 | 0.230 |
| JU323 | Europe | 0.992 | 0.534 | 0.917 | 0.000 | 1178 | 0.012 | 113780 | 18307 | 0.161 |
| JU3280 | Europe | 0.679 | 0.099 | 0.556 | 0.762 | 1436 | 0.014 | 137853 | 24509 | 0.178 |
| JU3282 | Europe | 0.977 | 1.000 | 0.949 | 0.636 | 1539 | 0.015 | 116377 | 28173 | 0.242 |
| JU3291 | Europe | 0.611 | 0.000 | 0.703 | 0.000 | 1424 | 0.014 | 139762 | 28136 | 0.201 |
| JU346 | Europe | 0.240 | 0.336 | 0.723 | 0.199 | 982 | 0.010 | 119808 | 16370 | 0.137 |
| JU360 | Europe | 0.305 | 0.829 | 0.717 | 0.476 | 1585 | 0.016 | 135813 | 29711 | 0.219 |
| JU367 | Europe | 0.292 | 0.706 | 0.948 | 0.134 | 355 | 0.004 | 47201 | 4533 | 0.096 |
| JU393 | Europe | 0.587 | 0.000 | 0.654 | 0.262 | 2038 | 0.020 | 135471 | 39470 | 0.291 |
| JU394 | Europe | 0.292 | 0.000 | 0.640 | 0.134 | 559 | 0.006 | 59583 | 13390 | 0.225 |
| JU397 | Europe | 0.506 | 0.000 | 0.740 | 0.000 | 1075 | 0.011 | 111079 | 16870 | 0.152 |
| JU406 | Europe | 0.292 | 0.000 | 0.679 | 0.134 | 546 | 0.005 | 63009 | 11367 | 0.180 |
| JU440 | Europe | 0.343 | 0.546 | 0.812 | 0.651 | 417 | 0.004 | 68622 | 8787 | 0.128 |
| JU561 | Europe | 0.786 | 0.450 | 0.899 | 0.646 | 1171 | 0.012 | 92742 | 22392 | 0.241 |
| JU642 | Europe | 0.073 | 0.508 | 0.853 | 0.519 | 929 | 0.009 | 117189 | 14321 | 0.122 |
| JU751 | Europe | 0.211 | 0.284 | 0.697 | 0.366 | 2179 | 0.022 | 149286 | 35978 | 0.241 |
| JU774 | Europe | 0.629 | 0.816 | 0.512 | 0.614 | 1911 | 0.019 | 143568 | 38573 | 0.269 |
| JU775 | Europe | 0.302 | 0.000 | 0.150 | 0.000 | 2625 | 0.026 | 243756 | 54370 | 0.223 |
| JU778 | Europe | 0.567 | 0.000 | 0.763 | 0.399 | 1916 | 0.019 | 143528 | 33372 | 0.233 |
| JU782 | Europe | 0.477 | 0.000 | 0.484 | 0.259 | 2820 | 0.028 | 209429 | 61273 | 0.293 |
| JU792 | Europe | 0.506 | 0.000 | 0.740 | 0.000 | 880 | 0.009 | 138559 | 15865 | 0.114 |
| JU830 | Europe | 0.077 | 0.544 | 0.792 | 0.366 | 2047 | 0.020 | 124853 | 28459 | 0.228 |
| JU847 | Europe | 0.821 | 0.126 | 0.774 | 0.111 | 1524 | 0.015 | 119085 | 24100 | 0.202 |
| KR314 | N. America | 0.457 | 0.696 | 0.894 | 0.862 | 1983 | 0.020 | 131018 | 25048 | 0.191 |
| LKC34 | Africa | 0.868 | 0.193 | 0.745 | 0.421 | 1248 | 0.012 | 123406 | 19662 | 0.159 |
| MY1 | Europe | 0.216 | 0.787 | 0.785 | 0.346 | 1387 | 0.014 | 138209 | 29190 | 0.211 |
| MY10 | Europe | 0.652 | 0.368 | 0.607 | 0.799 | 2475 | 0.025 | 174925 | 55624 | 0.318 |
| MY16 | Europe | 0.260 | 0.000 | 0.245 | 0.000 | 2315 | 0.023 | 183670 | 49123 | 0.267 |
| MY18 | Europe | 0.000 | 0.404 | 0.769 | 0.000 | 1640 | 0.016 | 154521 | 30222 | 0.196 |
| MY2147 | Europe | 0.151 | 0.000 | 0.754 | 0.133 | 860 | 0.009 | 108390 | 15509 | 0.143 |
| MY2212 | Europe | 0.391 | 0.466 | 0.852 | 0.455 | 1169 | 0.012 | 113204 | 16609 | 0.147 |

| | | | | | | | | | | |
|---|---|---|---|---|---|---|---|---|---|---|
| MY23 | Europe | 0.000 | 0.000 | 0.000 | 0.000 | 3822 | 0.038 | 235686 | 70646 | 0.300 |
| MY2453 | Europe | 0.394 | 0.000 | 0.866 | 0.149 | 1280 | 0.013 | 120121 | 23257 | 0.194 |
| MY2530 | Europe | 0.638 | 0.769 | 0.717 | 0.817 | 677 | 0.007 | 107038 | 13179 | 0.123 |
| MY2535 | Europe | 0.316 | 0.935 | 0.853 | 0.678 | 866 | 0.009 | 91689 | 11946 | 0.130 |
| MY2573 | Europe | 0.333 | 0.085 | 0.739 | 0.061 | 488 | 0.005 | 88791 | 5947 | 0.067 |
| MY2585 | Europe | 0.304 | 0.929 | 0.792 | 0.821 | 1612 | 0.016 | 118229 | 27078 | 0.229 |
| MY2693 | Europe | 0.141 | 0.727 | 0.897 | 0.755 | 1569 | 0.016 | 113210 | 27071 | 0.239 |
| MY2713 | Europe | 0.000 | 0.985 | 0.898 | 0.755 | 1779 | 0.018 | 119161 | 30390 | 0.255 |
| MY2741 | Europe | 0.141 | 0.523 | 0.768 | 0.370 | 2204 | 0.022 | 154168 | 40246 | 0.261 |
| MY518 | Europe | 0.318 | 0.562 | 0.953 | 0.170 | 1567 | 0.016 | 119578 | 21311 | 0.178 |
| MY679 | Europe | 0.395 | 0.914 | 0.805 | 0.839 | 1296 | 0.013 | 111818 | 19692 | 0.176 |
| MY772 | Europe | 0.257 | 0.219 | 0.779 | 0.000 | 1250 | 0.012 | 124005 | 17821 | 0.144 |
| MY795 | Europe | 0.260 | 0.523 | 0.768 | 0.000 | 1570 | 0.016 | 129122 | 27025 | 0.209 |
| MY920 | Europe | 0.638 | 0.756 | 0.717 | 0.707 | 943 | 0.009 | 112924 | 17326 | 0.153 |
| N2 | Europe | 0.376 | 1.000 | 0.890 | 0.257 | NA | NA | NA | NA | NA |
| NIC1 | Europe | 0.292 | 0.910 | 0.711 | 0.641 | 1318 | 0.013 | 99391 | 20769 | 0.209 |
| NIC1049 | Europe | 0.286 | 0.487 | 0.739 | 0.061 | 358 | 0.004 | 85411 | 5406 | 0.063 |
| NIC1107 | Europe | 0.292 | 0.000 | 0.713 | 0.134 | 381 | 0.004 | 63682 | 8695 | 0.137 |
| NIC1119 | S. America | 0.816 | 0.783 | 0.589 | 0.662 | 1869 | 0.019 | 152106 | 37905 | 0.249 |
| NIC166 | Europe | 0.073 | 0.622 | 0.723 | 0.269 | 1115 | 0.011 | 105698 | 15695 | 0.148 |
| NIC195 | Atlantic | 0.000 | 0.164 | 0.062 | 0.421 | 3307 | 0.033 | 199802 | 62556 | 0.313 |
| NIC199 | Atlantic | 0.307 | 0.000 | 0.141 | 0.000 | 2923 | 0.029 | 210803 | 56680 | 0.269 |
| NIC2 | Europe | 0.292 | 1.000 | 0.820 | 0.978 | 163 | 0.002 | 31341 | 2270 | 0.072 |
| NIC207 | Europe | 0.292 | 0.599 | 0.865 | 0.988 | 532 | 0.005 | 73533 | 7154 | 0.097 |
| NIC231 | Europe | 0.792 | 0.252 | 0.865 | 0.823 | 1140 | 0.011 | 101135 | 13884 | 0.137 |
| NIC236 | Europe | 0.236 | 0.287 | 0.673 | 0.399 | 1689 | 0.017 | 121592 | 23549 | 0.194 |
| NIC242 | Europe | 0.312 | 0.602 | 0.756 | 0.314 | 1191 | 0.012 | 87395 | 18871 | 0.216 |
| NIC251 | Atlantic | 0.000 | 0.000 | 0.000 | 0.000 | 3802 | 0.038 | 278599 | 77406 | 0.278 |
| NIC252 | Atlantic | 0.000 | 0.382 | 0.274 | 0.220 | 2431 | 0.024 | 192408 | 48136 | 0.250 |
| NIC255 | Europe | 0.292 | 0.652 | 0.862 | 0.979 | 1067 | 0.011 | 77276 | 13333 | 0.173 |
| NIC256 | Europe | 0.412 | 0.000 | 0.559 | 0.259 | 3151 | 0.031 | 182367 | 50546 | 0.277 |
| NIC258 | Europe | 0.398 | 0.234 | 0.058 | 0.000 | 3884 | 0.039 | 236759 | 69948 | 0.295 |
| NIC259 | Europe | 0.280 | 0.000 | 0.739 | 0.424 | 1120 | 0.011 | 129530 | 17763 | 0.137 |
| NIC260 | Europe | 0.302 | 0.636 | 0.650 | 0.078 | 2600 | 0.026 | 192297 | 49858 | 0.259 |
| NIC261 | Europe | 0.477 | 0.000 | 0.484 | 0.000 | 2914 | 0.029 | 212854 | 49906 | 0.234 |
| NIC262 | Europe | 0.434 | 0.000 | 0.484 | 0.074 | 3075 | 0.031 | 207470 | 54754 | 0.264 |
| NIC265 | Europe | 0.256 | 0.000 | 0.085 | 0.000 | 2901 | 0.029 | 250079 | 54501 | 0.218 |

| | | | | | | | | | | |
|---|---|---|---|---|---|---|---|---|---|---|
| NIC266 | Europe | 0.194 | 0.164 | 0.218 | 0.165 | 2197 | 0.022 | 166172 | 41867 | 0.252 |
| NIC267 | Europe | 0.073 | 0.714 | 0.868 | 0.275 | 1107 | 0.011 | 115423 | 15514 | 0.134 |
| NIC268 | Europe | 0.000 | 0.467 | 0.226 | 0.279 | 2243 | 0.022 | 181716 | 36982 | 0.204 |
| NIC269 | Europe | 0.397 | 0.000 | 0.339 | 0.069 | 2088 | 0.021 | 165440 | 31258 | 0.189 |
| NIC271 | Europe | 0.477 | 0.000 | 0.484 | 0.259 | 3178 | 0.032 | 193468 | 54014 | 0.279 |
| NIC272 | Europe | 0.264 | 0.000 | 0.758 | 0.000 | 1766 | 0.018 | 155743 | 29770 | 0.191 |
| NIC274 | Europe | 0.685 | 0.313 | 0.658 | 0.096 | 1339 | 0.013 | 139239 | 26480 | 0.190 |
| NIC275 | Europe | 0.840 | 0.522 | 0.847 | 0.616 | 2548 | 0.025 | 142895 | 40005 | 0.280 |
| NIC276 | Europe | 0.308 | 1.000 | 0.868 | 0.000 | 1249 | 0.012 | 109724 | 18135 | 0.165 |
| NIC277 | Europe | 0.410 | 0.624 | 0.953 | 0.134 | 791 | 0.008 | 104923 | 12374 | 0.118 |
| NIC3 | Europe | 0.605 | 0.602 | 0.817 | 0.124 | 1194 | 0.012 | 138590 | 18969 | 0.137 |
| NIC501 | Europe | 0.614 | 0.000 | 0.204 | 0.000 | 1938 | 0.019 | 167069 | 39544 | 0.237 |
| NIC511 | Europe | 0.151 | 0.826 | 0.829 | 0.257 | 839 | 0.008 | 113548 | 13223 | 0.116 |
| NIC513 | Europe | 0.184 | 0.370 | 0.695 | 0.355 | 1623 | 0.016 | 151535 | 33284 | 0.220 |
| NIC514 | Europe | 0.329 | 0.292 | 0.794 | 0.424 | 1733 | 0.017 | 141823 | 27931 | 0.197 |
| NIC515 | Europe | 0.081 | 0.685 | 0.395 | 0.706 | 2188 | 0.022 | 158133 | 37847 | 0.239 |
| NIC522 | Europe | 0.621 | 0.093 | 0.805 | 0.465 | 934 | 0.009 | 104875 | 19369 | 0.185 |
| NIC523 | Europe | 0.151 | 0.091 | 0.868 | 0.543 | 1398 | 0.014 | 151910 | 23626 | 0.156 |
| NIC526 | Europe | 0.222 | 0.000 | 0.817 | 0.426 | 1177 | 0.012 | 140427 | 23111 | 0.165 |
| NIC527 | Europe | 0.378 | 0.139 | 0.768 | 0.000 | 1129 | 0.011 | 127670 | 21845 | 0.171 |
| NIC528 | Europe | 0.376 | 0.859 | 0.707 | 0.760 | 1725 | 0.017 | 137874 | 34604 | 0.251 |
| NIC529 | Europe | 0.426 | 0.203 | 0.868 | 0.451 | 924 | 0.009 | 103634 | 14178 | 0.137 |
| PB303 | Unknown | 0.717 | 1.000 | 0.865 | 0.829 | 1447 | 0.014 | 98449 | 25381 | 0.258 |
| PS2025 | N. America | 0.081 | 0.517 | 0.000 | 0.069 | 2425 | 0.024 | 197969 | 48596 | 0.245 |
| PX179 | N. America | 0.171 | 0.791 | 0.904 | 0.991 | 513 | 0.005 | 65022 | 9408 | 0.145 |
| QG2075 | N. America | 0.237 | 0.758 | 0.130 | 0.566 | 2166 | 0.022 | 176964 | 44474 | 0.251 |
| QG2811 | Australia | 0.581 | 0.646 | 0.829 | 0.253 | 1259 | 0.013 | 101304 | 24664 | 0.243 |
| QG2813 | Australia | 0.519 | 0.732 | 0.768 | 0.511 | 1215 | 0.012 | 118225 | 21860 | 0.185 |
| QG2818 | Australia | 1.000 | 1.000 | 0.953 | 1.000 | 550 | 0.005 | 92910 | 9877 | 0.106 |
| QG2823 | Australia | 0.286 | 0.742 | 0.817 | 0.903 | 1354 | 0.014 | 117772 | 19541 | 0.166 |
| QG2824 | Australia | 0.390 | 0.666 | 0.953 | 0.851 | 766 | 0.008 | 112581 | 13457 | 0.120 |
| QG2825 | Australia | 0.384 | 0.671 | 0.817 | 0.828 | 1354 | 0.014 | 109802 | 18777 | 0.171 |
| QG2827 | Australia | 0.560 | 0.376 | 0.367 | 0.565 | 2109 | 0.021 | 176894 | 43889 | 0.248 |
| QG2828 | Australia | 0.560 | 0.376 | 0.391 | 0.565 | 2114 | 0.021 | 175041 | 43826 | 0.250 |
| QG2832 | Australia | 0.673 | 0.418 | 0.226 | 0.391 | 1497 | 0.015 | 146501 | 30344 | 0.207 |
| QG2835 | Australia | 0.677 | 0.418 | 0.374 | 0.783 | 1915 | 0.019 | 151287 | 32585 | 0.215 |
| QG2836 | Australia | 0.723 | 0.764 | 0.422 | 0.429 | 1597 | 0.016 | 134594 | 23837 | 0.177 |

| | | | | | | | | | | |
|---|---|---|---|---|---|---|---|---|---|---|
| QG2837 | Australia | 0.673 | 1.000 | 0.226 | 0.400 | 1792 | 0.018 | 143515 | 35248 | 0.246 |
| QG2838 | Australia | 0.985 | 0.756 | 0.316 | 0.645 | 1932 | 0.019 | 159248 | 39142 | 0.246 |
| QG2841 | Australia | 0.916 | 0.947 | 0.672 | 0.000 | 1302 | 0.013 | 136119 | 23138 | 0.170 |
| QG2843 | Australia | 0.976 | 1.000 | 0.901 | 0.999 | 560 | 0.006 | 92856 | 9613 | 0.104 |
| QG2846 | Australia | 0.248 | 0.320 | 0.000 | 0.209 | 2180 | 0.022 | 179522 | 41735 | 0.232 |
| QG2850 | Australia | 0.652 | 0.794 | 0.573 | 0.959 | 1162 | 0.012 | 123595 | 23978 | 0.194 |
| QG2854 | Australia | 0.593 | 0.701 | 0.298 | 0.849 | 1870 | 0.019 | 161265 | 37637 | 0.233 |
| QG2855 | Australia | 0.593 | 0.657 | 0.251 | 0.849 | 1827 | 0.018 | 166740 | 37714 | 0.226 |
| QG2857 | Australia | 0.580 | 0.675 | 0.465 | 0.849 | 1731 | 0.017 | 154760 | 34344 | 0.222 |
| QG2873 | Australia | 0.698 | 0.822 | 0.216 | 0.936 | 1779 | 0.018 | 155811 | 37394 | 0.240 |
| QG2874 | Australia | 0.698 | 0.791 | 0.216 | 0.936 | 1848 | 0.018 | 157752 | 38555 | 0.244 |
| QG2875 | Australia | 0.647 | 0.780 | 0.569 | 0.186 | 1739 | 0.017 | 147350 | 31179 | 0.212 |
| QG2877 | Australia | 0.615 | 0.619 | 0.369 | 0.786 | 2033 | 0.020 | 157354 | 38180 | 0.243 |
| QG2932 | Australia | 0.529 | 0.858 | 0.724 | 0.998 | 1086 | 0.011 | 125449 | 20320 | 0.162 |
| QG536 | N. America | 0.846 | 1.000 | 0.745 | 0.301 | 1712 | 0.017 | 122846 | 36058 | 0.294 |
| QG556 | N. America | 0.000 | 0.512 | 0.088 | 0.000 | 2783 | 0.028 | 168531 | 41279 | 0.245 |
| QG557 | N. America | 0.222 | 0.404 | 0.892 | 0.076 | 1876 | 0.019 | 123964 | 19839 | 0.160 |
| QW947 | S. America | 0.799 | 0.260 | 0.606 | 0.481 | 1740 | 0.017 | 156656 | 33634 | 0.215 |
| QX1211 | N. America | 0.000 | 0.000 | 0.000 | 0.000 | 5240 | 0.052 | 488149 | 123859 | 0.254 |
| QX1212 | N. America | 0.222 | 0.743 | 0.288 | 0.312 | 2032 | 0.020 | 163520 | 39953 | 0.244 |
| QX1233 | N. America | 0.141 | 0.381 | 0.601 | 0.222 | 1543 | 0.015 | 140996 | 21967 | 0.156 |
| QX1791 | Hawaii | 0.000 | 0.000 | 0.000 | 0.000 | 3694 | 0.037 | 263167 | 92669 | 0.352 |
| QX1792 | Hawaii | 0.000 | 0.000 | 0.479 | 0.289 | 2129 | 0.021 | 202531 | 39836 | 0.197 |
| QX1793 | Hawaii | 0.000 | 0.000 | 0.053 | 0.000 | 2178 | 0.022 | 214878 | 42961 | 0.200 |
| QX1794 | Hawaii | 0.000 | 0.000 | 0.000 | 0.000 | 2988 | 0.030 | 227050 | 57989 | 0.255 |
| RC301 | Europe | 0.231 | 1.000 | 0.865 | 0.829 | 1480 | 0.015 | 105523 | 25921 | 0.246 |
| WN2001 | Europe | 0.000 | 0.502 | 0.368 | 0.296 | 2487 | 0.025 | 188044 | 46097 | 0.245 |
| WN2002 | Europe | 0.289 | 0.743 | 0.698 | 0.427 | 1565 | 0.016 | 114573 | 32070 | 0.280 |
| WN2033 | Europe | 0.374 | 0.507 | 0.904 | 0.365 | 1696 | 0.017 | 132699 | 23982 | 0.181 |
| WN2050 | Europe | 0.292 | 0.000 | 0.000 | 0.134 | 1617 | 0.016 | 133276 | 35214 | 0.264 |
| WN2063 | Europe | 0.000 | 0.330 | 0.825 | 0.341 | 1079 | 0.011 | 120087 | 17700 | 0.147 |
| WN2064 | Europe | 0.339 | 0.857 | 0.903 | 0.061 | 439 | 0.004 | 68962 | 5196 | 0.075 |
| WN2066 | Europe | 0.320 | 0.618 | 0.640 | 0.076 | 752 | 0.007 | 81894 | 17704 | 0.216 |
| XZ1513 | Hawaii | 0.000 | 0.000 | 0.000 | 0.000 | 2623 | 0.026 | 235948 | 54106 | 0.229 |
| XZ1514 | Hawaii | 0.000 | 0.000 | 0.000 | 0.000 | 6603 | 0.066 | 497062 | 151663 | 0.305 |
| XZ1515 | Hawaii | 0.000 | 0.000 | 0.224 | 0.000 | 2253 | 0.022 | 234705 | 42956 | 0.183 |
| XZ1516 | Hawaii | 0.000 | 0.000 | 0.000 | 0.000 | 11673 | 0.116 | 676100 | 322172 | 0.477 |

| | | | | | | | | | | |
|---|---|---|---|---|---|---|---|---|---|---|
| XZ1672 | N. America | 0.171 | 0.520 | 0.904 | 0.558 | 846 | 0.008 | 100578 | 12368 | 0.123 |
| XZ1734 | S. America | 0.605 | 0.292 | 0.521 | 0.508 | 2120 | 0.021 | 154270 | 34769 | 0.225 |
| XZ1735 | S. America | 0.362 | 0.425 | 0.432 | 0.457 | 1592 | 0.016 | 140656 | 29981 | 0.213 |
| XZ1756 | N. America | 0.383 | 0.443 | 0.749 | 0.213 | 1792 | 0.018 | 125407 | 29700 | 0.237 |
| XZ2018 | Unknown | 0.096 | 0.826 | 0.904 | 0.999 | 834 | 0.008 | 74749 | 11020 | 0.147 |
| XZ2019 | N. America | 0.000 | 0.000 | 0.000 | 0.000 | 2717 | 0.027 | 235033 | 47753 | 0.203 |
| XZ2020 | N. America | 0.096 | 0.752 | 0.248 | 0.388 | 1051 | 0.010 | 120434 | 22352 | 0.186 |

**Supplementary Table 4 | Genome assembly and annotations metrics**

Genome assembly and annotations metrics shown for the N2 reference genome, the CB4856 long-read genome, and 14 wild isotype genomes generated here. BUSCO (version 4.0.6) completeness scores, using the nematoda_ob10 dataset, are shown for the genome (using the option *-m* genome) and gene set (using the option *-m* proteins) separately. Gene set completeness scores were calculated using only the longest isoform of each protein coding gene. Note: the CB4856 genome was scaffolded based on alignments to the N2 reference genome. To avoid accidentally introducing errors, particularly in hyper-divergent regions where nucleotide identity is low and alignment is difficult, we opted not to use the same approach.

| Isotype | Span (Mb) | Number of contigs | Number of contigs >50kb | Contig N50 length (Mb) | Contig N50 number | Longest contig (Mb) | Span of Ns (kb) | BUSCO genome complete (%) / fragmented (%) | Number of predicted genes | BUSCO gene set complete (%) / fragmented (%) |
|---|---|---|---|---|---|---|---|---|---|---|
| **N2 (reference)** | 100.29 | 7 | 6 | 17.49 | 3 | 20.92 | 0.00 | 99.4 / 0.1 | 20,190 | 99.4 / 0.3 |
| **CB4856 (Kim *et al.* 2019)** | 102.91 | 7 | 6 | 17.99 | 3 | 21.39 | 69.00 | 99.3 / 0.1 | 22,229 | 96.4 / 2.2 |
| **DL238** | 103.40 | 118 | 65 | 2.69 | 13 | 6.28 | 0.00 | 99.4 / 0.2 | 23,177 | 96.2 / 2.4 |
| **ECA36** | 105.62 | 82 | 63 | 2.84 | 12 | 8.75 | 0.00 | 99.5 / 0.2 | 23,610 | 96.3 / 2.4 |
| **ECA396** | 103.36 | 128 | 72 | 2.64 | 15 | 4.60 | 0.00 | 99.5 / 0.2 | 23,227 | 96.2 / 2.3 |
| **EG4725** | 103.85 | 80 | 48 | 3.61 | 11 | 6.26 | 0.00 | 99.6 / 0.1 | 23,178 | 96.4 / 2.2 |
| **JU1400** | 103.28 | 67 | 53 | 2.88 | 13 | 5.71 | 0.00 | 99.3 / 0.2 | 23,229 | 96.0 / 2.2 |
| **JU2526** | 102.93 | 98 | 78 | 2.07 | 16 | 5.05 | 0.00 | 99.5 / 0.1 | 23,140 | 95.9 / 2.6 |
| **JU2600** | 104.21 | 87 | 56 | 3.24 | 13 | 5.77 | 0.00 | 99.6 / 0.1 | 23,216 | 96.3 / 2.4 |
| **JU310** | 114.26 | 283 | 139 | 2.68 | 15 | 6.22 | 0.00 | 99.6 / 0.1 | 25,821 | 96.2 / 2.3 |
| **MY2147** | 103.78 | 74 | 57 | 3.67 | 12 | 5.80 | 0.00 | 99.6 / 0.1 | 23,164 | 96.4 / 2.3 |
| **MY2693** | 102.92 | 113 | 59 | 2.86 | 13 | 6.21 | 0.00 | 99.6 / 0.1 | 23,203 | 96.1 / 2.5 |
| **NIC2** | 103.06 | 89 | 67 | 2.80 | 13 | 6.79 | 0.00 | 99.7 / 0.1 | 23,171 | 96.2 / 2.5 |
| **NIC526** | 104.20 | 85 | 58 | 2.81 | 13 | 6.78 | 0.00 | 99.6 / 0.1 | 23,387 | 96.1 / 2.6 |
| **QX1794** | 108.69 | 185 | 107 | 2.49 | 14 | 6.50 | 0.00 | 99.5 / 0.1 | 24,207 | 96.1 / 2.2 |
| **XZ1516** | 105.79 | 149 | 89 | 2.05 | 19 | 5.11 | 0.00 | 99.5 / 0.2 | 23,791 | 96.1 / 2.5 |

**Supplementary Table 5 | Distribution of the species-wide hyper-divergent regions in *C. elegans***
The genomic coordinates of all hyper-divergent regions described in the manuscript. We calculated the frequency each 1 kb genomic bin was classified as hyper-divergent across the *C. elegans* species. For each hyper-divergent region, we report the average 1 kb bin frequency.

| Chromosome | Start | End | Size (bp) | Frequency |
|---|---|---|---|---|
| I | 863000 | 875000 | 12000 | 0.08663618 |
| I | 1018000 | 1063000 | 45000 | 0.12445799 |
| I | 1101000 | 1127000 | 26000 | 0.07422608 |
| I | 1169000 | 1179000 | 10000 | 0.01646341 |
| I | 1181000 | 1224000 | 43000 | 0.12712706 |
| I | 1337000 | 1425000 | 88000 | 0.13639828 |
| I | 1513000 | 1529000 | 16000 | 0.16215701 |
| I | 1563000 | 1587000 | 24000 | 0.01575203 |
| I | 1600000 | 1628000 | 28000 | 0.09037456 |
| I | 1749000 | 1772000 | 23000 | 0.10418876 |
| I | 1864000 | 1873000 | 9000 | 0.10060976 |
| I | 1895000 | 1938000 | 43000 | 0.17129892 |
| I | 1949000 | 1967000 | 18000 | 0.37466125 |
| I | 2044000 | 2055000 | 11000 | 0.04296009 |
| I | 2157000 | 2170000 | 13000 | 0.14798311 |
| I | 2328000 | 2363000 | 35000 | 0.41010453 |
| I | 2467000 | 2486000 | 19000 | 0.10943517 |
| I | 2799000 | 2809000 | 10000 | 0.01829268 |
| I | 3019000 | 3034000 | 15000 | 0.0550813 |
| I | 3060000 | 3073000 | 13000 | 0.06214822 |
| I | 3183000 | 3195000 | 12000 | 0.10670732 |
| I | 3229000 | 3241000 | 12000 | 0.20147358 |
| I | 3586000 | 3598000 | 12000 | 0.00304878 |
| I | 3799000 | 3809000 | 10000 | 0.00304878 |
| I | 4453000 | 4463000 | 10000 | 0.00426829 |
| I | 10306000 | 10315000 | 9000 | 0.00914634 |
| I | 10906000 | 10945000 | 39000 | 0.01657286 |
| I | 10953000 | 11010000 | 57000 | 0.03712024 |
| I | 11432000 | 11456000 | 24000 | 0.07050305 |
| I | 11468000 | 11482000 | 14000 | 0.02482578 |
| I | 11833000 | 11842000 | 9000 | 0.01490515 |
| I | 11915000 | 11925000 | 10000 | 0.14756098 |
| I | 12042000 | 12057000 | 15000 | 0.03780488 |

| | | | | |
|---|---|---|---|---|
| I | 12061000 | 12158000 | 97000 | 0.06389867 |
| I | 12206000 | 12216000 | 10000 | 0.00304878 |
| I | 12220000 | 12475000 | 255000 | 0.01846007 |
| I | 12606000 | 12682000 | 76000 | 0.05267169 |
| I | 12688000 | 12698000 | 10000 | 0.02042683 |
| I | 12751000 | 12763000 | 12000 | 0.21620935 |
| I | 12785000 | 12809000 | 24000 | 0.01816565 |
| I | 12986000 | 12995000 | 9000 | 0.00304878 |
| I | 13012000 | 13226000 | 214000 | 0.04628733 |
| I | 13341000 | 13368000 | 27000 | 0.03331075 |
| I | 13375000 | 13386000 | 11000 | 0.04711752 |
| I | 13799000 | 13820000 | 21000 | 0.0203252 |
| I | 14027000 | 14037000 | 10000 | 0.01402439 |
| I | 14281000 | 14319000 | 38000 | 0.01131258 |
| I | 14410000 | 14425000 | 15000 | 0.09898374 |
| I | 14491000 | 14500000 | 9000 | 0.0196477 |
| I | 14503000 | 14518000 | 15000 | 0.10386179 |
| I | 14535000 | 14554000 | 19000 | 0.01957638 |
| II | 262000 | 332000 | 70000 | 0.0527439 |
| II | 403000 | 424000 | 21000 | 0.00391986 |
| II | 429000 | 457000 | 28000 | 0.03059669 |
| II | 464000 | 629000 | 165000 | 0.0313932 |
| II | 631000 | 682000 | 51000 | 0.03186275 |
| II | 868000 | 994000 | 126000 | 0.06571816 |
| II | 1001000 | 1076000 | 75000 | 0.06617886 |
| II | 1108000 | 1117000 | 9000 | 0.01219512 |
| II | 1256000 | 1310000 | 54000 | 0.16327913 |
| II | 1353000 | 1362000 | 9000 | 0.06402439 |
| II | 1369000 | 1439000 | 70000 | 0.01019164 |
| II | 1515000 | 1700000 | 185000 | 0.22343441 |
| II | 1703000 | 2194000 | 491000 | 0.32531295 |
| II | 2219000 | 2509000 | 290000 | 0.40447855 |
| II | 2528000 | 2654000 | 126000 | 0.07108982 |
| II | 2715000 | 2724000 | 9000 | 0.00304878 |
| II | 2785000 | 2798000 | 13000 | 0.01242964 |
| II | 2799000 | 2810000 | 11000 | 0.0058204 |
| II | 2835000 | 2845000 | 10000 | 0.33414634 |

| | | | | |
|---|---|---|---|---|
| II | 2870000 | 2925000 | 55000 | 0.16385809 |
| II | 2965000 | 2974000 | 9000 | 0.00813008 |
| II | 3060000 | 3087000 | 27000 | 0.26343722 |
| II | 3177000 | 3271000 | 94000 | 0.52075765 |
| II | 3280000 | 3399000 | 119000 | 0.3860166 |
| II | 3499000 | 3512000 | 13000 | 0.04620075 |
| II | 3660000 | 3780000 | 120000 | 0.34751016 |
| II | 3800000 | 3887000 | 87000 | 0.09006168 |
| II | 3902000 | 3926000 | 24000 | 0.00304878 |
| II | 3942000 | 3954000 | 12000 | 0.00304878 |
| II | 3987000 | 4013000 | 26000 | 0.03470919 |
| II | 4174000 | 4186000 | 12000 | 0.00304878 |
| II | 4197000 | 4208000 | 11000 | 0.01801552 |
| II | 4303000 | 4312000 | 9000 | 0.0152439 |
| II | 4432000 | 4444000 | 12000 | 0.02769309 |
| II | 4720000 | 4731000 | 11000 | 0.05792683 |
| II | 5295000 | 5315000 | 20000 | 0.00304878 |
| II | 5839000 | 5869000 | 30000 | 0.00304878 |
| II | 10429000 | 10465000 | 36000 | 0.03353659 |
| II | 10626000 | 10674000 | 48000 | 0.01587907 |
| II | 11367000 | 11400000 | 33000 | 0.00554324 |
| II | 11980000 | 11994000 | 14000 | 0.11672474 |
| II | 12012000 | 12022000 | 10000 | 0.08323171 |
| II | 12028000 | 12038000 | 10000 | 0.10518293 |
| II | 12061000 | 12070000 | 9000 | 0.0304878 |
| II | 12083000 | 12108000 | 25000 | 0.04158537 |
| II | 12289000 | 12302000 | 13000 | 0.00938086 |
| II | 12397000 | 12407000 | 10000 | 0.00823171 |
| II | 12454000 | 12622000 | 168000 | 0.06854312 |
| II | 12623000 | 12653000 | 30000 | 0.00558943 |
| II | 12659000 | 12717000 | 58000 | 0.04331371 |
| II | 12718000 | 12732000 | 14000 | 0.01263066 |
| II | 12738000 | 12748000 | 10000 | 0.01310976 |
| II | 12773000 | 12786000 | 13000 | 0.00304878 |
| II | 12798000 | 12819000 | 21000 | 0.02642276 |
| II | 12867000 | 12881000 | 14000 | 0.07796167 |
| II | 12892000 | 12901000 | 9000 | 0.00304878 |

| | | | | |
|---|---|---|---|---|
| II | 12928000 | 12938000 | 10000 | 0.00304878 |
| II | 13120000 | 13154000 | 34000 | 0.00887733 |
| II | 13199000 | 13223000 | 24000 | 0.02566057 |
| II | 13244000 | 13265000 | 21000 | 0.12166086 |
| II | 13320000 | 13341000 | 21000 | 0.11541812 |
| II | 13354000 | 13415000 | 61000 | 0.20186925 |
| II | 13430000 | 13467000 | 37000 | 0.08833223 |
| II | 13472000 | 13496000 | 24000 | 0.00635163 |
| II | 13520000 | 13579000 | 59000 | 0.16577098 |
| II | 13580000 | 13590000 | 10000 | 0.00457317 |
| II | 13612000 | 13630000 | 18000 | 0.03895664 |
| II | 13642000 | 13679000 | 37000 | 0.06056361 |
| II | 13736000 | 13763000 | 27000 | 0.05770099 |
| II | 13771000 | 13780000 | 9000 | 0.00304878 |
| II | 13843000 | 13852000 | 9000 | 0.0304878 |
| II | 13862000 | 13886000 | 24000 | 0.14278455 |
| II | 13889000 | 13898000 | 9000 | 0.02134146 |
| II | 13936000 | 13957000 | 21000 | 0.02366434 |
| II | 13967000 | 13978000 | 11000 | 0.01108647 |
| II | 14076000 | 14090000 | 14000 | 0.07012195 |
| II | 14224000 | 14236000 | 12000 | 0.04674797 |
| II | 14303000 | 14338000 | 35000 | 0.08057491 |
| II | 14979000 | 14988000 | 9000 | 0.04234417 |
| III | 0 | 345000 | 345000 | 0.12352421 |
| III | 697000 | 708000 | 11000 | 0.66407982 |
| III | 920000 | 934000 | 14000 | 0.01067073 |
| III | 936000 | 1098000 | 162000 | 0.33278756 |
| III | 1117000 | 1140000 | 23000 | 0.03446448 |
| III | 1172000 | 1437000 | 265000 | 0.28515877 |
| III | 1465000 | 1549000 | 84000 | 0.06435105 |
| III | 1559000 | 1574000 | 15000 | 0.01869919 |
| III | 1578000 | 1619000 | 41000 | 0.06685009 |
| III | 1625000 | 1639000 | 14000 | 0.00979965 |
| III | 1666000 | 1679000 | 13000 | 0.01313321 |
| III | 1682000 | 1706000 | 24000 | 0.03125 |
| III | 1722000 | 1732000 | 10000 | 0.01737805 |
| III | 1746000 | 1799000 | 53000 | 0.04780258 |

| | | | | |
|---|---|---|---|---|
| III | 1827000 | 1842000 | 15000 | 0.10081301 |
| III | 1843000 | 1859000 | 16000 | 0.01333841 |
| III | 1861000 | 1870000 | 9000 | 0.03387534 |
| III | 1899000 | 1917000 | 18000 | 0.10162602 |
| III | 1984000 | 2.00E+06 | 16000 | 0.07602896 |
| III | 2037000 | 2101000 | 64000 | 0.02372332 |
| III | 2199000 | 2358000 | 159000 | 0.11058061 |
| III | 2369000 | 2445000 | 76000 | 0.11501123 |
| III | 2512000 | 2539000 | 27000 | 0.07983288 |
| III | 2567000 | 2576000 | 9000 | 0.09146341 |
| III | 2589000 | 2613000 | 24000 | 0.15612297 |
| III | 2687000 | 2696000 | 9000 | 0.09044715 |
| III | 2763000 | 2778000 | 15000 | 0.02947154 |
| III | 2788000 | 2803000 | 15000 | 0.04654472 |
| III | 2815000 | 2838000 | 23000 | 0.0292948 |
| III | 2901000 | 2917000 | 16000 | 0.00304878 |
| III | 2959000 | 3012000 | 53000 | 0.00719052 |
| III | 3069000 | 3089000 | 20000 | 0.10487805 |
| III | 3289000 | 3334000 | 45000 | 0.20406504 |
| III | 7669000 | 7678000 | 9000 | 0.07621951 |
| III | 9912000 | 9968000 | 56000 | 0.09097343 |
| III | 11037000 | 11050000 | 13000 | 0.065197 |
| III | 11093000 | 11143000 | 50000 | 0.06530488 |
| III | 11152000 | 11169000 | 17000 | 0.09379484 |
| III | 11273000 | 11283000 | 10000 | 0.21128049 |
| III | 11567000 | 11593000 | 26000 | 0.03764071 |
| III | 11922000 | 11931000 | 9000 | 0.00914634 |
| III | 12035000 | 12050000 | 15000 | 0.00304878 |
| III | 12062000 | 12074000 | 12000 | 0.13135163 |
| III | 12232000 | 12245000 | 13000 | 0.02110694 |
| III | 12251000 | 12260000 | 9000 | 0.00914634 |
| III | 12296000 | 12385000 | 89000 | 0.22163606 |
| III | 12409000 | 12423000 | 14000 | 0.05052265 |
| III | 12425000 | 12451000 | 26000 | 0.00422139 |
| III | 12526000 | 12535000 | 9000 | 0.04742547 |
| III | 12598000 | 12608000 | 10000 | 0.02439024 |
| III | 12673000 | 12683000 | 10000 | 0.14420732 |

| | | | | |
|---|---|---|---|---|
| III | 12729000 | 12760000 | 31000 | 0.00629426 |
| III | 12824000 | 12836000 | 12000 | 0.00304878 |
| III | 12992000 | 13002000 | 10000 | 0.00579268 |
| III | 13231000 | 13243000 | 12000 | 0.00304878 |
| IV | 16000 | 32000 | 16000 | 0.05945122 |
| IV | 1005000 | 1022000 | 17000 | 0.06258967 |
| IV | 1074000 | 1098000 | 24000 | 0.15866362 |
| IV | 1113000 | 1125000 | 12000 | 0.03455285 |
| IV | 1228000 | 1352000 | 124000 | 0.08448072 |
| IV | 1536000 | 1551000 | 15000 | 0.0050813 |
| IV | 1590000 | 1609000 | 19000 | 0.05327343 |
| IV | 1816000 | 1888000 | 72000 | 0.00821477 |
| IV | 1983000 | 2034000 | 51000 | 0.13259206 |
| IV | 2054000 | 2067000 | 13000 | 0.005394 |
| IV | 2105000 | 2116000 | 11000 | 0.00498891 |
| IV | 2184000 | 2210000 | 26000 | 0.09111163 |
| IV | 2264000 | 2275000 | 11000 | 0.18902439 |
| IV | 2335000 | 2366000 | 31000 | 0.10188828 |
| IV | 2464000 | 2481000 | 17000 | 0.01058106 |
| IV | 2487000 | 2507000 | 20000 | 0.01128049 |
| IV | 2519000 | 2533000 | 14000 | 0.05422474 |
| IV | 2552000 | 2603000 | 51000 | 0.43896461 |
| IV | 2685000 | 2746000 | 61000 | 0.32521991 |
| IV | 2759000 | 2770000 | 11000 | 0.02549889 |
| IV | 2781000 | 2800000 | 19000 | 0.01941592 |
| IV | 2809000 | 2829000 | 20000 | 0.03612805 |
| IV | 2831000 | 2843000 | 12000 | 0.00304878 |
| IV | 2859000 | 2891000 | 32000 | 0.2105564 |
| IV | 2949000 | 2960000 | 11000 | 0.01274945 |
| IV | 3369000 | 3383000 | 14000 | 0.07142857 |
| IV | 3455000 | 3480000 | 25000 | 0.00304878 |
| IV | 3506000 | 3517000 | 11000 | 0.01773836 |
| IV | 3684000 | 3694000 | 10000 | 0.15640244 |
| IV | 3732000 | 3744000 | 12000 | 0.00813008 |
| IV | 3810000 | 3822000 | 12000 | 0.00304878 |
| IV | 3823000 | 3833000 | 10000 | 0.05487805 |
| IV | 3887000 | 3973000 | 86000 | 0.07111458 |

| | | | | |
|---|---|---|---|---|
| IV | 4052000 | 4068000 | 16000 | 0.02477134 |
| IV | 4098000 | 4171000 | 73000 | 0.39308386 |
| IV | 4347000 | 4358000 | 11000 | 0.11419069 |
| IV | 4370000 | 4380000 | 10000 | 0.04481707 |
| IV | 4407000 | 4416000 | 9000 | 0.05792683 |
| IV | 5447000 | 5465000 | 18000 | 0.02828591 |
| IV | 5821000 | 5843000 | 22000 | 0.02757761 |
| IV | 6206000 | 6302000 | 96000 | 0.0593877 |
| IV | 6606000 | 6626000 | 20000 | 0.05304878 |
| IV | 6629000 | 6661000 | 32000 | 0.03544207 |
| IV | 7451000 | 7471000 | 20000 | 0.11402439 |
| IV | 12376000 | 12446000 | 70000 | 0.00688153 |
| IV | 12827000 | 12902000 | 75000 | 0.03495935 |
| IV | 12947000 | 12956000 | 9000 | 0.00304878 |
| IV | 12987000 | 12996000 | 9000 | 0.02743902 |
| IV | 13416000 | 13428000 | 12000 | 0.12754065 |
| IV | 13517000 | 13543000 | 26000 | 0.00879456 |
| IV | 13623000 | 13641000 | 18000 | 0.00304878 |
| IV | 13980000 | 14004000 | 24000 | 0.0152439 |
| IV | 14170000 | 14185000 | 15000 | 0.02764228 |
| IV | 14186000 | 14195000 | 9000 | 0.00914634 |
| IV | 14519000 | 14551000 | 32000 | 0.03134527 |
| IV | 15310000 | 15321000 | 11000 | 0.00304878 |
| IV | 15439000 | 15479000 | 40000 | 0.0601372 |
| IV | 15487000 | 15496000 | 9000 | 0.00304878 |
| IV | 15612000 | 15634000 | 22000 | 0.03616962 |
| IV | 15759000 | 15769000 | 10000 | 0.07103659 |
| IV | 15897000 | 15908000 | 11000 | 0.01995565 |
| IV | 15944000 | 15954000 | 10000 | 0.00304878 |
| IV | 16221000 | 16255000 | 34000 | 0.02501793 |
| IV | 16285000 | 16362000 | 77000 | 0.10433164 |
| IV | 16425000 | 16447000 | 22000 | 0.04226718 |
| IV | 16462000 | 16492000 | 30000 | 0.01351626 |
| IV | 16709000 | 16770000 | 61000 | 0.1170032 |
| IV | 16788000 | 16806000 | 18000 | 0.04674797 |
| IV | 16838000 | 16924000 | 86000 | 0.06891662 |
| IV | 17075000 | 17085000 | 10000 | 0.0222561 |

| | | | | |
|---|---|---|---|---|
| IV | 17134000 | 17150000 | 16000 | 0.03429878 |
| IV | 17207000 | 17230000 | 23000 | 0.00596501 |
| IV | 17254000 | 17396000 | 142000 | 0.17264256 |
| IV | 17404000 | 17422000 | 18000 | 0.10687669 |
| IV | 17446000 | 17458000 | 12000 | 0.18267276 |
| V | 10000 | 1141000 | 1131000 | 0.07596612 |
| V | 1486000 | 1541000 | 55000 | 0.05515521 |
| V | 1621000 | 1664000 | 43000 | 0.12344016 |
| V | 1672000 | 1722000 | 50000 | 0.05469512 |
| V | 1844000 | 1853000 | 9000 | 0.00304878 |
| V | 1954000 | 1965000 | 11000 | 0.0058204 |
| V | 2005000 | 2031000 | 26000 | 0.00574578 |
| V | 2037000 | 2066000 | 29000 | 0.25851556 |
| V | 2127000 | 2185000 | 58000 | 0.00846299 |
| V | 2227000 | 2376000 | 149000 | 0.07055165 |
| V | 2549000 | 3195000 | 646000 | 0.05841765 |
| V | 3237000 | 3693000 | 456000 | 0.1346879 |
| V | 3712000 | 3731000 | 19000 | 0.00304878 |
| V | 3758000 | 4333000 | 575000 | 0.05503181 |
| V | 4339000 | 4355000 | 16000 | 0.0390625 |
| V | 4358000 | 4388000 | 30000 | 0.02296748 |
| V | 4404000 | 4418000 | 14000 | 0.00500871 |
| V | 4434000 | 4465000 | 31000 | 0.00304878 |
| V | 4531000 | 4540000 | 9000 | 0.00609756 |
| V | 4599000 | 4615000 | 16000 | 0.00304878 |
| V | 4743000 | 4769000 | 26000 | 0.01512664 |
| V | 4855000 | 4873000 | 18000 | 0.02879404 |
| V | 4920000 | 4957000 | 37000 | 0.00988794 |
| V | 4976000 | 5077000 | 101000 | 0.02460155 |
| V | 5078000 | 5091000 | 13000 | 0.00750469 |
| V | 5177000 | 5270000 | 93000 | 0.0035733 |
| V | 6362000 | 6373000 | 11000 | 0.01829268 |
| V | 6380000 | 6394000 | 14000 | 0.00304878 |
| V | 6416000 | 6427000 | 11000 | 0.00609756 |
| V | 6877000 | 6890000 | 13000 | 0.00304878 |
| V | 7102000 | 7170000 | 68000 | 0.01129842 |
| V | 7209000 | 7669000 | 460000 | 0.06605912 |

| | | | | |
|---|---|---|---|---|
| V | 8741000 | 8775000 | 34000 | 0.01398852 |
| V | 8960000 | 8990000 | 30000 | 0.07042683 |
| V | 10454000 | 10464000 | 10000 | 0.00304878 |
| V | 12362000 | 12455000 | 93000 | 0.04756753 |
| V | 13044000 | 13124000 | 80000 | 0.00499238 |
| V | 13165000 | 13175000 | 10000 | 0.00304878 |
| V | 13500000 | 13511000 | 11000 | 0.01967849 |
| V | 13542000 | 13552000 | 10000 | 0.04512195 |
| V | 13588000 | 13601000 | 13000 | 0.06707317 |
| V | 13676000 | 13767000 | 91000 | 0.00512597 |
| V | 13806000 | 13962000 | 156000 | 0.00603893 |
| V | 14305000 | 14320000 | 15000 | 0.00304878 |
| V | 14501000 | 14662000 | 161000 | 0.04192547 |
| V | 15160000 | 15200000 | 40000 | 0.00914634 |
| V | 15210000 | 16161000 | 951000 | 0.08622503 |
| V | 16175000 | 16452000 | 277000 | 0.11743858 |
| V | 16458000 | 16512000 | 54000 | 0.3023374 |
| V | 16533000 | 16675000 | 142000 | 0.08457145 |
| V | 16693000 | 16781000 | 88000 | 0.06492517 |
| V | 16833000 | 17120000 | 287000 | 0.14553412 |
| V | 17132000 | 17742000 | 610000 | 0.19784086 |
| V | 17743000 | 17888000 | 145000 | 0.12626156 |
| V | 17901000 | 17910000 | 9000 | 0.00304878 |
| V | 17920000 | 18001000 | 81000 | 0.01637308 |
| V | 18048000 | 18074000 | 26000 | 0.3570591 |
| V | 18127000 | 18262000 | 135000 | 0.31935411 |
| V | 18270000 | 18333000 | 63000 | 0.02855207 |
| V | 18335000 | 18383000 | 48000 | 0.02337398 |
| V | 18391000 | 18421000 | 30000 | 0.00955285 |
| V | 18430000 | 19779000 | 1349000 | 0.13560858 |
| V | 19808000 | 19824000 | 16000 | 0.02839177 |
| V | 19834000 | 19874000 | 40000 | 0.09032012 |
| V | 19886000 | 19952000 | 66000 | 0.09774575 |
| V | 19960000 | 19999000 | 39000 | 0.05534709 |
| V | 20060000 | 20087000 | 27000 | 0.32588076 |
| V | 20099000 | 20665000 | 566000 | 0.1189186 |
| X | 1000 | 109000 | 108000 | 0.12347561 |

| | | | | |
|---|---|---|---|---|
| X | 115000 | 292000 | 177000 | 0.10438198 |
| X | 1460000 | 1516000 | 56000 | 0.0341899 |
| X | 1534000 | 1547000 | 13000 | 0.03822702 |
| X | 1558000 | 1596000 | 38000 | 0.00401155 |
| X | 1645000 | 1695000 | 50000 | 0.01384146 |
| X | 1711000 | 1733000 | 22000 | 0.03464523 |
| X | 1755000 | 1776000 | 21000 | 0.15926249 |
| X | 1800000 | 1840000 | 40000 | 0.0242378 |
| X | 2030000 | 2039000 | 9000 | 0.00304878 |
| X | 2744000 | 2753000 | 9000 | 0.00304878 |
| X | 4342000 | 4356000 | 14000 | 0.02743902 |
| X | 5046000 | 5056000 | 10000 | 0.07743902 |
| X | 5114000 | 5123000 | 9000 | 0.00609756 |
| X | 5309000 | 5318000 | 9000 | 0.00304878 |
| X | 6112000 | 6128000 | 16000 | 0.00381098 |
| X | 6905000 | 6917000 | 12000 | 0.06097561 |
| X | 7345000 | 7365000 | 20000 | 0.35015244 |
| X | 7896000 | 7908000 | 12000 | 0.0152439 |
| X | 7914000 | 7930000 | 16000 | 0.05182927 |
| X | 8101000 | 8110000 | 9000 | 0.03353659 |
| X | 11768000 | 11784000 | 16000 | 0.00304878 |
| X | 11798000 | 11807000 | 9000 | 0.01930894 |
| X | 12201000 | 12224000 | 23000 | 0.02889714 |
| X | 12602000 | 12611000 | 9000 | 0.01422764 |
| X | 12992000 | 13001000 | 9000 | 0.03353659 |
| X | 13451000 | 13466000 | 15000 | 0.02134146 |
| X | 14174000 | 14183000 | 9000 | 0.00304878 |
| X | 14215000 | 14227000 | 12000 | 0.55919715 |
| X | 14355000 | 14370000 | 15000 | 0.03882114 |
| X | 14371000 | 14396000 | 25000 | 0.09792683 |
| X | 14635000 | 14644000 | 9000 | 0.03658537 |
| X | 15533000 | 15543000 | 10000 | 0.0054878 |
| X | 15617000 | 15637000 | 20000 | 0.01082317 |
| X | 15812000 | 15836000 | 24000 | 0.01892785 |
| X | 17260000 | 17269000 | 9000 | 0.00609756 |
| X | 17287000 | 17302000 | 15000 | 0.00914634 |
| X | 17703000 | 17718000 | 15000 | 0.03597561 |

**Supplementary Table 6 | Summary of statistics of hyper-divergent regions**
Mean variant density (per kb), mean coverage (%), and size (Mb) of genomic regions relative to the N2 reference genome (entire chromosome, arm, center, and tip), hyper-divergent regions (HD), and non-divergent regions (ND) are summarized.

| Chr | Statistics | All | All_HD | All_ND | Arm | Arm_HD | Arm_ND | Center | Center_ HD | Center_ ND | Tip | Tip_HD | Tip_N D |
|---|---|---|---|---|---|---|---|---|---|---|---|---|---|
| I | Mean variant density (per kb) | 1.08 | 15.97 | 1.03 | 1.81 | 16.93 | 1.7 | 0.33 | 3.59 | 0.33 | 1.41 | NA | 1.41 |
| | Mean coverage (%) | 99.93 | 43.84 | 100.12 | 80.52 | 44.56 | 80.91 | 109.22 | 34.44 | 109.22 | 199.8 | NA | 199.8 |
| | Size (Mb) | 15.07 | 1.61 | 15.07 | 7.17 | 1.49 | 7.17 | 7.18 | 0.12 | 7.18 | 0.72 | NA | 0.72 |
| II | Mean variant density (per kb) | 2.32 | 18.49 | 1.82 | 3.71 | 18.11 | 2.65 | 1.07 | 24.51 | 1.06 | 1.3 | 20.92 | 1.25 |
| | Mean coverage (%) | 97.82 | 52.22 | 99.69 | 83.71 | 49.36 | 87.68 | 112.73 | 98.36 | 112.73 | 92.34 | 67.71 | 92.36 |
| | Size (Mb) | 15.28 | 3.58 | 15.28 | 7.16 | 3.34 | 7.16 | 7.14 | 0.19 | 7.14 | 0.98 | 0.05 | 0.98 |
| III | Mean variant density (per kb) | 1.76 | 14.97 | 1.53 | 2.84 | 12.99 | 2.47 | 0.8 | 28.48 | 0.78 | 1.57 | 22.2 | 0.8 |
| | Mean coverage (%) | 94.48 | 41.61 | 95.16 | 76.47 | 39.3 | 77.75 | 111.34 | 74.35 | 111.36 | 92.94 | 46.88 | 94.29 |
| | Size (Mb) | 13.78 | 2.16 | 13.78 | 6.11 | 1.74 | 6.11 | 6.62 | 0.06 | 6.62 | 1.06 | 0.36 | 1.06 |
| IV | Mean variant density (per kb) | 1.27 | 17.97 | 1.13 | 2.07 | 13.58 | 1.97 | 0.55 | 26.05 | 0.49 | 1.91 | 20.09 | 1.21 |
| | Mean coverage (%) | 97.63 | 54.83 | 97.88 | 84.04 | 44.06 | 84.41 | 108.19 | 73.54 | 108.29 | 96.43 | 61.6 | 97.08 |
| | Size (Mb) | 17.49 | 2.18 | 17.49 | 6.92 | 1.22 | 6.92 | 9.07 | 0.56 | 9.07 | 1.5 | 0.4 | 1.5 |
| V | Mean variant density (per kb) | 1.97 | 26.47 | 1.12 | 2.8 | 22.41 | 1.75 | 1.15 | 33.83 | 0.6 | 3.05 | 33.56 | 1.04 |
| | Mean coverage (%) | 99.68 | 57.42 | 101.82 | 85.24 | 47.17 | 89.22 | 112 | 79.5 | 112.59 | 99.07 | 65.8 | 101.19 |
| | Size (Mb) | 20.92 | 10.04 | 20.92 | 9.04 | 6.45 | 9.04 | 10.65 | 2.63 | 10.65 | 1.23 | 0.96 | 1.23 |
| X | Mean variant density (per kb) | 1.23 | 8.95 | 1.21 | 1.26 | 12.17 | 1.24 | 1.28 | 15.81 | 1.27 | 0.87 | 1.78 | 0.87 |
| | Mean coverage (%) | 108.95 | 43.55 | 109.01 | 105.61 | 55.17 | 105.68 | 114.83 | 79.16 | 114.83 | 105.99 | 13.8 | 106.18 |
| | Size (Mb) | 17.72 | 0.92 | 17.72 | 9.5 | 0.47 | 9.5 | 6.34 | 0.12 | 6.34 | 1.87 | 0.32 | 1.87 |
