## Supplementary Data Guide for "Balancing selection maintains hyper-divergent haplotypes in *C. elegans*"

**Supplementary Data 1: Distributions of hyper-divergent regions across 327 non-reference *C. elegans* isotypes**

This file contains information for location and size of hyper-divergent regions across 327 non-reference *C. elegans* isotypes

(File type: XLS, size: 1.4 MB)

**Supplementary Data 2: Gene-set enrichment analysis results from the WormCat tools**

This file contains gene-set enrichment analysis (WormCat tool) for hyper-divergent regions.

(File type: XLS, size: 37 kb)

**Supplementary Data 3: Gene-set enrichment analysis results from the Gene Ontology (GO) enrichment analysis**

This file contains gene-set enrichment analysis (GO enrichment analysis) for hyper-divergent regions.

(File type: XLS, size: 57 kb)

**Supplementary Data 4: Functional annotations of genes in the three characterized hyper-divergent regions**

This file contains BLASTP and InterProScan annotations for all haplotype-specific genes in the three characterized hyper-divergent regions.

(File type: XLS, size: 99 kb)
